# Neandertal ancestry through time: Insights from genomes of ancient and present-day humans

**DOI:** 10.1101/2024.05.13.593955

**Authors:** Leonardo N. M. Iasi, Manjusha Chintalapati, Laurits Skov, Alba Bossoms Mesa, Mateja Hajdinjak, Benjamin M. Peter, Priya Moorjani

## Abstract

Gene flow from Neandertals has shaped the landscape of genetic and phenotypic variation in modern humans. We identify the location and size of introgressed Neandertal ancestry segments in more than 300 genomes spanning the last 50,000 years. We study how Neandertal ancestry is shared among individuals to infer the time and duration of the Neandertal gene flow. We find the correlation of Neandertal segment locations across individuals and their divergence to sequenced Neandertals, both support a model of single major Neandertal gene flow. Our catalog of introgressed segments through time confirms that most natural selection–positive and negative–on Neandertal ancestry variants occurred immediately after the gene flow, and provides new insights into how the contact with Neandertals shaped human origins and adaptation.

## Introduction

The sequencing of the Neandertal (*1*–*4*) and Denisovan (*5, 6*) genomes has revealed extensive gene flow between the ancestors of modern humans and archaic hominins. As a result, most non-Africans harbor 1–2% of Neandertal ancestry, with East Asians exhibiting ∼20% more Neandertal ancestry compared to West Eurasians (*6*). This gene flow has been inferred to have occurred between 41,000–54,000 years ago, but it remains debated if there were secondary interactions between Neandertals and early modern humans (e.g., Oase, Bacho Kiro and Ust’-Ishim) or in the ancestors of East Asians or potential dilution in ancestors of West Eurasians from a group without Neandertal ancestry (*7*–*11*). Moreover, previous studies have identified that the distribution of Neandertal ancestry is not uniform across the genome: some regions are significantly depleted of Neanderthal ancestry (referred as “archaic deserts”), while other regions contain variants at unusually high frequency possibly because they harbor beneficial mutations (“candidates of adaptive introgression”) (*12*–*17*). The evolutionary forces––e.g., genetic drift or natural selection–– that have shaped these patterns are not fully understood.

Most of the previous studies have focused on present-day individuals, where separating the effects of past demography and selection is challenging (*18*). Here, we generate a catalog of Neandertal ancestry in 59 ancient (sampled between 45,000–2,200 years before present (yBP)) and 275 diverse present-day modern humans, providing a systematic analysis of Neandertal ancestry through time and space. We recover the origin and trajectory of variants inherited from Neandertals, which allows us to refine the estimates of when the gene flow occurred (*7, 9, 19*) and directly observe how selection has shaped the patterns of ancestry across the genome (*18, 20*). Together, these analyses help characterize the population history and legacy of Neandertal gene flow in modern humans.

### Identifying the location of Neandertal ancestry in modern humans

We use genomic data from 59 ancient modern human individuals ranging between 45,000–2,200 yBP, including 33 individuals that are older than 10,000 years. We also include the genomes from 275 diverse present-day individuals from worldwide populations that are part of the Simons Genome Diversity Project (SGDP) (Materials and Methods Section 1). Our data set contains a combination of whole-genome sequences and variants enriched for target positions using two different SNP capture arrays; the “1240k” (containing ∼1.2 million sites segregating in modern humans) and the archaic admixture array (containing ∼1.7 million Neandertal ancestry informative sites (*8*)) (table S1, Materials and Methods Section 2). We cluster individuals into 14 population groups that are stratified by geographic location and time using the data on the 1240k array (Fig. 1A, fig. S10, table S8, Materials and Methods Section 4.1).

**Fig. 1:**
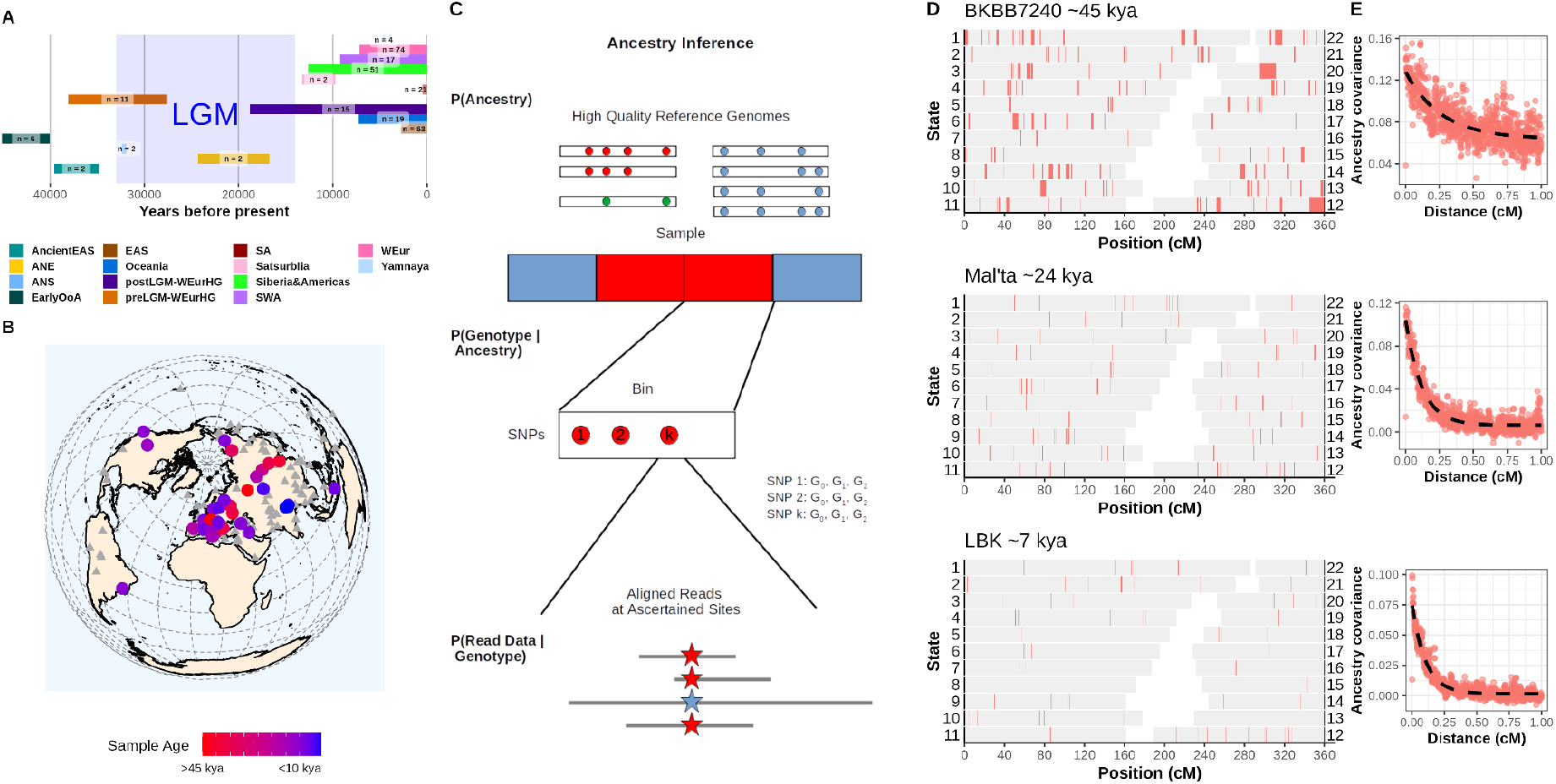
Calling of Neandertal ancestry in ancient and present-day individuals world wide. (**A**) Sample overview of the population clusters through time. ancient East Asians (ancientEAS), ancient North Eurasians (ANE), ancient North Siberians (ANS), early out of Africa (EarlyOoA), East Asians (EAS), post-Last Glacial Maximum West Eurasian Hunter-Gatherers (postLGM-WEurHG), pre-Last Glacial Maximum West Eurasian Hunter-Gatherers (preLGM-WEurHG), South Asians (SA), Southwest Asians (SWA), West Eurasians (WEur). Bars represent the span from oldest to youngest individual; *n* is the number of individuals. (**B**) Sample location and age of individuals. Triangles: present-day individuals, dots: ancient individuals (**C**) Workflow overview for *admixfrog*. High quality genomes are used as representatives for the unobserved ancestries (blue, red and green) of a given target genome. The genotypes of all SNPs in a small bin are latent states that are coestimated from aligned sequencing reads. (**D**) Inferred ancestry segments across the autosomes in three ancient modern human individuals. Red segments represent Neandertal ancestry. (**E**) Ancestry covariance for the same individuals calculated on the genotype likelihoods from *admixfrog*. The dotted line indicates the inferred decay of coancestry.

To infer Neandertal ancestry segments in a single target modern human genome, we use *admixfrog*, a hidden Markov model-based approach (*21*). For each diploid individual at each window (0.005cM), *admixfrog* estimates a combination of two ancestries from three possibilities: i) Neandertal (using the three high-coverage Neandertal genomes as reference (*2*–*4*)), ii) Denisovan (using the high-coverage Altai Denisovan genome (*6*)), or iii) modern human (using a panel of individuals of sub-Saharan African-related ancestry who have minimal Neandertal ancestry (*22*)). *admixfrog* coestimates genotype likelihoods and contamination, and is thus well-suited for ancient DNA (Fig. 1C Materials and Methods Section 3.1).

We performed extensive simulations to test the performance of *admixfrog*, and find that the method works reliably for shotgun genomes and the archaic admixture array, for samples with a coverage of at least 0.2x, but it has lower power for 1240k capture array that has few archaic informative markers (fig S2-8, table S1 and S3, Materials and Methods Section 3.1 - 3.3). Moreover, since we have access to only one Denisovan genome that is highly diverged from the introgressing Denisovan into most human populations (*23*), our ability to reliably detect Denisovan ancestry is limited (fig. S9, table S6). Thus, we focused on Neandertal ancestry only. Using 58 ancient and 231 present-day non-African individuals (Fig. 1B), we generated a comprehensive catalog of Neandertal ancestry segments that we use to infer the source, timing and function of Neandertal ancestry in modern humans (Fig. 1D,E).

### Spatiotemporal patterns of Neandertal ancestry

The distribution of Neandertal ancestry across populations offers insights into the historical interactions between modern humans and Neandertals. After the initial gene flow, Neandertal variants would be shaped by the demographic history of the modern populations including genetic drift, bottlenecks and secondary gene flow events. Neandertal segments that originated from the same introgression event would be shared by descendant populations and the sharing patterns would thus reflect the relatedness observed at random neutral sites in the genome. The amount of unique Neandertal ancestry in any individual would in turn be small. In contrast, secondary Neandertal gene flow events (private to some populations) would introduce ancestry at new genomic locations, and would thus lead to populations with largely uncorrelated ancestry patterns and increased level of unique ancestry (*24*). Furthermore, gene flow events from genetically differentiated Neandertal populations would result in differences in divergence estimates between the introgressing segments and the reference Neandertal genomes (*23*).

We examine the population structure at Neandertal ancestry segments using Principal component analysis (PCA). We find PC1 differentiates individuals from East Asia and Europe and PC2 separates individuals from Oceania and other worldwide groups. The differentiation observed in PCA is driven by differences in frequency, rather than the presence/absence of Neandertal segments (fig. S11). Moreover, the correlation matrix of shared Neandertal ancestry segments is highly concordant with the *f*_*3*_-drift-matrix (inferred using 1240k sites) measuring genome-wide allele sharing across individuals (Pearson correlation = 0.78, p < 2.2e-16, Materials and Methods Section 4.2 and 4.3, fig. 2C, fig S12, table S7-9). The notable exception is the EarlyOoA cluster (that includes individuals with sampling age older than 40,000 yBP such as Ust’ Ishim, Oase, Bacho Kiro and Zlatý kůň) which has a significantly weaker correlation (mean correlation = 0.05, maximum p-value < 0.0012 Holm-Bonferroni adjusted t-test, table S10).

**Fig. 2:**
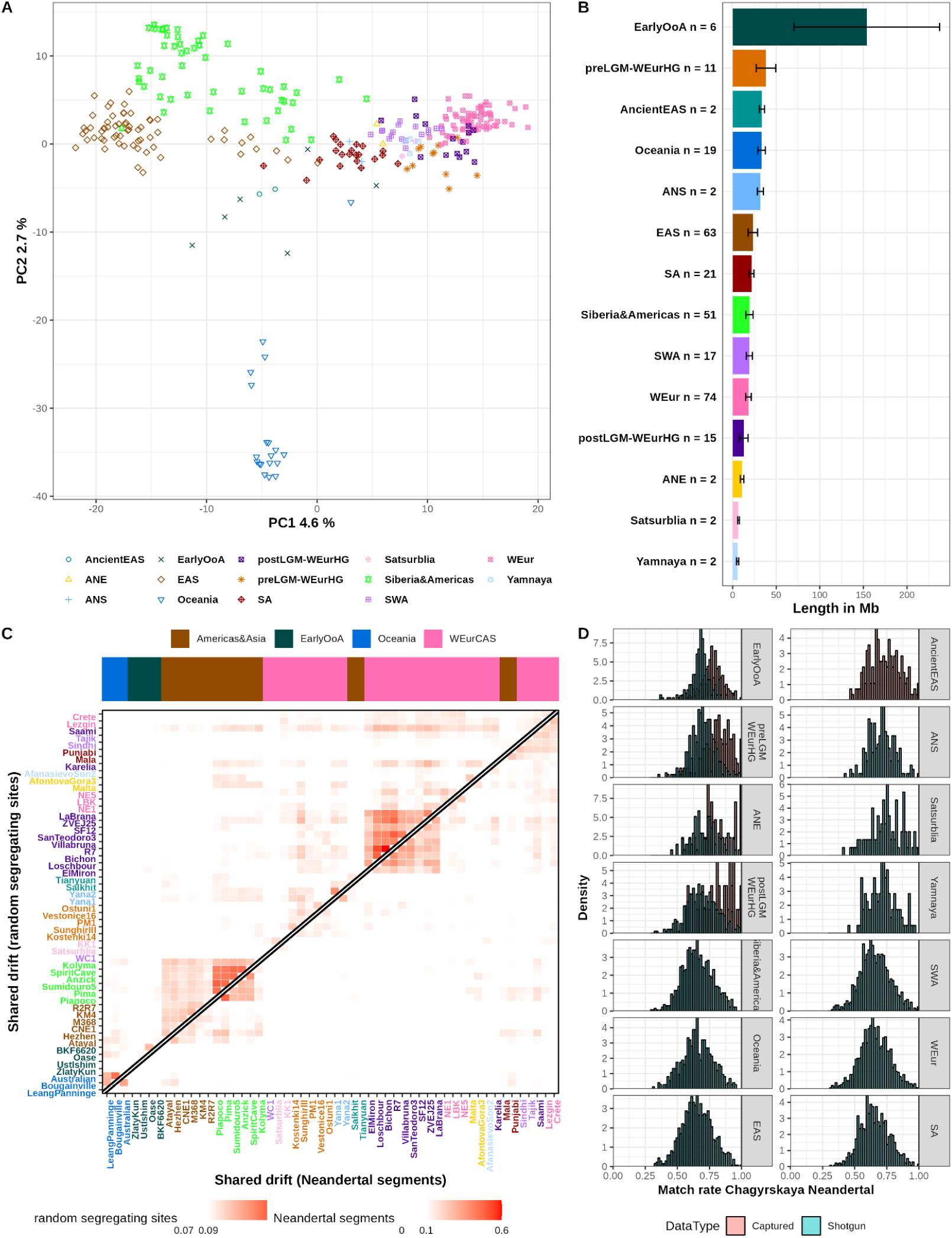
Analysis of the spatiotemporal pattern on inferred Neandertal segments on the autosomes. (**A**) PCA of the sharing of Neandertal segments. (**B**) Amount of unique Neandertal ancestry per population cluster (with *n* giving the number of individuals per cluster) for a randomly sampled individual. Error bars are calculated by resampling the individuals. (**C**) Comparison between the differences in pairwise f_3_ values (upper part of matrix) and the pairwise correlation of Neandertal segments (lower part of matrix). One individual per site. Two random present-day SGDP individuals are included for clusters containing present-day samples. (**D**) Matching rate of Neandertal segments with at least 15 informative SNPs to the Chagyrskaya Neandertal, stratified by data type (shotgun sequenced in blue or captured in red). Individuals are grouped in population clusters ordered from oldest to youngest. Population clusters have the same labels as in Fig 1.

Comparing the sharing of Neandertal ancestry segments across individuals, we find that the proportion of unique Neandertal ancestry in most individuals that postdate the Last Glacial Maximum (LGM) including present-day individuals is low at an average of 6%, with little variation between population clusters (Materials and Methods Section 4.4 fig. S13, table S11). Individuals that predate the LGM have a higher amount of unique Neandertal ancestry, with the highest proportion in EarlyOoA individuals (296 MB or 34%). The results remain significant even after controlling for sample size ( p-value < 0.0002 for pairwise t-test with Holm-Bonferroni multiple testing correction) (Fig. 2B, table S12 and 2.11).

Among post-LGM individuals, we find that the amount of unique Neandertal ancestry is not significantly different between West Eurasians and East Asians, despite East Asians having ∼20% greater overall Neandertal ancestry (22 Mb unique in EAS vs 19 Mb unique in WEur, p-value = 1) (table S13). Interestingly, there is no significant difference in the overall Neandertal ancestry between ancient East Asians and pre/post LGM West Eurasian Hunter-Gatherers, albeit the sample size is small. Furthermore, we find the largest amount of unique Neandertal ancestry in the Oceanian cluster, possibly due to a contribution of some misclassified Denisovan ancestry segments (*6*) (table S6). The lowest amount of unique Neandertal ancestry per individual is seen in Satsurblia and Yamnaya clusters. The Caucasus Hunter-Gatherer ancestry in these population clusters is widespread in present-day individuals, with substantial contributions in West Eurasians and South Asians in our data (*25, 26*).

Next, we investigated whether a single or multiple Neandertal populations have contributed to the introgression. We calculated the number of differences from inferred segments to the Chagyrskaya Neandertal (which was not used in the design of the archaic admixture array) and find a unimodal distribution in all clusters (Fig 2D, Materials and Methods Section 4.5), except the EarlyOoA cluster that has ∼6% of segments that are more diverged from Chagyrskaya (table S14). We find consistent results when comparing the proportion of introgressed segments coming from any of the four reference archaics estimated using a Generalized Mixed Linear model (GLM) (fig S16 and S18, table S15 and 2.14).

The EarlyOoA cluster has large amounts of unique Neandertal ancestry, a significantly different matching profile to the sequenced Neandertals, and the weakest correlation in the locations of introgressed segments with other population clusters. These observations are consistent with the insight that some Neandertal ancestry in these older individuals is not shared with modern humans after 40,000 years (*7*–*9*).

### Timing of Neandertal gene flow

We infer the timing of Neandertal gene flow by measuring the ancestry covariance between pairs of markers across the genome ((*19, 27, 28*), see Materials and Methods Section 5.1-5.2). As estimates become noisier and potentially biased in younger individuals (*19*), we focussed on 22 individuals older than 20,000 yBP for this analysis. Moreover, some EarlyOoA individuals have very recent Neandertal ancestry and possibly evidence for multiple pulses of gene flow (*7*–*9, 19*), thus we analyze these individuals separately.

If we assume the gene flow occurred instantaneously (IG model), we expect the decay of ancestry covariance in each individual to follow an exponential distribution. By measuring the ancestry covariance for each of the 16 ancient individuals that lived between 40,000 and 20,000 yBP, we infer that the Neandertal gene flow occurred between 321 and 950 generations before these individuals lived (Materials and Methods Section 5.1 and 5.2 fig S.21, table S17). By leveraging the linear relationship between the dates of Neandertal gene flow (in generations) and individuals’ sampling age (in years), we jointly infer the average generation interval as 28.4 years [95.5 % CI: 27–30 years] and the time of the shared pulse of Neandertal gene flow as 46,364 yBP [95.5 % CI: 45,682-47,045 yBP]. This is consistent with previous estimates (*19, 28*), though more precise due to our larger sample size (Fig. 3A). We find our results are robust to the inclusion of individuals in the EarlyOoA cluster (table S18).

**Fig. 3.**
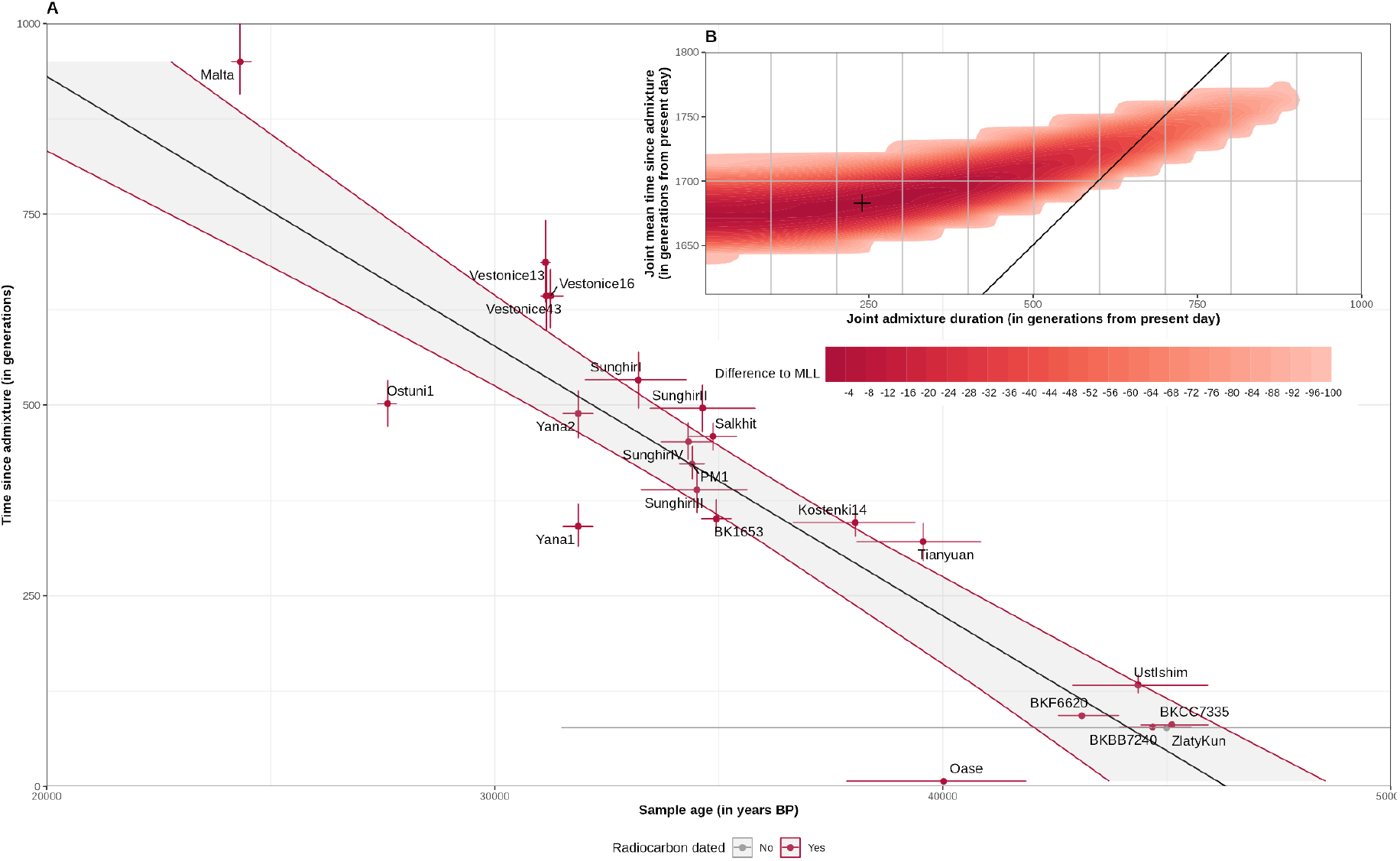
Dating of the Neandertal admixture event in 22 individuals older than 20,000 yBP. (**A**) Time since the Neandertal admixture using ancestry covariance curves with 95.5% confidence intervals (y-axis) versus the age of the individual and the 95.5% confidence intervals (x-axis). The date for Zlatý kůň is a genetic date. Black line indicates the fit of a linear model with the uncertainty in gray shades. (**B**) Log-likelihood surfaces of the joint estimate of gene flow duration under the extended pulse model with the Early out of Africa (EarlyOoA) individuals excluded (n = 16). Shades of red indicate the difference between any log-likelihood and the highest log-likelihood. Black diagonal line indicates the inferred timing of Neandertal disappearance (*30*).

To determine whether these estimates would differ if admixture with Neandertals took place over an extended duration, we next compared the IG model to the extended pulse (EP) model which assumes gene flow occurred over multiple generations (*29*). We focused on individuals younger than 40,000 yBP, since the joint modeling of the IG and EP models assumes that no major gene flow occurs after the sampling age of the oldest individual (due to large uncertainty in the sampling age of EarlyOOA individuals, we excluded them). We obtained a significantly better fit for the EP model than the IG model (Likelihood ratio test, p < 2.2e-16), with the mean time of gene flow of around 47,124 yBP [46,872–47,404 yBP], and a duration of around 6,832 years [2,044–9,968 years]. This timing is compatible with archaeological evidence for the overlap of modern humans and Neandertals in Europe (*30*). We note, however, that uncertainties in local recombination rates over time and sampling ages of ancient individuals make the estimates of duration tentative (Fig. 3B; fig. S24 and S26; table S21 and S22) (*31, 32*).

In summary, the sharing of Neandertal segments mirrors the population structure among non-Africans and supports a single major Neandertal gene flow event into the common ancestors of all surviving lineages of non-Africans that occurred ∼47,000 years ago with a duration of ∼6,800 years. This gene flow continued, to some extent, as early modern humans spread throughout Eurasia but did not leave detectable traces in later populations.

### Neandertal ancestry across the genome

Neandertal ancestry plays a major role in human adaptation and disease (*33, 34*), but few studies have tracked how the frequency of Neandertal variants has changed through time (*35*–*37*). Using Neandertal segments in ancient and present-day individuals, we recover Neandertal ancestry in 61.7% (1,551 Mb) of the autosomal callable genome (fig. S27). On the X-chromosome, we find Neandertal ancestry only in 18.7% (29Mb / 154.84 Mb) of the genome. The distribution of Neandertal ancestry segments on X chromosome is non-uniform and non-random distribution, with large regions devoid of any Neandertal segments (Fig 4A). Indeed, when we measure entropy on the X chromosome vs. autosomes, we find the distribution on X is significantly more ordered (Shannon’s entropy H = 0.03 (X-chromosomes) vs. 0.11 (autosomes), Wilcoxon rank sum test p-value < 2.2e-16).

**Fig. 4.**
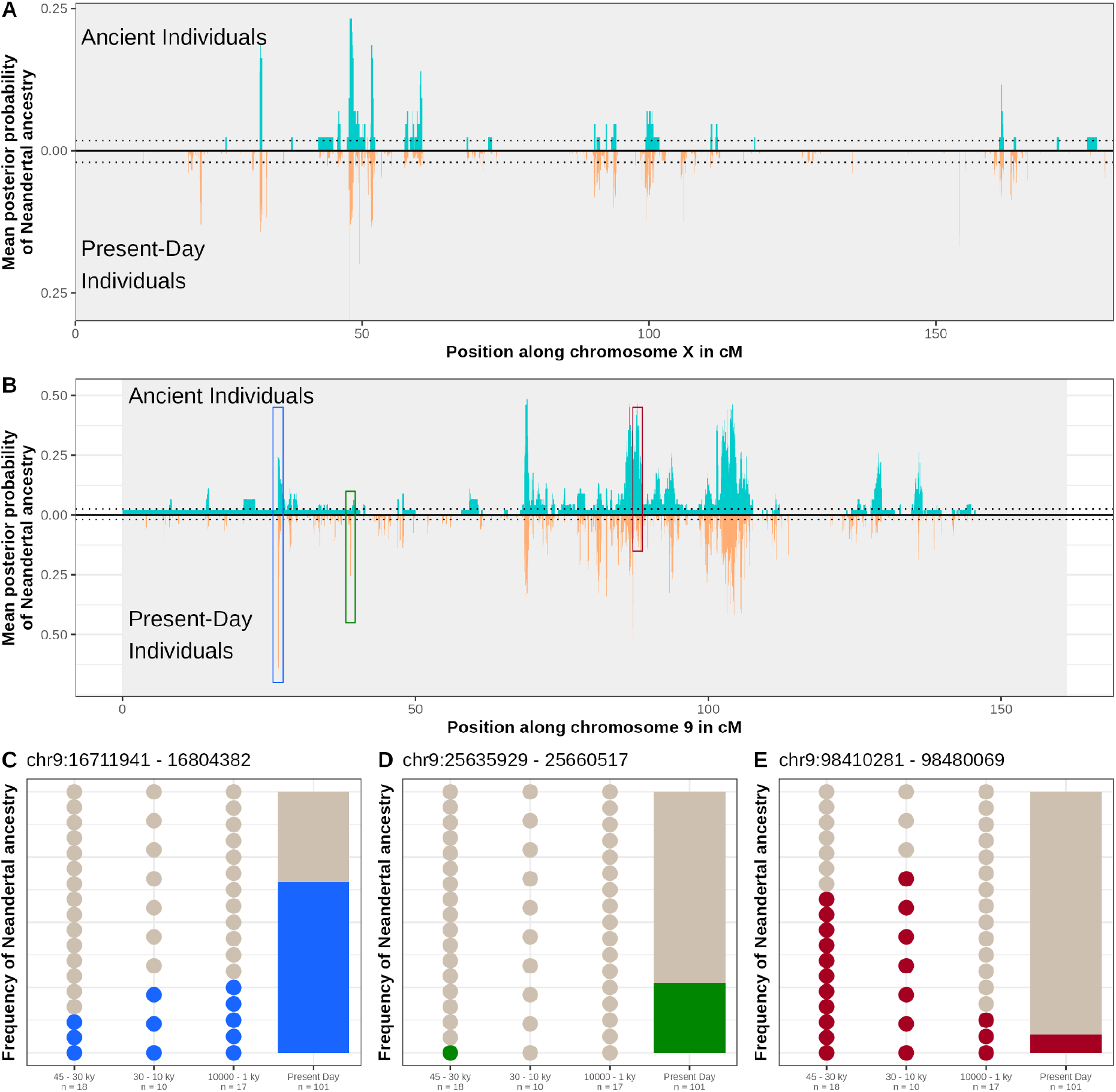
Regions of high Neandertal ancestry through time. (**A**) Mean posterior probability of Neandertal ancestry on chromosome X for ancient (teal) and present-day (orange) individuals. The dotted lines give the average posterior probability for Neandertal ancestry throughout the genome. **(B)** Mean posterior probability of Neandertal ancestry on chromosome 9. Colored rectangles indicate the position of the high-frequency regions **(C)** regions of high frequency in both present-day and ancient genomes. An example of such a region contains gene *BNC2*, shown in the figure. **(D)** regions at high frequency in present-day, but not ancient individuals, and **(E)** regions at high frequency in ancient, but not present-day individuals. Number of dots corresponds to the number of individuals at the sampling time depth shown on the X-axis. Genomic coordinates are shown in human genome reference build, hg19.

In present-day individuals, previous studies have shown that the landscape of Neandertal ancestry is correlated with recombination rate (*38*) and *B*-statistics, a measure of background selection (*12, 39, 40*). We find that local recombination rate is positively correlated with Neandertal ancestry throughout time, except for EarlyOoA where we lack power (table S23, Materials and Methods Section 6.2). Across time intervals, we also find that regions under constraint (low B-scores) consistently harbor less Neandertal ancestry compared to the rest of the genome(fig. S29). For instance, in individuals older than 30,000 yBP, mean Neandertal ancestry in the lowest B-score bin is 3.2% and in the highest B-score bin is 6.2%, suggesting that initial gene flow may have been >5% in modern humans. For the X chromosome, we find a weak correlation with recombination rate at fine-scales (*r*^*2*^ at 20 kb = 0.05, *p*-value = 0.04) but the correlation is not significant at larger distances due to limited power (table S33).

To identify candidate regions of natural selection, we examined how the frequency of Neandertal segments changed with time. Segments that harbor beneficial alleles may increase in frequency as a result of positive selection or adaptive introgression, while segments carrying deleterious alleles are predicted to be purged quickly, leading to Neandertal deserts (*12*). We thus scanned for regions where the frequency of Neandertal ancestry is unexpectedly high (or low) compared to the genome-wide average estimates or has changed drastically over time (Fig. 4B, table S24).

We identified 86 regions (347 genes) that are at high-frequency (99.9th percentile, Z > 4.5) in both present-day and ancient individuals (Fig. 4B,C, Materials and Methods Section 6.3). Using Gene Ontology (GO) analysis, we find these candidate regions are enriched for pathways related to skin pigmentation, metabolism and immunity (table S25). These pathways have also been identified in surveys of present-day individuals (*12, 13*), suggesting that many of these genes were immediately beneficial to modern humans as they encountered new environmental pressures outside Africa.

We find 91 candidate regions (169 genes) that are present at high frequency in present-day individuals but not in ancient individuals, indicating that these regions may contain variants that became adaptive later on (Fig 4B,D). We also find 32 candidate regions (102 genes) that were at high frequency in ancient DNA individuals but not in present-day individuals (Fig. 4B,E). Many of these regions (∼44%) are located within 1Mb of candidate regions at high frequency, suggesting that these haplotypes hitchhiked with beneficial mutations and decreased in frequency as recombination occurred.

Examining the trajectory of 11 previously published candidate regions of adaptive introgression inferred in multiple surveys of present-day individuals, we replicate 72% of the candidate regions, including seven that were immediately selected and one region of selection on standing variation. Among these regions, the most significant is a 2 Mb region on chromosome 2 where the highest Neandertal ancestry in ancient individuals is 64% and in present-day individuals is 67%. This region contains 12 genes, including TANC1 and BAZ2B, that have been associated with intellectual disability and autism disorders (*41, 42*). Another example is *BNC2*, a gene that plays a role in skin pigmentation (*43*), that is at ∼25% frequency in EarlyOOA and ∼65% in present-day individuals (Fig 4C), indicating that variants at this locus may have been immediately beneficial and increased over time in modern humans, unlike previous reports (*37*) (fig. S28, table S26).

To understand the genesis of regions of low (or no) Neandertal ancestry, we examined Neandertal ancestry over time in five putative deserts identified in previous studies (*13, 15*). We find that the deserts are in the 0.1th percentile of the empirical distribution of archaic ancestry genome-wide across time, including in individuals in the earliest time intervals (50,000–30,000 yBP) (table S30). Notably, we find almost no introgressed Neandertal segments within the boundaries of four out of five deserts in ancient or present-day individuals (fig. S30) (similar to a recent study in South Asians (*44*) we do not replicate the desert on chromosome 1). This indicates that the deserts formed rapidly after the initial gene flow, consistent with the theoretical expectations (*18, 45*).

We find that Neandertal ancestry on the X chromosome is already depleted in EarlyOoA individuals, and the X-to-autosome ratio of Neandertal ancestry remains stable over time (0.229–0.408 for females and 0.131–0.147 for males) (fig. S34). Concordantly, we find large regions that are depleted of Neandertal ancestry in our earliest time intervals (fig. S37). Most of these regions overlap with previously reported putative selective sweeps (*46*). Among the two previously identified deserts on the X chromosome (*13*), only one of them is completely devoid of Neandertal ancestry (chrX:62,000,000-78,000,000), while we find substantial Neandertal ancestry in the other (chrX:109,000,000-143,000,000) (fig. S38).

In summary, the majority of positive and negative selection on Neandertal ancestry happened very quickly, and left clear signals in the genetic diversity of the first modern humans outside Africa. Only a smaller proportion of variants became adaptive later on.

## Acknowledgments

We thank Janet Kelso, Molly Schumer, Nick Patterson, Stéphane Peyrégne, Monty Slatkin, Elena Zavala, Sarah Johnson and Elise Kerdoncuff for helpful comments on the manuscript.

## Data and materials availability

No new data was generated for this study. The catalog of Neandertal segments will be publicly available through Dryad DOI: 10.5061/dryad.zw3r228gg. All scripts are publicly available through github: https://github.com/LeonardoIasi/Neandertal-ancestry-through-time.

## Supplementary Materials

**This PDF file includes:**

Materials and Methods

Figs. S1 to S38

Tables S3,S18,S22,S30,S31,S31,S32

References (47–91)

### Other Supplementary Material for this manuscript includes the following

Tables S1,S2,S4 to S17, S19 to S21, S23 to S29

