## Supplementary Materials for "Neandertal ancestry through time: Insights from genomes of ancient and present-day humans"

**This PDF file includes:**

Materials and Methods

Figs. S1 to S38

Tables S3,S18,S22,S30,S31,S31,S32

References (*47–91*)

**Other Supplementary Material for this manuscript includes the following:**

Tables S1,S2,S4 to S17, S19 to S21, S23 to S29

#### Table of Contents

**Materials and Methods** **25**
1. Data 25
    1.1 DNA sequence data 25
    1.2 Recalibration of radiocarbon dates 26
2. SNP Ascertainment 26
    2.1 Archaic ascertainment 26
    2.2 ArchaicX ascertainment 26
    2.3 Coverage and properties of the archaic admixture + ArchaicX array 27
    2.4 1240k ascertainment 28
    2.5 “All diagnostic sites” ascertainment 28
    2.6 Acov ascertainment 28
3. Local ancestry inference 28
    3.1 Local ancestry inference using admixfrog 28
    3.2 Calling Fragments 29
    3.3 Comparison to 1240k 29
    3.4 Ancestry proportions 31
    3.5 Comparison to hmmix 32
4. Spatiotemporal patterns of Neandertal ancestry 33
    4.1 Clustering of individuals into genetically informed groups 33
    4.2 PCA of Neandertal Ancestry 34
    4.3 Correlation of Neandertal Segments 34
    4.4 Identification of unique introgressed segments 35
    4.5 Archaic lineage assignment of introgressed segments 36
        4.5.1 Match rate distribution 36
        4.5.2 Proportions of Segments matching the reference archaic individuals 37
        4.5.3 Validation of lineage assignment 38
5. Genetic Dating of Neandertal Gene flow 39
    5.1 Ancestry covariance curves 39
    5.2 Single-sample dating 40
    5.3 Generation Time estimate 40
    5.4 Assessment of instantaneous gene flow (IG) vs. extended admixture pulse (EP) model 40
        5.4.1 Simulated ancestry segments 40
        5.4.2 Simulated Ancestry Covariance 41
        5.4.3 Adding Noise 41
        5.4.4 Application to real data 41
            5.4.4.1 Fitting an Instantaneous gene flow model 42
            5.4.4.2 Fitting the Extended pulse model 43
            5.4.4.3 Comparing IG and EP model 44
6. Neandertal Ancestry across the genome 44
    6.1 Total recovered Neandertal genome and total deserts 44
    6.2 Neandertal frequency correlation 44

6.2.1 B statistics 44
6.2.2 Recombination rate 45
6.3 Analysis of high frequency regions 45
6.3.1 Outlier estimation 46
6.3.2 Comparison to previously identified regions 47
6.3.4 Gene Ontology Enrichment 47
6.3.5 Trajectory of archaic frequency over time 48
6.4 Deserts through time 48
7. Neandertal ancestry on the X chromosome 49
**Supplementary Figures** **51**
**Supplementary Tables** **89**

### Materials and Methods

#### 1. Data

##### 1.1 DNA sequence data

In total, we assembled data from 59 ancient modern humans from 26 studies (*7–9*, *11*, *24*, *25*, *35*, *47–66*). We complemented these data with 275 present-day individuals from the Simons Genome Diversity Panel (SGDP) (*67*) (see table S1) and the 1000 Genomes Project (1k Genomes) (The 1000 Genomes Project Consortium et al. 2015). We also used the four high-coverage archaic genomes available to date: Vindija 33.19 (*3*), Denisova 5 (*2*) (henceforth, Altai Neandertal or Altai), Chagyrskaya 8 (*4*), and Denisova 3 (*6*). We used the genotype calls based on snpAD (*69*) published in (*3*, *4*). We applied the following filters: i) remove tandem-repeats and insertions-deletions, ii) filter for coverage stratified by GC-content (requiring at least 10-fold coverage) and select only mappable regions with unique 35-mers, (iii) remove genotypes with mapping quality of less than 25. For the ancient modern human genomes, we used the bam files containing aligned reads from the European Nucleotide Archive. For present-day individuals, we used the genotype calls in the vcf files for 1k Genomes and SGDP. SGDP individuals' genotypes were required to have a genotype quality score >= 1 and for the 1k Genomes >= 100 (as these individuals have a lower coverage). We assumed all present-day individuals have homozygous reference genotype for sites that are absent from the SGDP and 1k Genomes vcf files but present in the accessible genome. For all analyses, we used the hg19 human reference genome (https://www.ncbi.nlm.nih.gov/grc). To polarize our data to the ancestral allele, we used the great apes reference genomes aligned to the human reference genome, including chimpanzee (PanTro6), gorilla (GorGor3), bonobo (PanPan1) and orangutan (Pan Abe3) thate were downloaded from http://hgdownload.cse.ucsc.edu/goldenpath/hg19/. We used ancient samples that were either whole-genome shotgun sequenced or sequenced after capturing the libraries with the archaic admixture and the 1240k arrays (panel 4 and panels 1 and 2 respectively from Fu et al. 2015). For the shotgun data, we expanded the archaic admixture array to include sites on the X-chromosome ("ArchaicX").

##### 1.2 Recalibration of radiocarbon dates

To estimate the age of the ancient modern human individuals, we retrieved the uncalibrated radiocarbon dates plus the estimated error in years before present (yBP) (present here refers to the year 1950) or thousand years BP (kyBP). We re-calibrated all raw dates using OxCal v4.4.4 with the IntCal20 calibration curve (*70*, *71*) (table S2). We obtained radiocarbon dating data from several sources. In turn, some samples had multiple radiocarbon dates, and some others did not have direct dating information. For each individual, we thus defined a "most likely" date:

- For individuals with unique radiocarbon dates, the most likely date is the recalibrated radiocarbon date with its upper and lower bounds (95.5% highest posterior density interval).
- For individuals with multiple dates, the most likely date is the mean of the recalibrated dates, and we use the range of the highest upper and lowest lower bounds.
- In case of Yana1, Yana2, San Teodoro 3, I3388 (Afanasievo Mother), Karelia, KM4, KS20_KS25, KS5, M2113 and M368, there are indirect dates for the layers these specimens stem from, and which we then take as most likely dates.
- In case of NE1 and NE5, no uncalibrated dates were available. The most likely date in these cases is the published calibrated date.
- In the case of Zlatý kůň, there are no reliable carbon dates due to contamination, therefore we used the date from the original publication (*65*).

#### 2. SNP Ascertainment

We used five different SNP ascertainment schemes. The main ascertainment used in this study is the archaic admixture array (panel 4 of (*8*)). We combined this array with the new "ArchaicX" ascertainment that contains variants on the X chromosome. The combined ascertainment is referred to as archaic admixture + X that we used for local ancestry inference in all individuals. For a subset of analyses, we also used the 1240k ascertainment described in (*8*) and the all diagnostic sites ascertainment. For dating the admixture between modern humans and Neandertals, we used the ACov ascertainment to calculate the ancestry covariance statistics (*65*). All ascertainments were restricted to the mappability track map35_99 . We describe each of these below.

##### 2.1 Archaic ascertainment

This ascertainment (panel 4) capture set contains ~1.7 million archaic ancestry informative sites, i.e. sites where at least one of the archaic genomess (Denisova; Altai, low coverage Vindija and Mezmaiskaya Neandertals) carries one allele but nearly all individuals in a panel of sub-Saharan Africans carry a different allele (*8*).

##### 2.2 ArchaicX ascertainment

The archaic admixture capture set contains ~1.7 million archaic ancestry informative sites across all autosomes. It, however, does not include any SNPs on the X-chromosome. Thus, we augmented our sets of informative sites with a number of SNPs located on the X-chromosome. This "ArchaicX" ascertainment for the X chromosome is designed based on the four high-coverage archaic genomes together with 504 Sub-Saharan African-related genomes from the 1k Genomes (ESN, GWD, LWK, MLS and YRI), and a high-coverage Mbuti individual (HGDP0456) from the Human Genome Diversity Project (HGDP) (*6*). To call ancestral alleles, we used the chimpanzee and gorilla reference genomes.

Our pre-filtered SNP-set contains 83,324 biallelic SNPs that are polymorphic in the archaic hominins, 3,516 SNPs which are differentially fixed between archaic hominins and present-day Africans (i.e. SNPs where all sub-Saharan Africans carry the derived allele and all archaics carry the ancestral allele and vice-versa) and 25,122 SNPs which are segregating (i.e., not fixed) differences between the archaic hominins and present-day sub-Saharan Africans (i.e. SNPs with different derived frequencies). Specifically, the latter were included to estimate contamination rates, and are sites with at least one ancestral allele in each of the three archaic individuals (Denisova 3, Altai and Vindija 33.19) and at least one derived allele in Mbuti (HGDP0456), after randomly pseudo-haploidizing the genotypes where we assign a haploid genotype determined by randomly selecting one allele at the variant site.

For the archaic genomes, we removed tandem-repeats and insertions-deletions (indels), filtered for coverage (stratified by GC-content, requiring at least 10-fold coverage), and selected only mappable regions with unique 35mers and selecting reads with a minimum mapping quality of 25. A similar filtering was applied to the high-coverage individual, HGDP0456, except that the minimum mapping quality was set to 30. We also removed sites where we did not have consistent genotype calls across the chimpanzee and the gorilla reference genomes.

In anticipation of using these SNPs in a future capture array (with 60 base pair (bp) probes), we further set a minimum distance of 61 bp between SNPs (to mitigate potential reference bias due to a probe overlap), and removed sites where the GC content of 60 surrounding base pairs (29 bp upstream and 30 bp downstream of the SNP) was lower than 25% or higher than 75%. Moreover, we excluded SNPs in repetitive sequences using the UCSC genome browser tracks for tandem repeat, simple repeat (hg19, GCA_000001405.1) as well as DUST (*72*, *73*) Additionally, nuclear mitochondrial DNA segments (NumtS) as defined in (*74*) were also filtered out. The final X chromosome ascertainment contained 79,487 unique SNPs after filtering, including both transitions and transversions.

##### 2.3 Coverage and properties of the archaic admixture + ArchaicX array

The archaic admixture + X ascertainment contains in total 1,310,813 informative sites. The genome coverage, i.e. the distribution of sites targeted throughout the genome, is shown in fig. S1. We created a callable genome mask for our analysis by defining all 20 kb windows throughout the human genome with at least one target locus in the archaic admixture + X array. The transition to transversion ratio of the array is 1.64. We used the GenCode annotations to check for the amount of SNPs in our capture array targeting exonic regions. We sub-sampled the annotation file to all exons annotated in ENSEMBL. The proportion of exons using this filter is 0.0067 and the proportions of sites in exonic regions is 0.0047.

##### 2.4 1240k ascertainment

This ascertainment (panel 1 and 2) contains ~1.2 million sites segregating in modern humans (*8*). We filtered these SNPs for the mappability track map35_99 filter which results in 1,097,136 sites.

##### 2.5 "All diagnostic sites" ascertainment

The "all diagnostic sites" ascertainment consists of diagnostic sites between the high coverage genomes of the SGDP B_Mbuti-4 and the four high coverage archaic genomes including Altai, Vindija and Chagyrskaya Neandertals and the Altai Denisovan. For each individual, we removed tandem-repeats and indels. We only considered regions with uniquely mappable 35 mers and applied a GC content conditioned coverage filter of 10x. We followed the approach in (*75*) to construct the "diagnostic positions". We randomly sampled an allele from each individual and we only considered sites where all individuals have non-missing genotypes. To determine the ancestral alleles, we used the chimpanzee, bonobo, gorilla and orangutan reference genomes aligned to the hg19 human reference, where at least three species match with each other and at most one is missing. Only sites where at least one of the hominins has a derived allele were retained. The resulting sites were filtered again using the mappability track map35_99 filter. This results in 2,674,580 diagnostic positions.

##### 2.6 Acov ascertainment

This ascertainment includes SNPs that are informative of Neandertal ancestry and used for measuring ancestry covariance across the genome. Following (*19*), we used the "ascertainment 0" where the Altai Neandertal carries at least one derived allele and all individuals in a panel of Sub-Saharan Africans (1000 genomes YRI and LWK) carry the ancestral allele. This ascertainment contains 227,346 sites.

#### 3. Local ancestry inference

##### 3.1 Local ancestry inference using *admixfrog*

We performed local ancestry inference using *admixfrog 0.7.1* (*21*). *admixfrog* uses a hidden Markov model (HMM) to infer archaic ancestry segments in a diploid modern human genome, using high-coverage archaic and modern reference genomes. Specifically, the genomes are binned into small windows (here, 0.005 cM), and each window is assigned either a probability of homozygous or heterozygous ancestry between all pairs of possible ancestries (e.g., Neandertal-Neandertal, Neandertal-African, etc.). As many ancient individuals have been sequenced only to low-coverage and may be contaminated with present-day DNA, admixfrog co-estimates genotype likelihoods and the rate of contamination.

We built our reference panels as follows:
Denisovan (DEN): Denisova 3
Neandertal (NEA): Vindija 33.19, Altai Neandertal and Chagyrskaya 8.
Modern Humans/ African (AFR): We use all female individuals from the 1k Genomes including Mende, Mandenka, Yoruba, Esan and Luhya individuals.
Chimpanzee (PAN): chimpanzee reference genome *panTro6*.

We estimated the number of informative sites for three capture arrays including 1240k, archaic admixture + X and the "all diagnostic sites" sites. Given the paucity of informative sites for archaic ancestry in the 1240k, we restricted the downstream analyses to the sites in the archaic admixture + X capture arrays (see Materials and Methods Section 2, table S3). We used the chimpanzee reference genome (panTro6) to infer the ancestral allele. For sites where the chimpanzee does not align with hg19, we used a uniform prior between the alleles. The genetic distances on the autosomes were assigned using the European-specific Shared map (*32*) downloaded from https://www.chg.ox.ac.uk/~anjali/AAmap/ lifted over to hg19 . Since the Shared map does not include the X chromosome, we used the African-American map (AA map) for the genetic distances on the X.

For each individual, we ran *admixfrog* with the following parameter setting using the respective bam files as input:

admixfrog --states AFR NEA DEN --cont-id AFR --ll-tol 0.01 --bin-size 5000 --est-F --est-tau --freq-F 3 --freq-contamination 3 --e0 0.01 --est-error --ancestral PAN --run-penalty 0.1 --max-iter 250 --n-post-replicates 200 --filter-pos 50 --filter-map 0.000 –input {input_file} –ref {ref_file} –output {output_name}

##### 3.2 Calling Fragments

We obtained the posterior probability for each bin and each ancestry state (i.e., Neandertal-Neandertal, Denisovan-Denisovan, African-African and Neandertal-African, Neandertal-Denisovan, Denisovan-African). To call segments, we used a penalty for classification of archaic ancestry of 0.25 with the following setting:

admixfrog-rle --penalty 0.25 –input {input_file} –output {Output_name}

The penalty determines how likely it is that long ancestry segments should be split up. To call Neandertal/Denisovan fragments, we required regions to be either homozygous or heterozygous for that ancestry (using the "state" calls in admixfrog). Unless stated otherwise, we only kept segments with a minimum length of 0.05 cM (~50 Kb) for present-day individuals or 0.2 cM (~200 kb) for ancient individuals. For each segment, we then recorded the total length (in cM and bp) of the called segments overlapping the bin, the total number of SNPs and the number of SNPs matching the AFR, NEA or DEN reference genomes. We only kept archaic ancestry (Neandertal and Denisovan) bins that overlapped an archaic ancestry segment that passes all the filtering criteria, including the minimum length cut-off. Otherwise, the posterior probability for the bin was set to zero.

##### 3.3 Comparison to 1240k

We compared the effectiveness of the archaic admixture + X array sites to the 1240k array sites that are tailored to capture modern human variation on the autosomes. For our evaluations, we used two high-coverage ancient genomes for whom we have whole genome sequences: 37x coverage Ust'-Ishim (*7*) and 20x coverage Loschbour (*11*). We compared the 1240k and archaic admixture + X ascertainment to an "all-diagnostic sites" ascertainment which uses all sites that are variable between the sequenced high-coverage Denisovans and Neandertals and a present-day Mbuti individual. To investigate the effect of coverage, we downsampled Ust'-Ishim and Loschbour to a proportion of 1, 0.2, 0.1, 0.02, 0.01 and 0.005 of the original coverages. We then classified all the segments inferred from the ascertained ("test") data in comparison to the full-genome ascertainment ("reference") without a length cutoff as follows: missing in test (MT) if there is no test segment overlapping a reference segment, missing in reference (MR), if we infer segment from the test data that do not overlap any segment in the reference data, and match (M) if a segment from the test data and high-coverage reference data uniquely overlap each other. We also classify a segment as a gap (GAP) if there are two or more test segments overlapping one high-coverage reference segment and segments are classified as being merged (MERGED) if one segment inferred from the test data overlaps two or more segments from the reference data . We find that the 1240k tends to miss or split up segments at low coverage, especially for the younger Loschbour individual, whereas the archaic admixture + X ascertainment accurately classifies segments even for coverage down to 0.2x (fig. S2).

To assess the impact of the penalty used for calling the segments, we compared the results of all possible combinations of penalties 0.2, 0.25, 0.3 and 0.4 for called segments between the original and downsampled data. Based on these results, we find that a penalty of 0.25 provides the best trade-off between calling segments and keeping the rates of false merged segments low (fig. S3).

We also tested the impact of uncertainty in the ancestral state of an allele on the inference. If the ancestral allele is unknown or *admixfrog* is run without it, a symmetric prior is used for the frequency at both reference or alternative alleles at the locus. We examined the difference in local ancestry inference with varying coverages for: (i) missing ancestral state (error2NOAncestralInfo), (ii) only considering sites where the ancestral state is known (error2UnpolarizedSNPsRemoved) to (iii) default of using the ancestral state if available and defaulting to a uniform prior if information is not available (used as reference here) (fig. S4). We do not find many differences in the overall results across the three setups and thus we chose to use the approach to consider all sites and use the ancestral state as the prior if it is available to maximize the observed data for low coverage samples.

Further, we tested the impact of using different genetic maps. We compared the inference using the European-specific Shared map or the African-American map. Fig. S5 shows the matching of the segments inferred using the African-American map compared to the segments inferred under the Shared map as a function of target coverage. Fig. S6 compares the distribution of the length of the ancestry segments inferred using the two genetic maps. We find there is very little difference in the called ancestry segments and also the inferred length of segments for Ust'-Ishim. For Loschbour, we find that when using the Shared map the overall length of inferred segments tends to be a little bit shorter compared to the African-American map. An explanation is that the Shared map is more specific for individuals with west Eurasian related ancestry, whereas the African-American map is enriched for recombination events in sub-Saharan African related ancestry groups and includes hotspots that are not active in west Eurasians (*32*)). For Ust'-Ishim, where the Neandertal ancestry segments are long, the accuracy of the genetic map has limited impact. However, for Loschbour, the ancestry segments are short and hence the choice of the recombination map has a more pronounced effect.

Finally, we tested the impact of heterogeneous coverage of SNPs across a target genome. Therefore, we calculated Pearson's correlation between the number of SNPs per bin in the accessible genome and the sum of the posterior probability of Neandertal ancestry (either homozygous or heterozygous) for all SNPs in the bin, for the downsampled Loschbour and Ust'-Ishim genomes using the *cor* function in *R* (*76*). Figure S7 shows that there is no substantial correlation between the number of SNPs in a genomic region and the probability of Neandertal ancestry.

##### 3.4 Ancestry proportions

To compare the proportion of Neandertal ancestry on the autosomes over time, we considered the overall posterior probability of being in the Neandertal ancestry state provided by admixfrog (either Neandertal-African or Neandertal-Neandertal) and compared it to $f_4$-ratio test (*77*) under multiple setups: i) using admixfrog input file for the archaic admixture array + X ascertainment, ii) same ascertainment as (i) but restricted to deaminated reads, and iii) by taking the highest genotype likelihood provided by the admixfrog output file. We follow Petr et al. 2019 and computed the $f_4$-ratio in the form of *$f_4$(Altai Neandertal, Chimp; X, Africans) / $f_4$(Altai Neandertal, Chimp; Vindija 33.19 Neandertal, Africans),* where *Africans* are all female individuals from the 1k Genomes including Mende, Mandenka, Yoruba, Esan and Luhya and X = is the test individual *(*see table S4*)*.

Given the population cluster we define (see Materials and Methods Section 4.1), we estimated the mean amount of Neandertal ancestry per population cluster using the overall Neandertal ancestry estimates from *admixfrog*, $f_4$-ratios on the archaic admixture +X sites and in 1240k sites. To compare the mean values we used a pairwise t test with Holm-Bonferroni multiple testing correction as implemented in the *pairwise.t.test* function from the stat package in *R* (table S5). Ancient (the Mota individual) and modern individuals in the cluster of Sub-Sahara-Africa did only show very low amounts of Neandertal ancestry with very short segments detected. North-African individuals show a higher amount of Neandertal ancestry but lower than non-African clusters. For all further analysis we focused on non-African individuals with 58 ancient and 231 present day individuals, if not stated otherwise.

We correlated the proportion of ancestry with the coverage, contamination and sample age (fig. S8). We excluded individuals from the correlation that have been previously suggested to have more than one admixture event with Neandertals (*8*, *9*). We detect a slight decrease in Neandertal ancestry over time both for the $f_4$-ratio test and admixfrog posterior probabilities (fig S8). This slight ascertainment bias falls into the variation observed in (*36*). We find that the ancient Oceanian individual from Leang Panninge is an outlier; however, it is an individual with the lowest coverage (0.2x) and highest level of present-day contamination (40%), and the only one where the overall proportion of Neandertal ancestry estimated using admixfrog and the $f_4$-ratio test differ from each other. We therefore excluded this individual from the correlations and did not consider it for analysis, if it is the only representative of a cluster (i.e. ancient Oceania). Further, we detect a correlation with coverage and contamination, though the direction of the effect is opposite to the expected effect. We expect contamination to reduce archaic ancestry rather than increase it as observed (fig S8). An explanation for this is that when an individual's age increases, the chance of detecting Neandertal ancestry increases, too, since this ancestry is in longer segments which are then easy to detect. However, older individuals are also usually of lower coverage (Pearson's correlation between age and coverage = -0.284, p-value < 0.04) and with a higher contamination (Pearson's correlation between age and contamination = 0.417, p-value < 0.002) i.e. contamination and coverage are collinear with age.

##### 3.5 Comparison to *hmmix*

We compared the putative introgressed archaic segments identified by *admixfrog* to those called by another published method, *hmmix* (*78*). We first identified all positions with derived variants found in individuals with sub-Saharan ancestry from 1k Genomes and HGDP (*79*) and lifted over the variants from hg38 to hg19.

*hmmix create_outgroup -ind={individuals} -vcf={vcffiles} -weights={weightsfile} -out={outgroupfile} -ancestral={ancestralfiles} -refgenome {refgenomes}*

where:

- "individuals" is a list of all individuals with sub-Saharan ancestry from 1k Genomes (n=426 from the following populations YRI, MSL and ESN) or HGDP (n=64 individuals from Africa, excluding Mozabite)
- "weightsfile" is the strict callability mask: ftp.1000genomes.ebi.ac.uk/vol1/ftp/release/20130502/supporting/accessible_genome_masks/20141020.strict_mask.whole_genome.bed
- ancestral information ftp://ftp.ensembl.org/pub/release-74/fasta/ancestral_alleles/homo_sapiens_ancestor_GRCh37_e71.tar.bz2
- human reference genome (hg19) ftp://hgdownload.soe.ucsc.edu/goldenPath/hg19/bigZips/chromFa.tar.gz

Following *hmmix* program documentation, we calculated the average mutation rate in windows of 1 Mb using variants from the outgroup (sub-Saharan Africans).

*hmmix mutation_rate -outgroup={outgroupfile} -weights={weightsfile} -window_size=1000000 -out {mutationratefile}*

We then identified all positions with derived biallelic variants in each non-African SGDP individual:

*hmmix create_ingroup -ind={individuals} -vcf={vcffiles} -weights={weightsfile} -out={observations} -ancestral={ancestralfiles} -outgroup={outgroup}*

For each target individual, we estimated the hmm parameters that maximize the likelihood given the data using:

*hmmix train -obs={observations} -weights={weightsfile} -mutrates={mutationratefile} -out={trainparameters}*

We averaged the trained parameters for each continental group (America, Central Asia and Siberia, East Asia, Oceania, South Asia and West Eurasia) and used these as priors for each individual in that continental group.

We used the four high coverage archaic reference genomes– Altai Neandertal, Vindija 33.19, Chagyrskaya 8, Denisova– to annotate the source of the archaic ancestry in each segment.

*hmmix decode -obs={observations} -weights={weightsfile} -mutrates={mutationratefile} -out={outputfile} -param={trainparameters} -admixpop={admixpopfiles}*

We retained all putative introgressed archaic segments with a posterior probability greater than 0.8 and contained at least one derived variant that is shared with at least one of the high coverage archaic genomes. If segments shared more derived variants with Neandertals than Denisovans, they were classified as Neandertal and vice-versa.

We then calculated the overlap of archaic segments identified by *hmmix* and *admixfrog* across the genome (fig. S9, table S6). By overlap, we mean the number of base pairs that are shared by *admixfrog* and *hmmix* segments, .i.e. we do not require a perfect overlap of coordinates. For Neandertal segments, 63.4% of the total sequence identified by *hmmix* is also identified by *admixfrog*. In turn, 69.2% of the total sequence identified by *admixfrog* is also identified by *hmmix*. For individuals from west Eurasia, *admixfrog* identifies 56 Mb of introgressed Neandertal sequence on average, while *hmmix* identifies 54 Mb on introgressed Neandertal sequence on average. The overlap of introgressed Neandertal sequence is similar to the results for overlap of *hmmix* with other published methods like *CRF* (67%), *Sprime* (84.9%), and *Sstar* (74%) (*78*).

For Denisovan segments, only 16.8% of the total sequence identified by *hmmix* is also identified by *admixfrog*. 81% of the total Denisovan sequence found by *admixfrog* is recovered by *hmmix*. In Oceanians, *admixfrog* only finds 24 Mb of introgressed Denisovan sequence on average, while *hmmix* identifies 70 Mb (fig S9). This is likely because the introgressing Denisovan population is more distantly related to the sequenced Denisovan genome that was used for the design of the admixture capture array (*17*, *23*, *80*). This will in turn limit the amount of introgressing Denisovan sites, which will lower the power of *admixfrog*.

#### 4. Spatiotemporal patterns of Neandertal ancestry

In this section, we compare the segments of Neandertal ancestry inferred using *admixfrog* between all non-African individuals in our data set. We restrict our analysis to the autosomes only.

##### 4.1 Clustering of individuals into genetically informed groups

We assigned all ancient individuals to one of the six SGDP superpopulation based on their sampling location. In addition to the geographical clustering we assigned the individuals into genetically informed groups using variants on the 1240k array. We ran *admixfrog* with the same parameters, but on the 1240k sites to obtain contamination-corrected genotype likelihoods for ancient individuals. For present-day individuals, we used the genotypes directly and for ancient individuals and chose the genotype with the highest likelihood.

We calculated *outgroup*-$f_3$ statistic (*81*) for all pairwise non-African populations (from SGDP) and on an individual basis for ancient individuals in the form of $f_3$*(Mbuti;ind1/pop1,ind2/pop2)* for autosomal SNPs on the 1240k and archaic admixture array. We used the $f_3$ function in ADMIXTOOLS2 (*82*) with precomputed $f_2$ (*extract_*$f_2$ with maxmiss = 1, auto_only = T, adjust_pseudohaploid = FALSE, $F_{st}$=FALSE, apply_corr=FALSE). We remove sites where more than 50% of the ancient individuals have no data.

To infer the dissimilarity matrix from the $f_3$ matrix, we set the diagonal to 0 and subtracted the maximum $f_3$ from each value. We exclude individuals with close genetic relationships from the calculation of the maximum $f_3$ (e.g., the Afanasievo sons) (table S7). We used a hierarchical clustering approach on the distances of the $f_3$-values to assign the individuals into groups using the *hclust* function in *R*. We inspected the clusters and refined them with information from previous studies (*35*). This leads to two sets of clustering depths (table S8): One set of superpopulation clusters consisting of: Americas and Asia (Americas&Asia), Oceania, West-Eurasia and Central Asia and Siberia (WEurCAS), and an Early-Out-of-Africa cluster (EarlyOoA). We refine some of the superpopulation clusters into sub clusters as follows: Americas&Asia into ancient East-Asia (ancientEAS), East-Asia (EAS) and Siberia&Americas (SA), WEuCAS into West-Eurasia (WEur), South-West-Asia (SWA), pre-Last Glacial Maximum West-Eurasian Hunter-Gatherers (preLGM-WEurHG), post-Last Glacial Maximum West-Eurasian Hunter-Gatherers (postLGM-WEurHG), ancient-North-Siberians (ANS), ancient North-Eurasians (ANE), Satsurblia and Yamnaya. We inspect the clustering result using multidimensional scaling on the euclidean distances among all pairwise $f_3$ values (excluding Sub-Saharan-Africans) using the *cmdscale* function in *R* (fig. S10).

##### 4.2 PCA of Neandertal Ancestry

We assessed the structure of Neandertal ancestry by performing a Principal component analysis (PCA) on the matrix of ancestry segments across the autosomal genome of all individuals. We only considered Neandertal segments with minimal genetic length of 0.2 cM for ancient individuals and 0.05 cM for present-day individuals. We calculated the intersection of segments using the *bedtools multiinter* function. We masked out all the regions in our non-callable-region mask. We used the resulting intersect matrix of 0 and 1, where 1 denotes that a given individual has a Neandertal segment at a given region and 0 implies otherwise. We compute a PCA using the *prcomp(.,center=T)* function in *R*. We focused on the top 5 regions with the highest loadings on PC1 and PC2 to check for population-defining segments, i.e. segments that have a high loading on the PC's and are therefore highly informative to separate individuals and populations (fig. S11).

##### 4.3 Correlation of Neandertal Segments

In this section, we want to compare the sharing of the Neandertal introgressed segments with the overall population structure. If a majority of Neandertal ancestry is due to gene flow from one Neandertal population into the common ancestors of all individuals in our study, we would expect the introgressed Neandertal segments to mirror the population structure at random sites, as they are subject to drift in the same way as other sites in the genome. Therefore, we calculate a genome-wide correlation of shared Neandertal-ancestry segments across individuals. We excluded individuals with close genetic relatedness ("AfanasievoSon1","AfanasievoSon2"). The data is based on a matrix X whose entry $x_{ij}$ is the *admixfrog* posterior probability that the bin j in individual i having heterozygous or homozygous Neandertal ancestry. Bins outside a called Neandertal segment with minimum length of 0.2 and 0.05 cM for ancient and present-day individuals respectively, are set to zero while all others are set to 1. We removed bins that do not overlap the callability-mask. We calculated the correlation using the *cor* function in the package base in *R* (table S10). We tested if the EarlyOoA cluster has the lowest mean correlation using a one sided t-test using the *t.test* function in *R* with the option alternative = "less". We corrected the p-values for multiple testing using the Holm-Bonferroni method.

To contrast the correlation of Neandertal segments to the overall population structure, we calculated the *outgroup-$f_3$* statistic for all pairwise non-African individuals *(ind1*, *ind2*) in the form of $f_3$(Mbuti;*ind1*,*ind2*) as described in Materials and Methods Section 4.1 for archaic admixture + X and the 1240 k sites (table S7 and S9).

We standardize the $f_3$ matrix and the correlation matrix by subtracting the population mean from each value and dividing by the standard deviation for plotting it. We excluded individuals with close genetic relatedness from the calculation of the population mean and standard deviation (specifically, we excluded the Afanasievo sons). We analyzed the relation between the two non-standardized statistics using a simple linear regression. We also did a linear regression for the $f_3$-values calculated using the 1240 k ascertainment and the archaic admixture + X (fig. S12, table S7 and S9). There is a significant correlation between Neandertal segment correlation and outgroup-$f_3$ on 1240 k sites (Pearson correlation R= 0.78, p < 2.2e-16), between Neandertal segment correlation and outgroup-$f_3$ on archaic admixture sites (Pearson correlation R= 0.81, p < 2.2e-16) and between the two $f_3$ estimates (Pearson correlation R= 0.88, p < 2.2e-16).

##### 4.4 Identification of unique introgressed segments

In this section, we identified the amount of introgressed segments that is unique to one of the genetically informed clusters, or uniquely shared between *n* clusters. First, we computed the intersection of segments for all individuals in one segment cluster. To do this, we first merged all individuals in a group by taking the minimum start and maximum end of the union of overlapping segments. Then, we calculated the intersection using *bedtools multiinter* function to all other groups. We masked out all the non-callable regions using our callable-region mask (fig. S1) and computed the proportion of Neandertal ancestry (in bp) as the ratio of the total amount of the callable genome that is unique to the cluster or shared with one or more other clusters (fig. S13A, table S11). We excluded individuals with close genetic relatedness ("AfanasievoSon1","AfanasievoSon2").

To identify how much unique Neandertal ancestry can be retrieved from a single individual per cluster, we randomly sampled one individual per cluster and intersected these individuals by the segment start and end positions and masked out non-callable regions. We obtained the mean and standard deviation by repeating the process by randomly sampling an individual per group and calculating the intersection 100 times (Main Figure 2C, table S12). To assess if there is any group size or coverage bias, we correlated the amount of unique segments per group to the number of individuals in the group and the average proportion of callable genome in the individuals from that particular group. We repeated this analysis for the EarlyOoA cluster by leaving out Oase1 that is a clear outlier for Neandertal ancestry (and does not contribute to present-day individuals). We neither found a significant correlation between number of individuals and the unique Neandertal ancestry (Pearson correlation p-value = 0.43) nor the average coverage and average proportion of callable genome in the samples of any particular group of (Pearson correlation p-value = 0.12) (fig. S13B and S13C). To assess if there are any significant differences in the amount of segments unique to one group compared to all other groups, we used a pairwise t-test with Holm-Bonferroni multiple testing correction implemented in the *pairwise.t.test* function in *R* (table S13).

##### 4.5 Archaic lineage assignment of introgressed segments

We used diagnostic sites between the SGDP B_Mbuti-4, the Altai, Vindija and Chagyrskaya Neandertals as well as the Denisovan to identify the closest reference individual for any target non-African individual (all diagnostic sites ascertainment see Materials and Methods Section 2.5). For each ancient target individual, we used a pileup of all reads overlapping the diagnostic positions in regions called as Neandertal ancestry. We only considered reads with a minimum mapping quality of 25 and a minimum length of 35 bp. For this analysis, we extend the sites from the archaic admixture + ArchaicX array to all diagnostic sites present on Neandertal segments. We used two approaches to calculate the matching positions between the admixture segment and the reference archaic genomes.

1. We randomly sampled an allele for every covered position on the segment and calculated the matching of the segment to all reference individuals as the proportion of shared derived alleles, divided by the number of covered sites on the segment.
2. For every covered position, we counted how many reads overlap the derived allele and divide by the total amount of reads overlapping a segment. We evaluated both methods of estimation and found no differences (see Materials and Methods Section 4.5.1).

We continued using estimates from randomly sampled alleles. For all analysis we only looked at the autosomes. If segments are overlapping (sharing the same genetic location) in a cluster we randomly sampled one segment to obtain a set of non-overlapping segments.

##### 4.5.1 Match rate distribution

We plotted the density of the matching of the non-overlapping segments per cluster to a single sequenced Neandertal (Chagyrskaya and Vindija)  and also contrasted two sequenced Neandertals (Vindija versus the Altai Neandertal) for captured and shotgun data per population cluster (Main Figure 2D, fig. S14 and 15). We note that there is a shift in the density related to the data type used (shotgun vs. capture) and that we found very noisy estimates for the post-LGM Oceania cluster which only consists of the individual from Leang Panning which is an outlier in contamination and coverage (see Materials and Methods Section 3.1). We therefore excluded the post-LGM Oceania cluster from further analysis. Furthermore, we recorded the overall closest reference (MButi, Denisova, Vindija, Altai Chagyrskaya) to all segments harbored by an individual. We tested if one or two normal distributions fit the match rate to the Chagyrskaya Neandertal better. We chose this Neandertal since it was not part of the archaic admixture ascertainment, on which some sampled individuals were captured with and should therefore be unbiased. We followed (*23*) and only considered segments with a match rate to the Chagyrskaya Neandertal of 0.3 or higher. Furthermore, we required every segment to have at least 15 informative positions for the match to Chagyrskaya (i.e. in the genomic region of the segment the Chagyrskaya Neandertal has at least 15 positions that are included in the all diagnostic sites ascertainment). We fitted the single normal distribution using the *optim* function in *R*. For the mixture of two normal distributions we implemented an Expectation Maximization (EM) algorithm. The initial starting parameters for $\mu$, $\sigma$ and the mixture proportion $w$ were 0.3 and 0.7, 0.1 and 0.1 and 50/50, respectively. To prevent numerical underflow in the EM we set the minimum value of each parameter to 1e-12. We compared the log-likelihoods using a likelihood ratio test (LRT) the corresponding p values were estimated from a chi square distribution with one degree of freedom. We did the test for all clusters and data type (shotgun vs. capture) independently. The results are in table S14. We found that for capture sequenced individuals of the EarlyOoA cluster, namely BKF6620, BKCC7335, BKBB7240 and Oase1, we find an additional normal distribution fits significantly better (p-value < 0.0001) with 6 % of the segments explained by a distribution centered at a matchrate of 0.51 and the rest with a distribution centered around 0.78.

##### 4.5.2 Proportions of Segments matching the reference archaic individuals

Additionally we looked at the proportion of segments which uniquely match best one of the reference archaic individuals. Therefore, each segment is assigned to the highest matching reference individual. It is possible that two or more references match equally well and that this matching is discordant to the overall population tree. For each population we visualized the proportion of non-overlapping segments matching a particular archaic reference from all individuals in a population cluster. We only considered segments that match uniquely to an archaic reference (fig. S16). We statistically test if the proportion of segments between population clusters is significantly different while taking the data type (shotgun vs. capture) and the set of sites chosen (all diagnostic sites vs.only sites that also are found in the archaic admixture array). We test this for superpopulation and population clusters independently. To explicitly model the proportion with multiple predictors, we use a Bayesian generalized mixed model with a multinomial likelihood distribution as our maximum entropy distribution. To gain computational efficiency, we realized the multinomial likelihood as a series of Poisson distributions (eq. 1). The outcome variable *Y* is the proportion of assigned ancestry for each segment. We used random intercepts for each population cluster and predictor variables for the data type *D* (shotgun or capture) and sites *S* (all diagnostic sites vs all diagnostic sites found also in the archaic admixture ascertainment). We denote *Y* as the outcome variable, *pop* as an indicator variable and *D* and *S* as binary predictors. We obtained the posterior probability using a Hamiltonian Monte Carlo MCMC algorithm, as implemented in STAN (*83*) using R (*84*, *85*).

We tested the power of the model of proportions using simulated data. To this end, we simulated a data set of 100 individuals from 4 groups with a total 10,000 segments in *R*. The number of segments per individual and the membership of an individual in a group is randomly assigned. The ancestry of the segment is sampled from 4 possible ancestries with probabilities given by the *softmax* function taking scores for each of the 4 possible ancestries. These scores can be modulated with the effect variables of group membership, data type (binary variable which is randomly assigned to an individual), coverage (continuous variable drawn from a gamma distribution with shape 1 and rate 1/4) . The baseline probabilities to be from each of the 4 ancestries was chosen to be equal (0.25). We simulated four data sets. One with no differences between the groups, one with differences between groups caused by different group membership, one with differences caused by groups with different proportions of individuals having different data types and the last with differences caused by groups with different proportions of individuals having different data types and coverages. The estimated difference between the ancestry proportions of each population against a pivot population for the simulation is depicted in fig. S17.

For the real data, we ran 6 Markov chains with 1000 warm-up steps and 1000 samping steps each. The Markov chains converged to the target distribution (Rhat = 1) and efficiently sampled from the posterior (table S15).

$$y_{i,k} \sim Multinomial(n, p_1, .., p_n) \qquad \text{(eq. 1)}$$
$$p_k = q_k/\sigma$$
$$q_k \sim Poisson(\lambda_k)$$
$$\sigma = \sum_k^n \lambda_k$$

$$log(\lambda_k) = \alpha_{pop[i]} + \beta_{DT}D_i + \beta_s S_i$$
$$\alpha_{pop[i]} \sim Normal(0, 1.5)$$
$$\beta_{DT} \sim Normal(0, 2)$$
$$\beta_S \sim Normal(0, 2)$$

From this model, we obtain posterior predictions for the proportion of segments best matching any of the four sequenced archaic individuals for each target population, conditioned on data type and sites chosen. This can be thought of as the proportion of segments coming from a certain archaic ancestry corrected by data type and sites used. We compared the differences between the predicted proportions by subtracting each population predicted proportion from a pivot population. We chose WEurCAS as the pivot for comparisons of superclusters and West-Eurasians for population clusters (fig. S18, table S16). The EarlyOoA cluster shows in comparison to the superpopulation and population cluster an increase in the proportion of segments closer to the Altai Neandertal and a decrease of segments close to Chagyrskaya, as was also evident in the match rate distribution analysis. The ANE cluster, consisting of the Mal'ta and Afontova Gora 3 individuals, is the only additional cluster that shows a difference compared to the pivot populations (in this case it has more segments closer to the Vindija Neandertal). We however, don't think that their Neandertal ancestry is really different since we only have two samples and the descendant populations of this cluster (Siberians&Americans (*55*)) don't show such a signal.

##### 4.5.3 Validation of lineage assignment

To gauge the effect of different data types (shotgun vs. captured data) and ways of estimating the matching (proportion of derived reads or random sampling of an allele), we used the Kostenki14 individual which was shotgun sequenced and captured on the archaic-admixture array (*35*, *52*). Fig. S19 shows the overall proportion of matching to a specific reference is proportionally reduced between the captured data compared to the shotgun (shift on the diagonal) but not biased (no shift off the diagonal). We checked the assignment of segments to a single reference (Altai Neandertal, Chagyrskaya Neandertal, Vindija Neandertal and Denisova). Pearson's Chi–square test with two degrees of freedom did not reveal a significant dependence between either segment assignment composition and data type (p-value = 1) nor way of estimation (p-value = 0.23). Nevertheless, we also modeled the probability of the assignment with a Bayesian generalized mixed model in the following form (eq. 2):

$$y_{i,k} \sim Multinomial(n, p_1, .., p_n) \quad \text{(eq. 2)}$$
$$p_k = q_k/\sigma$$
$$q_k \sim Poisson(\lambda_k)$$
$$\sigma = \sum_k^n \lambda_k$$
$$log(\lambda_k) = \alpha_{ind} Ind_i + \beta_{method} M_i + \beta_{DT} D_i$$
$$\alpha_{ind} \sim Normal(0, 1.5)$$
$$\beta_{method} \sim Normal(0, 2)$$
$$\beta_{DT} \sim Normal(0, 2)$$

Compared to the shotgun data, there is an estimated slight 0.09 probability decrease (95% HPDI: 0–0.18%) in assigning segments to the Altai Neandertal when using captured data. There is no effect on the assignment based on the method used (fig. S20).

#### 5. Genetic Dating of Neandertal Gene flow

The distribution and length of archaic ancestry segments can be used to estimate parameters of gene flow events (*86*). The mean time and duration of the gene flow between Neandertals and modern humans was estimated using present-day individuals (*28*, *29*). This can either be done using inferred segments directly, or using Linkage Disequilibrium (LD) at SNPs ascertained for archaic ancestry. The two statistics are equivalent, but LD has the advantage that it integrates out the uncertainty in inferred fragments (*29*). Recently, Moorjani et al. extended the LD based method to single ancient individuals by measuring the extent of covariance between Neandertal introgressed sites along the genome resulting in ancestry covariance curves that can be used for inferring parameters of the gene flow event. Assuming an instantaneous gene flow (IG model) model, we can fit an exponential to the ancestry covariance curves to infer the time of gene flow for a single individual (*19*, *87*). We then examined the correlation in the dates of gene flow (in generation) and sampling age (in years) and estimated the historical generation time interval and the joint date of Neandertal admixture across all ancient individuals. We further evaluated an approach for jointly fitting the IG model and an extended admixture pulse model (EP model) (*29*) using both the called Neandertal segments and the ancestry covariance curves. We applied the joint fitting of the IG model for segments and ancestry covariance curves and the EP model for coancestry curves only to our data.

##### 5.1 Ancestry covariance curves

To obtain ancestry covariance curves, we used the ACov ascertainment (see Materials and Methods Section 2.6) containing a set of 227,346 markers across the genome that match the ascertainment scheme that is informative for Neandertal introgression [i.e., sites where Neandertals carry at least one derived allele (relative to chimpanzees) and all individuals in a panel of Sub-Saharan Africans carry the ancestral allele]. For each early modern human individual, we inferred the genotypes for the ascertained markers based on the genotype likelihoods estimated by *admixfrog*. Following (*19*), we estimate the covariance across pairs of ancestry-informative markers separated by increasing genetic distances across the genome using the Shared genetic map. Assuming an instantaneous gene flow model, this statistic is expected to decay approximately exponentially with genetic distance, and the rate of decay is informative of the time of mixture. It was shown in (*19*) using simulations that this statistic works reliably for single ancient individuals that are older than 10,000 years, even with low coverage (~1x).

##### 5.2 Single-sample dating

We inferred the timing of the Neandertal admixture for 22 early modern human genomes with radiocarbon ages ranging between ~24,000–48,000 years (table S2). We obtained significant estimates for the dates of admixture, regardless of the genome coverage, age of the individuals or number of non-missing genotypes for the ascertained SNPs ($|Z| > 2$) (table S17). For individuals older than 40,000 years, we found that ancestry covariance extended to large genetic distances (>1cM) as shown in fig. S21 and so we ran our analysis to longer genetic distances until the intercept of the exponential decay was close to 0 (e.g., 10–60cM as needed).

##### 5.3 Generation Time estimate

We leverage the linear relationship between the dates of Neandertal admixture and sampling age (Fig. 3A) to jointly estimate the generation interval (reflected by the slope) and the time of the shared pulse of Neandertal admixture in modern humans (reflected by the intercept). To do this, we used a Bayesian model to account for the uncertainty in both the dates of Neandertal admixture and radiocarbon dates estimated using importance sampling as described in (*19*). We tested if our generation interval estimates are robust to the exclusion of very early modern humans like Zlatý kůň, Ust' Ishim, Oase or Bacho Kiro individuals that have previously been inferred to have gene flow from multiple pulses of Neandertal introgression (*7–9*, *19*) (table S18).

##### 5.4 Assessment of instantaneous gene flow (IG) vs. extended admixture pulse (EP) model

In addition to the single sample statistics for admixture dating, we jointly fitted all individuals older than 20 kyBP (22 individuals) using a model of instantaneous one generation long gene flow (IG) or an extended pulse model (EP). For the joint fitting of all individuals together, we have to assume that no major gene flow happened after the sampling time of the oldest individual. This might be violated by the individuals of the EarlyOoA. We investigated the reliability of ancestry covariance and the ancestry segments inferred by *admixfrog* for fitting the models using simulations under each model.

###### 5.4.1 Simulated ancestry segments

We simulate 150 segment lengths ($l$) from the analytical distributions per individual, taking the difference in sampling time into account. Specifically, for the IG model we sampled admixture segment lengths from an exponential distribution with parameter $t_m - s_t$, where $t_m$ is the admixture time, and $s_t$ is the sample time, using the *rexp* function from the *stats* package in *R*. For the two-pulse model, we used a 50/50 mixture of two exponential distributions, with parameters $t_{m1}$ - $s_t$ *and* $t_{m2}$ - $s_t$, where $t_{m1}$ and $t_{m2}$ are the dates of the two pulses. For the extended pulse model, we simulated segments using a Lomax distribution as implemented in the *rlomax* function from the *VGAM* package in *R,* with scale $k/(t_m - s_t)$ and $shape=(k+1)$ where the duration is given by $t_d = 4(t_m - s_t)\, k^{-1/2}$ (*29*). All times are measured in generations, using a generation time of 28 years.

###### 5.4.2 Simulated Ancestry Covariance

To mimic ancestry covariance data, we use the scaled distribution functions of the respective distribution (equation 3a-c) with the distance $d$ from 0 to 0.1cM at intervals of 0.00005 cM for each simulated individual.

###### 5.4.3 Adding Noise

In both cases we added Gaussian noise to the segment length or covariance data of 0, 0.002 or 0.004. For segments we used a lower cutoff of 0.2 cM and for the ancestry covariance data a minimum of 0.0001 and a maximum of 0.015 cM. In equations below for the ancestry covariance (ACov) curves, the values for the constant *c* that models background covariance and the amplitude $A$ or $A1$ and $A2$ were sampled from a uniform distribution between 0 and 0.002 for *c*, 0.02 and 0.04 for $A$ and $A1$ and 0.01 and 0.02 for $A2$ for each of the 50 simulation runs.

$$AC \sim A\, e^{-(t_m - s_t d)}\, c \quad \text{(eq. 3. a)}$$

$$AC \sim c\, A_1\, e^{-(t_{m1} - s_t d)} + A_2\, e^{-(t_{m2} - s_t d)} + \quad \text{(eq. 3. b)}$$

$$AC \sim A\left(1 + \frac{(t_m - s_t)}{k} d\right)^{-k} c \quad \text{(eq. 3. c)}$$

We fitted both the IG or EP model to each simulation, as described in Materials and Methods Section 5.5.4.1 and 5.5.4.2 using equation 4 and 6. The simulation results are summarized in fig. S22. Panel A and B show the estimates using the joint IG model for scenarios of a single, two and extended pulses of gene flow either for ancestry covariance curves or segments under the three noise levels. There are only minor differences between ancestry covariance estimates and segments except for the two pulse model, where segments only capture the date for the recent pulse whereas the coancestry estimates infer the mean time of the two pulses. Moreover, using segments seems to underestimate the mean time of the extended pulse and the effect is exacerbated for longer durations. Panel C and D show the estimates using the joint extended pulse model for scenarios of an extended pulse with varying durations, again for ancestry covariance curves or segments under the three noise levels. We find that both the segments and ancestry covariance curves provide unbiased results, but the estimates based on segments are much noisier. Together, with previous results that showed that inferred segments give downward biased estimates in coalescent simulations (*29*), we only used the ancestry covariance curves for the inference of the extended admixture pulse model.

##### 5.4.4 Application to real data

We estimated a joint instantaneous generation pulse model (IG model) and a joint extended pulse model (EP model) for the timing of Neandertal gene flow using the 22 individuals older than 20 kyBP.

For ancestry covariance curves, we only consider ancestry covariances between SNPs that located at genetic distance (*d*) of 0.02 cM– 1 cM, except for BK1653, Zlatý kůň, Ust'-Ishim, BKBB7240, BKF6620, BKCC7335 and Oase1 where we consider longer distances. For these individuals the maximum length ($d_{max}$) was either set to 60 cM for Oase1 and 10 for BK1653, Zlatý kůň, Ust'-Ishim, BKBB7240, BKF6620 and BKCC733 (so that inferred covariance curves are not truncated) or we identified at which cM value the estimates first are 0 and rounded the cM value up to the next bigger cM. This resulted in a set of maximum distances of 2, 3, 5, 4, 5, 5 and 27 cM for BK1653, Zlatý kůň, Ust'-Ishim, BKBB7240, BKF6620, BKCC7335 and Oase1 respectively. This set is further referred to as "estimated truncation".

For all estimates we subsampled our data into:

- All individuals older than 20 kyBP (n = 22)

- Only individuals from the Early out of Africa cluster (n = 6)
- Early out of Africa cluster removed (n = 16)
- All above and leaving out the Oase1 individual. This individual has evidence for very recent Neandertal gene flow (less than 10 generations) and thus the ancestry covariance curves or distribution of Neandertal segments violate the assumptions of independent and identically exponential distributed segments.

**5.4.4.1 Fitting an Instantaneous gene flow model**

We fit a IG model using both the ancestry covariance and the segments inferred by admixfrog. We took the difference in sample time into account by defining an offset ($s_t$) for each individual given by its mean most likely date (table S2). To convert the years into generations, we use 28 years per generation.

For the ancestry covariance, we used equation 4a. The amplitude parameter *A* and x-intercept parameter *c* are fitted for each individual while $t_m$ is a joint parameter. Fitting was done using the "port" algorithm implemented in the *nls* function from the *stats* package in R, with the upper and lower limits for *c* and *A* between 0 and 1 and for $t_m$ between 1 and 5000. Since the amount of data is determined by maximum length between SNPs considered ($d_{max}$), which varies substantially between individuals (e.g. not truncated Oase1 has 60 times more data points computed than most individuals), we included a weighting parameter *w* which is the ratio between the smallest maximum distance of any individual in the data set and the maximum distance of the individual being considered. The weighting parameter ensures that all individuals have the same weight on the log-likelihood. If we wanted the weight to be proportional to the amount of data points for each individual we simply put *w = 1.* To convert the residual-sum-of-squares (RSS) from the ancestry covariance fit, we used equation 5 with *n* being the degrees of freedom.

$$AC \sim \left[A_{ind}\, e^{-(t_m - s_{t_{ind}}\, d)}\, c_{ind}\right] w_{ind} \quad (4.a)$$

$$P(L_{i,ind} = l) = \left(t_m - s_{t_{ind}}\right) e^{-(t_m - s_{t_{ind}})\, l} \quad (4.b)$$

$$LL \propto -\frac{n}{2}\, log(RSS) - \frac{n}{2} log(\frac{2\pi}{n}) - \frac{n}{2} \quad (5)$$

We obtained the log-likelihoods for the segments by fitting the truncated version of equation 4b as described in (*29*) with the *dexp* and *pexp* functions form the *stats* package in *R* with rate parameter being $\lambda = t_m - s_{t_{ind}}$. The optimization of the sum of the log-likelihoods was done using the *optim* function from the same package. We used three different lower length cutoffs of 0.05, 0.1 and 0.2 cM and an upper cutoff being the longest segment per individual. The results for the estimate of the mean time using the IG model fit on the ancestry covariance curves and segments are displayed in fig. S23 and in table S19 and S20 for point estimates. To estimate the uncertainty we did a Jackknife by leaving each chromosome out one at a time and estimated the weighted Jackknife mean and standard error.

**5.4.4.2 Fitting the Extended pulse model**

In addition to the IG model, we also fitted an extended pulse model which fits a lomax distribution with two parameters, one for the mean time of admixture ($t_m$) and an additional parameter ($k$) for the duration of the gene flow $t_d$, where $t_d = 4(t_m - s_t)\ k^{-½}$ and $s_t$ the offset of the sample time of an individual again. We estimated the log-likelihood for a grid of all possible combinations of pulses with durations ($t_d$) ranging from 1–2,000 generations and mean admixture ($t_m$) times starting from 1 generation before the oldest most likely date of all individuals to 500 generations in steps of 1 generation. The amplitude $A$ and x-intercept $c$ per individual were fitted for each combination of $t_m$ and $t_d$. The fitting was done similar to the IG model using equations 6 a and b. Based on the simulation results, we restricted the analysis of the real data to only ancestry covariance estimates.

$$AC \sim \left[ A_{ind} \left( 1 + \frac{\left(t_m - s_{t_{ind}}\right)}{k} d \right)^{-k} c_{ind} \right] w_{ind} \quad (6.\ a)$$

$$P(L_{i,ind} = l) = \left(t_m - s_{t_{ind}}\right)^{\frac{(k+1)}{k}} \left( 1 + \frac{\left(t_m - s_{t_{ind}}\right)}{k} l \right)^{-(k+2)} \quad (6.\ b)$$

The log-likelihood surfaces are shown in the fig. S24. The summary of the extended pulse fitted to the grid of possible values for $t_m$ and $t_d$ is given in table S21. Uncertainty is given by the minimum and maximum values for $t_m$ and $t_d$ for all estimates with a smaller log-likelihood difference to the maximum log-likelihood of 2 (the fits are not significantly different from one another). We notice that the estimates are stable over the weighting and truncation parameter, except when including the Oase1 individual. Furthermore, excluding the EarlyOoA individuals leads to a longer duration of the gene flow. The joint fit of the EP model assumes all major gene flow (more specifically the mean time of gene flow) occurring before the sampling time of the oldest individual. Which might be violated for individuals very close to the gene flow event (namely individuals from the EarlyOoA cluster). Therefore, we fitted the EP model to every EarlyOoA individual independently (fig. S25). The estimates on a single sample are noisy. However, some individuals are estimated to still have major gene flow after the sample time of other individuals. This can either mean i) the radiocarbon dates are not correct, ii) some individuals have private gene flow events or iii) individuals are sampled during the major pulse, or a combination of the three. For the estimate of the major gene flow event we therefore resolve to estimate the EP model on the 16 individuals sampled after 40,000 years. The model prediction of the migration rate over time is given in fig. S26.

###### 5.4.4.3 Comparing IG and EP model

We compared the models of the instantaneous one generation pulse fit to the ancestry covariance curves with that of the extended pulse using a likelihood ratio test. Additionally we used empirical significance cutoffs from Kozubowski et al. 2008 and required a log-likelihood difference equal or bigger than 2.5 for a significance of $\alpha = 0.05$. The results are given in table S22.

#### 6. Neandertal Ancestry across the genome

In this section, we looked into the amount of Neandertal ancestry in each individual, how it is distributed across the genome and how the frequency has changed over time. To obtain frequency estimates, we grouped our data into 4 different time intervals: non-African present-day individuals (n=231), ancient individuals younger than 10 kyBP (25), ancient individuals between 10 and 30 kyBP (n=13) and ancient individuals older than 30 kyBP (n=20). For examining the changes in frequency trajectories, we only considered individuals sampled from West Eurasia, Central Asia and Siberia to minimize the effects of population structure impacting the results.

##### 6.1 Total recovered Neandertal genome and total deserts

We calculated the total amount of Neandertal introgressed material across individuals by aggregating all overlapping segments that are longer or equal than 0.05 cM in present-day individuals and 0.2 cM in ancient individuals. We then took the union of the overlapping segments and removed the segments not included in our callable genome mask.

For deserts, we took the union of segments inferred to be homozygous African by *admixfrog*. We applied the same length and coverage cutoffs as for calling Neandertal segments. We removed all called Archaic regions in our individuals with a minimum length of 0.01 cM (two consecutive *admixfrog* bins). We annotated regions that have significantly reduced archaic ancestry in present-day individuals, previously referred as 'archaic deserts' (*13*, *15*). The result is depicted in fig. S27.

In total, we recovered 1,551 Mb of haploid Neandertal genome which amounts to 61.7 % of the callable genome. We also calculated the frequency of Neandertal ancestry throughout the genome. Fig. S28 shows the result on a subset of our data that includes all ancient individuals and two SGDP individuals per genetic cluster.

##### 6.2 Neandertal frequency correlation

Previous studies have shown that Neandertal ancestry varies across the genome and is correlated to several factors such as B-statistic (a measure of linked selection) and recombination rate in present-day individuals (*12*, *38*). Below we investigate the correlation between Neandertal ancestry and each of these factors at four time points.

###### 6.2.1 B statistics

We investigated the relationship between the amount of introgressed Neandertal ancestry with B-statistics across the callable genome (*39*). The B-scores are distributed from 0 to 1000, whereas a low B score indicates evolutionary constrained regions. We binned the B-scores in 5 windows (0-200, 201-400, 401-600, 601-800, 801-1000) and calculated the average amount of basepairs covered by Neandertal ancestry for all genomic windows in each of the B-score bins. Fig. S29 shows the results for the four time intervals. We observe a similar pattern as (*12*) where we find lower amounts of Neandertal ancestry in evolutionary constrained regions in present-day individuals. Notably, we also find this pattern in ancient individuals across all time intervals, even for individuals that are closest to the admixture event. For instance, EarlyOoA individuals harbor a higher average amount of Neandertal ancestry (~6%) but it is nearly half in constrained genomic regions. This suggests that the depletion in evolutionary constrained regions occurred soon after the gene flow as predicted by theoretical models (*18*).

###### 6.2.2 Recombination rate

We computed the per base pair amount of Neandertal ancestry for all non-African individuals and for individuals in the time intervals separately in 20, 50, 100, 200, 500 kb and 1 MB windows across the callable genome and correlated it with the recombination rate in that window based on the Shared map. We used Spearman's rank test as implemented in *R*. We also computed the correlation again using a lower cutoff for the segments of 0.01 cM for all individuals. We find significant correlation between recombination rate and the amount of Neandertal ancestry for all window sizes and time intervals for all individuals younger than 30 ky. For individuals older than 30 ky, we only find a significant correlation for window size larger than 100 kb. For six EarlyOoA individuals, we infer either a negative correlation or non-significant values indicating that we have limited power (*38*). Our results are robust to changing the minimum length cutoff for archaic segments from 0.2 cM for ancient and 0.05 cM for present-day individuals to 0.01 cM for all individuals (table S23).

##### 6.3 Analysis of high frequency regions

To identify candidates of archaic adaptive introgression, we used an outlier approach where we identified regions with high frequency of Neandertal ancestry, compared to the genome-wide average frequency. We grouped individuals into two groups, present-day and ancient individuals. To minimize the impact of population stratification, we focused on individuals from Eurasia including Europeans, Central Asians and Siberians (table S1 SGDP_Superpopulation "WEurAs" and "CeAsSi") as they have the largest sample size. In total, we have data for 45 ancient Eurasian and 101 present-day populations from SGDP. We performed the analysis on a set of Neandertal ancestry calls in windows. We used the bins defined by *admixfrog* (0.005 cM long) as windows. We used bins where i) all bins are on a Neandertal segment called by *admixfrog* to be equal or longer than 0.2 cM in ancient and 0.05 cM in present day individuals and ii) all of these bins have a posterior probability of Neandertal ancestry that is equal to or higher than 0.8.

###### 6.3.1 Outlier estimation

We estimated the average posterior probability of Neandertal ancestry in ancient and present-day individuals separately. By considering the distribution across the genome, we inferred a *Z*-score for each window that measures the deviation from the average genome-wide Neandertal ancestry accounting for the variation across the genome. Then, we identified windows across the genome which are enriched for archaic ancestry (99.9th percentile tail or Z-score > 4.5).
Using these data, we searched three types of regions, i) candidate regions that are immediately adaptive with high frequencies (Z > 4.5) in ancient and present-day individuals, ii) candidate regions with selection on standing variation that are at high frequency in present-day individuals (Z>4.5) but not in ancient individuals (Z < 1), and iii) candidate regions that were adaptive soon after the gene flow and thus are at high frequency for the ancient individuals (Z > 4.5) but are no longer at high frequency in present-day individuals (Z < 1).

To this end, we constructed a matrix of genomic bins defined by *admixfrog* (0.005 cM long) as our windows in columns and individuals in rows with the values in each cell reflecting the posterior probability of Neandertal ancestry inferred by *admixfrog*. We only considered bins overlapping a Neandertal segment longer than 0.2 cM for ancient and 0.05 cM for present-day individuals and additionally all bins to have a posterior probability of Neandertal ancestry equal or higher 0.8. Archaic regions outside these thresholds were filtered out and the correspondingly these bin values were set to 0.

We divided the individuals into two groups: ancient and present-day individuals. For each group, we estimated the mean archaic ancestry per window using the posterior probabilities. We then estimated the genome-wide distribution of the archaic ancestry and computed the mean and standard deviation. We used these estimates to compute a Z-score per window using equation 7 for both cohorts of ancient and present-day individuals separately.

$$z = \frac{(\mu 1 - \bar{u})}{\bar{\sigma}} \qquad \text{eq(7)}$$

We first merged consecutive identified bins into regions for each of the three types of region categories we defined. We merge these regions again if they are 0.05 cM or less apart from each other. In total we identified 209 regions with 143 being equal or longer than 0.05 cM. The mean length of these regions is 0.12 cM or 134 Kb. We annotated these regions using Human Gene annotation from Gencode (release 44, GRCh37), Havana and identified 614 genes in total (table S24).

We found 86 candidate regions (65 >= 0.05 cM) that are immediately adaptive with high frequencies in ancient and present-day individuals with 347 genes and an average length of 0.16 cM or 211 Kb.

We identified 91 (56 >= 0.05 cM) candidate regions with selection on standing variation that are at high frequency in present-day individuals but not in ancient individuals with an average length of 0.07 cM or 76 Kb. These regions contain 169 genes

Finally, we found 32 (22 >= 0.05 cM) candidate regions that were adaptive soon after the gene flow and thus are at high frequency for the ancient individuals but are no longer at high frequency in present-day individuals with an average length of 0.16 cM or 96 Kb. These regions contain 102 genes. From visual inspection, we saw that regions that decrease in frequency over time are close to candidate regions of immediate selection. We estimated that 44% of these regions are 1 Mb up or downstream of candidate regions of immediate selection. The observed values were outside of the empirical distribution, which we estimated using the *ecdf* function in *R* from randomly shuffling the start and end coordinates of high frequency immediate or standing variation candidate regions throughout the genome and re-estimating how many are in 1 MB up or downstream 1000 times. We performed the same test for proximity of candidate regions of immediate and standing variation (8 %, empirical p-value > 0.24) and for the proximity of regions decreasing between ancient and present day individuals to those candidate regions of selection on standing variation (9 %, empirical p-value > 0.15).

We ranked the regions for each type using the Z-scores. For candidate regions of immediate selection, we ranked them by the highest mean Z-score between the ancient and present-day cohort. For candidate regions of selection on standing variation, we ranked them by highest difference between the Z-score for the present-day and ancient cohort. Finally, we ranked the regions that decreased over time by the highest difference between the Z-score of the ancient and present-day cohort.

##### 6.3.2 Comparison to previously identified regions

We assessed the trajectory of Neandertal frequency at the candidate regions of adaptive introgression previously identified in present-day individuals (*12*, *13*, *15*, *89*). Following (*37*), we selected 42 non-overlapping Neandertal regions on the autosomes found in present-day European populations from the list of (*89*). We only considered regions that were found in at least two studies and arrived at 12 regions, from which 11 were callable in this study (region chr20:62160001-62200000 has no recombination distances assigned in the Shared recombination map we used) (fig S30, table S26). We found that the three regions not categorized with our approach have high Neandertal ancestry in present-day individuals (Z>4.5) but which are neither outliers nor average in the ancient cohort (1<Z<4.5) (fig S31, table S26). We also compared our trajectory classification with (*37*) and found that we agreed on 6 out of 8 occasions (table S26). For a region on chr8:13880001-13920000 we see already high frequency (~ 33%) in the earliest time interval and classified it a candidate region of immediate adaptation in contrast to (*37*). Our classification also disagrees for the region containing BNC2 being suggested to not be immediately adaptive in (*37*). We detect ~22% frequency in our earliest time interval.

##### 6.3.4 Gene Ontology Enrichment

We assessed the functional impact of archaic ancestry by identifying pathways that are enriched for archaic ancestry using Panther DB (Version 18, https://www.pantherdb.org). We looked for enrichment in pathways related to molecular function, biological process and cellular components. We also looked for enrichment in categories related to protein class and reactome pathways. We used Gencode (release 44, GRCh37), Havana build as the reference gene list and the candidate regions identified in our analysis as the target gene list. We performed Fisher's exact test for identifying overrepresented categories and performed multiple testing corrections using the Benjamin-Hochberg test to infer the false discovery rate (FDR) (table S25).

##### 6.3.5 Trajectory of archaic frequency over time

To visualize the trajectory of the candidate regions over time, we divided our dataset into 4 time bins of 50-30k (Paleolithic, individuals = 18 ), 30-10k (Last Glacial Maxima, individuals size=10), 10-0k (Holocene, individuals = 20) and 0k (present-day, individuals = 101). We inferred the frequency of Neandertal ancestry by estimating the number of individuals overlapping the region that have Neandertal ancestry inferred by *admixfrog*. We infer the frequency for each time window separately (Fig. 4B-D,fig S30 and 31).

#### 6.4 Deserts through time

We analyzed in detail the formation of previously identified archaic deserts detected in present-day individuals in our ancient individuals. To do so we took into account both Neandertal and Denisovan ancestry. To this end, we took the union of archaic deserts identified in previous surveys (*13*, *15*). As described in the Materials and Methods Section 6.1, we called homozygous African ancestry with a minimum length of 0.2 cM per segment and archaic ancestry with at least 0.01 cM length for all ancient individuals in the region of the deserts. We removed regions outside of our callable genome mask. We further excluded segments that end or start within 200 kb of the borders of the deserts (in case the deserts from Vernot and Sankararaman overlap, the end and start of the desert is the union of the two). We ought to differentiate between archaic ancestry from processes such as ILS, which is usually found in short segments, and recently introgressed segments. In the previous sections we classified segments to be recently introgressed when they are at least 0.2 cM in ancient and 0.05 cM in present-day individuals. To increase our sensitivity of detecting recently introgressed archaic segments in deserts, we decreased our length cutoff from 0.2 cM to 0.1

cM for ancient individuals. We inspected each segment of potentially introgressed archaic ancestry in the deserts by plotting the frequency of reads matching uniquely derived sites in the Denisovan or Neandertal reference genomes. In total, we found 107 segments which are longer than 0.01 cM which are closer to an archaic reference overlapping the deserts (fig. S32) with 95 of them shorter than 0.1 cM, and thus likely the result of either ILS or false positives (average length 49,301 bp / 0.03 cM). Twelve segments were longer than 0.1 cM and putatively introgressed segments (average length 533,710 bp / 0.51 cM). Among these we found eight segments that show strong support of recent introgression with multiple SNPs unique matching an archaic reference, wharease segments in Afontova Gora 3, Loschbour, ZEVJ31 and Satsurblia only show weaker support (fig. S33). The putatively introgressed segments are found at the edges of the deserts. If we restrict our analysis to the intersection of the deserts previously identified in present-day individuals, we only find two putatively introgressed archaic segments in Loschbour and Afontova Gora 3 with weaker support on chromosome 9 (102200000 - 112300000). In regions which are only identified by Vernot et al. 2016, we find on chromosome 8 (53900000 - 66000000) archaic segment in Yana1 and on chromosome 13 (49000000 - 61000000) WC1 and ZVEJ31, suggesting that these desert likely formed after 310000 years and as recent as 5900 years ago, respectively .

We further compared the average amount of archaic ancestry in the intersection of the deserts to the amount of archaic ancestry in genomic windows of similar size across the genome. If two deserts overlap, we consider the union of the two. We estimated the archaic ancestry in the deserts as the average number of bp covered by all segments assigned either to Neandertal or Denisovan ancestry that is at least 0.01 cM in length, per individual divided by the length of the desert. We did the same for all genomic windows of a size of 15,180,000 bp (which is the average length of deserts) and shifted this window by 1 Mb throughout the callable genome. We excluded windows that overlap the previously annotated deserts. We find that the deserts are in the lower 0.1 percentile of the empirical cumulative distribution of archaic ancestry genome-wide, implying that there are at most 30 out of 516 windows that have lower average archaic ancestry than any of the deserts in all non-African individuals throughout the four time intervals (table S30).

#### 7. Neandertal ancestry on the X chromosome

Previous studies have shown that the X chromosome is substantially depleted in archaic ancestry compared to autosomes (*13*, *16*). Several explanations for this have been proposed– from decreased male fertility of human-neandertal offspring (*13*) to the male-biased introgression of archaic individuals (*90*). In addition there have been repeated selective sweeps on the X chromosome replacing regions of archaic ancestry (*91*) in the ancestral population of present-day humans and Ust'-Ishim. These sweeps are defined as regions of minimum 500 kb where the maximum divergence is less than 0.005% and the swept haplotype is shared by at least 25% of non-Africans. However the timing of the sweeps as well as the distribution of archaic ancestry on the X chromosome in ancient individuals remains elusive.

In this section, we explored the amount of archaic variation on the X chromosome through time and established when the selective sweeps happened. The archaic admixture + X array contains 83,660 SNPs, ranging between positions 61,019 to 155,235,202 bp. When grouped into 20 kb windows, there are 7742 windows with an average of 10.9 SNPs in each 20 kb window. 511 windows contained no SNPs and these noncallable windows span a total of

10,220,000 bp. Thus, we investigated 93% of the X chromosome. Since the Shared recombination map does not contain estimates for the X chromosome, we used the African-American map for all analysis in this section (*32*)).

We assessed the patterns on the X chromosome in 44 ancient and 275 present-day individuals. 15 ancient individuals did not have data for the X chromosome. These were:

Villabruna, AfontovaGora3, BK1653, BKBB7240, BKCC7335, BKF6620, ElMiron, OaseNew, Ostuni1, SalkhitArchAdm, Tianyuan, Vestonice13, Vestonice16, Vestonice43, LeangPanninge.

Of these remaining individuals, there were 144 females with two copies of an X chromosome and 205 males with one copy of an X chromosome yielding a total of 493 haploid X chromosome copies. We kept putative introgressed Neandertal segments with a minimum length of 0.2 cM in ancient individuals and 0.05 cM in present-day individuals.

First, we explore the total amount of Neandertal ancestry on the autosomes and X chromosome from our callable region using the African American map for all non-African individuals. We retrieved 1319 Mb on the autosomes and 29 Mb or 18.7% on the X chromosome. We found that 22.35 Mb or 78% of Neandertal segments are shared with at least one other individual and 6.4 Mb or 22% are unique.

We then looked at the ratio of Neandertal ancestry between the X chromosome and the autosomes for all non-African individuals. We stratified the ratios by sampling age into 4 different time windows. We investigated the proportion of Neandertal ancestry on the X and the autosomes for males and females separately. The proportion on the X chromosome is consistently lower compared to the autosomes throughout time (fig. S34A). The higher autosomal and X chromosome proportion in present-day individuals compared to ancient individuals is due to the lower minimum length cutoffs for Neandertal ancestry in present day (0.05 cM vs. 0.2 cM). We followed (*90*) and plotted the amount of Neandertal ancestry per bp on the autosomes versus X chromosomes in fig. S34B. We find stable ratios of around 0.229-0.408 for females and 0.131-0.147 for males with high variation amongst individuals in all time intervals.

We further investigated the previously identified sweep regions. We find that all except for one of the previously identified sweep regions, there is no Neandertal ancestry in any of the individuals at any of the four time windows (table S31). We note that we find no evidence of Ust'-Ishim having introgressed ancestry in any of the sweep regions in agreement with a previous analysis that used *hmmix* (*91*).

We calculate the frequency of introgressed archaic segments on the X chromosomes excluding sweep regions (5.6 Mb) and non-callable regions (10.2 Mb). We find that there is a significant decrease of archaic ancestry on the X chromosome over time (p value = 4.51e-06). Assuming uniform decrease over time, this translates to 0.013 % of archaic ancestry lost per 1000 years (fig. S35).

We group the individuals by ancestry and region and examine the change in Neandertal ancestry across time and space. Interestingly, we observe that most of the reduction in Neandertal ancestry occurs after the last glacial maximum in West Eurasian individuals (fig. S36). This effect could be related to the increased power to detect archaic segments in EarlyOoA individuals and the fact that they had more recent archaic ancestry.

We plotted the distribution of introgressed Neandertal segments across the X chromosome and found that archaic ancestry is localized to only a few regions (fig. S37). We statistically tested if the distribution on the X chromosome is more ordered compared to the autosomes by calculating Shannon's entropy (*H*), a measure of order in a system where 0 is totally ordered and 1 random, for each 0.005 cM window (using the bins from *admixfrog*) across individuals for all chromosomes. Individuals with an inferred Neandertal segment in the window were assigned a one else zero. We calculated the mean of the entropy estimates for the autosomes (H = 0.11) and chromosomes (H = 0.03). We assessed if the difference is significant using Wilcoxon rank sum exact test as implemented in *wilcox.test(x = H_X,y=H_autosomes, alternative=”less”)* from the *stats* package in *R*.

We further investigated this pattern by correlating the amount of Neandertal ancestry with the B-Statistic of as a measure of background selection (*39*) but could not find any conclusive pattern (fig. S29, table S32). We correlated the local recombination rate with the amount of Neandertal ancestry for different window sizes and minimum segment length cutoff and found that only one significant correlation (table S33). We conclude that we do not have statistical power in this case.

We also investigated if we could replicate previously identified deserts on the X chromosome. There are two previously identified deserts from 62 - 78 MB and 109 - 143 Mb (*13*). We replicate one desert but find 120 putative introgressed archaic segments in the second desert (fig. S38). Of these 5 segments are found in ancient individuals. We find 115 segments which are present in Europeans, South Asians, East Asians and the Americas.

**Supplementary Figures**

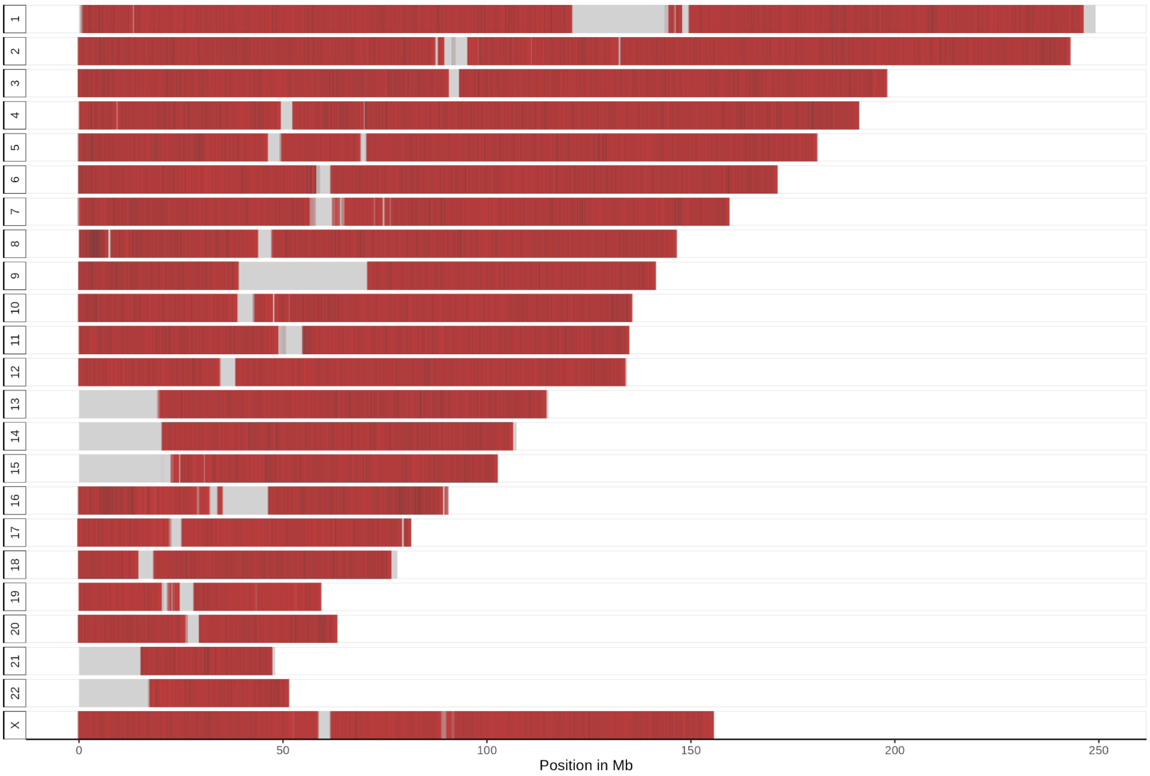

**Fig. S1.** Genomic coverage of the archaic admixture + X sites throughout the human genome. Red bars indicate SNPs included in our analysis, gray background represents the inaccessible regions . Positions on the X-axis (in hg19 coordinates) are given in Mb.

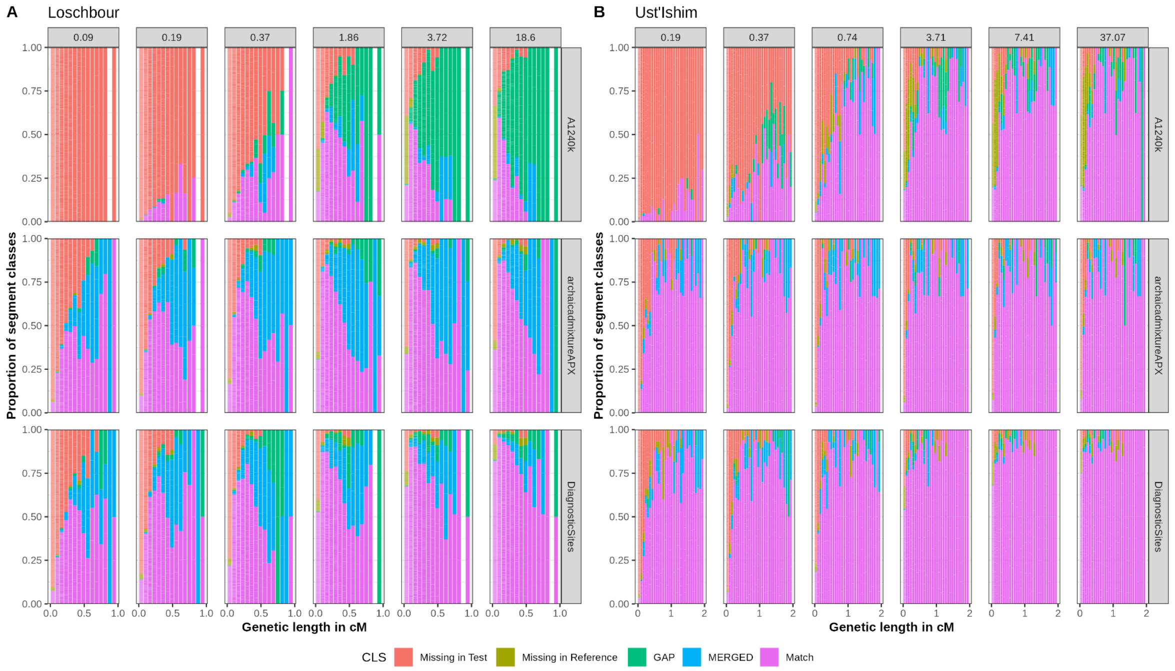

**Fig. S2.** Comparison of segment inference on the autosomes by *admixfrog* using three different ascertainment schemes, including 1240k, archaic admixture + X and "all-diagnostic sites" (in rows) with varying levels of coverage of two downsampled high coverage ancient individuals. The proportion of missing in test data (Missing in Test), missing in the reference (Missing in Reference), segments that are broken up (GAP), segments that are merged (MERGED) and matches (Match) is given per segment length bins, binned in centiMorgans (cM). The maximum coverage result on the "all-diagnostic sites" ascertainment per individual (rightmost column) is taken as the reference that lower coverages are compared to. (**A**) Comparison for the ~8,025 yBP old post-LGM hunter-gatherer genome of Loschbour. (**B**) Comparison for the ~44,300 yBP old early out-of-Africa Ust'-Ishim.

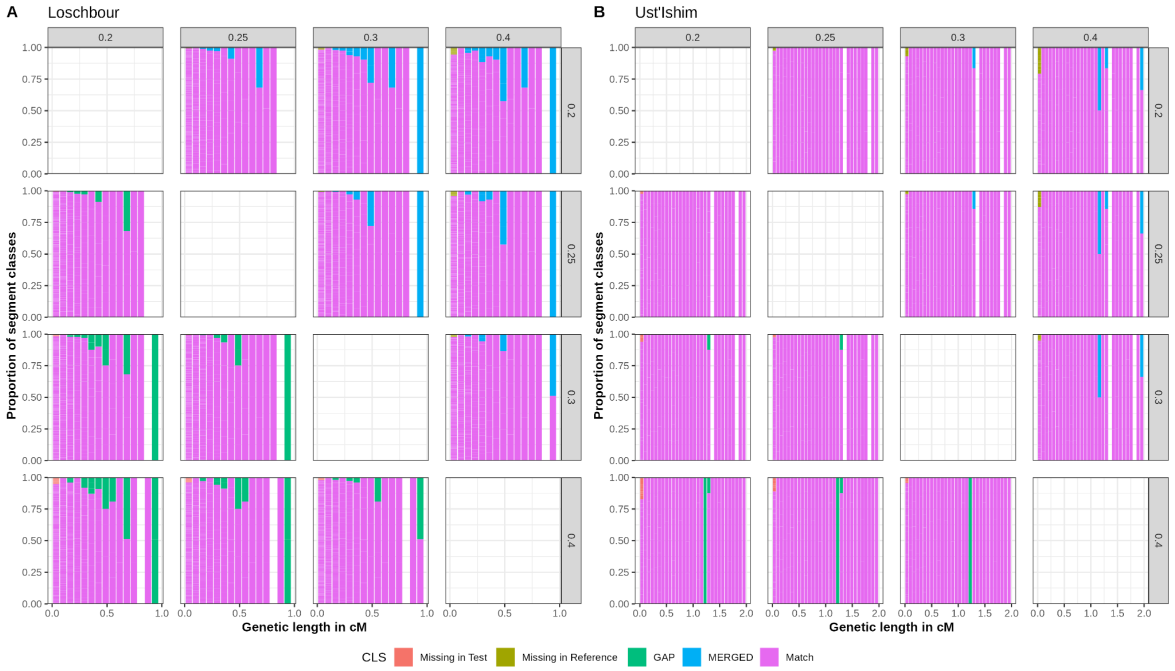

**Fig. S3.** Comparison of segment inference on the autosomes by *admixfrog* using four different values for the penalty parameter which regulates how generous the segments are being called for the archaic admixture + X ascertainment on high coverage ancient individuals. The proportion of missing in test data (Missing in Test), missing in the reference (Missing in Reference), segments that are broken up (GAP), segments that are merged (MERGED) and matches (Match) is given per segment length bins, binned in centiMorgans. Left facets give the reference parameter value and upper facets the test parameter. (**A**) Comparison for the ~8,025 yBP old post LGM hunter gatherer genome of Loschbour. (**B**) Comparison for the ~44,300 yBP old early out of Africa Ust'-Ishim.

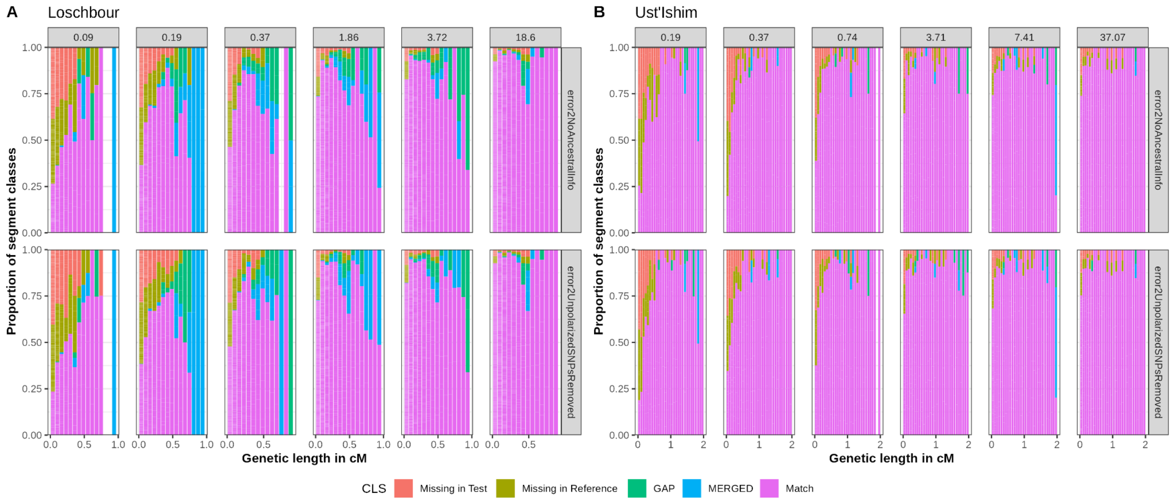

**Fig. S4.** Comparison of segment inference on the autosomes by *admixfrog* with unknown ancestral allele state (error2NOAncestralInfo) and only considering sites with ancestral state known (error2UnpolarizedSNPsRemoved) to the default setting on the archaic admixture + ArchaicX array. The proportion of missing in test data (Missing in Test), missing in the reference (Missing in Reference), segments that are broken up (GAP), segments that are merged (MERGED) and matches (Match) is given per segment length bins, binned in centiMorgans (cM). All parameters are compared to the reference results using the default parameter (error2), which adjusts the prior if an ancestral allele is present otherwise gives a uniform prior. Everything is compared under the same downsampled coverage (upper facts). (**A**) Comparison for the ~8,025 yBP old post-LGM hunter gatherer genome of Loschbour. (**B**) Comparison for the ~44,300 yBP old early out of Africa Ust'-Ishim.

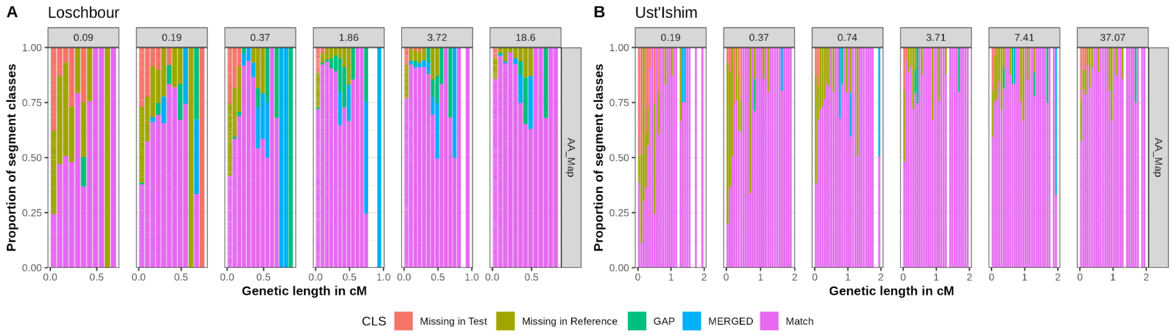

**Fig. S5.** Comparison of segment inference on the autosomes by *admixfrog* using a different genetic map (African-American map) on the archaic admixture + ArchaicX array. The proportion of missing in test data (Missing in Test), missing in the reference (Missing in Reference), segments that are broken up (GAP), segments that are merged (MERGED) and matches (Match) is given per segment length bins, binned in centiMorgans. All parameters are compared to the reference results using the default parameter (error2), which adjusts the prior if an ancestral allele is present otherwise gives a uniform prior. Everything is compared under the same downsampled coverage (upper facts) with segments called using the African-American map (AA map). (**A**) Comparison for the ~8,025 yBP old post-LGM hunter gatherer genome of Loschbour. (**B**) Comparison for the ~44,300 yBP old early out of Africa Ust'-Ishim.

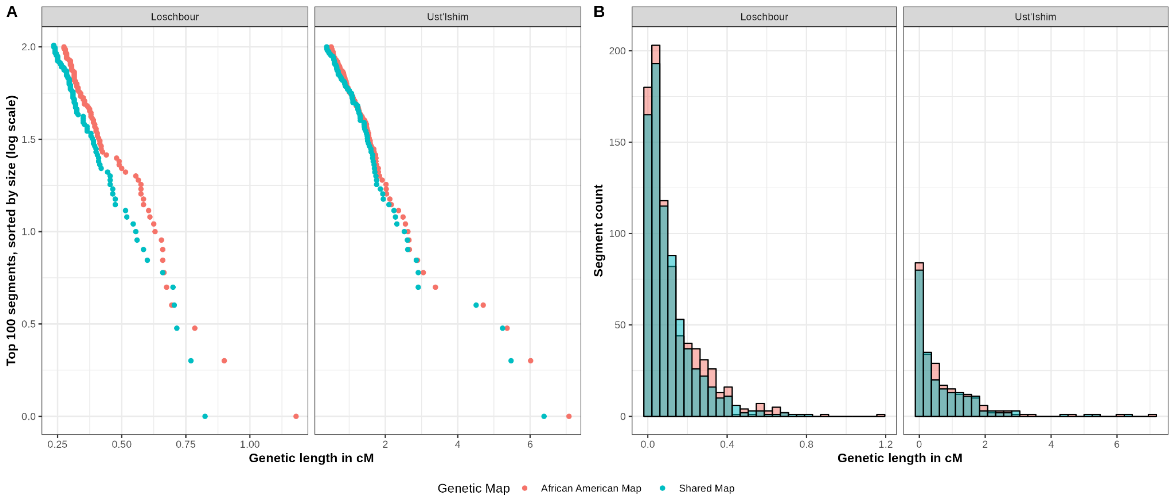

**Fig. S6.** Comparison of segment length distribution on the autosomes inferred by *admixfrog* using the Shared map (blue) vs. the African-American map (red) for the inference on the archaic admixture + ArchaicX array. (**A**) Comparison for the ~8025 yBP old post-LGM hunter gatherer genome of Loschbour and the ~44,300 yBP old early out of Africa Ust'-Ishim on the distribution of the top 100 segments sorted by size (in cM) and plotted in a log scale per map and individuals. (**B**) Histogram of the counts of segments with a given genetic length for the same individuals.

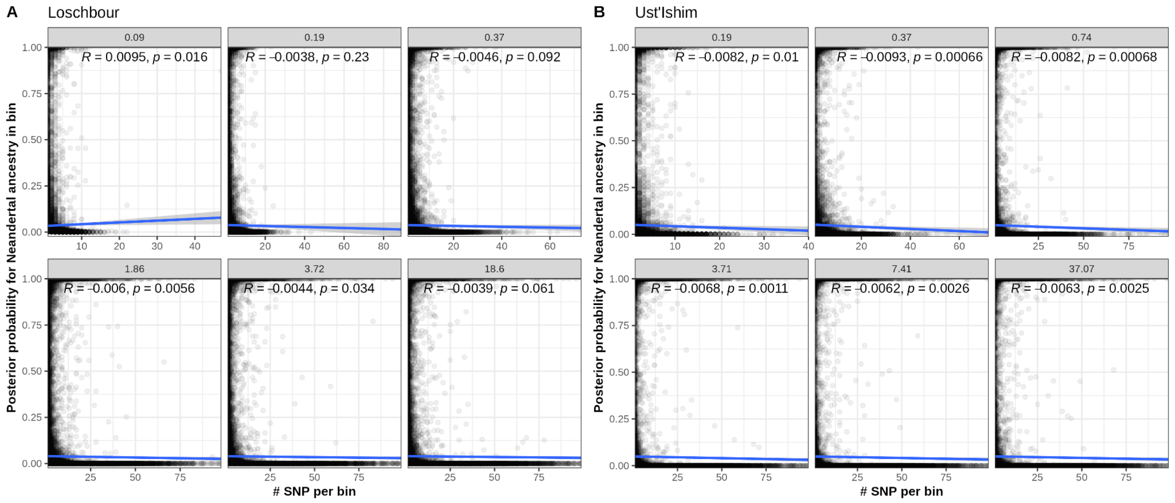

**Fig. S7.** Correlation of the number of SNP per bin vs. posterior probability for Neandertal ancestry for different downsampled coverages. Bins with zero SNPs are removed (**A**) Correlation for the ~8,025 yBP old post-LGM hunter gatherer genome of Loschbour. (**B**) Correlation for the ~44,300 yBP old early out of AfricaUst'-Ishim.

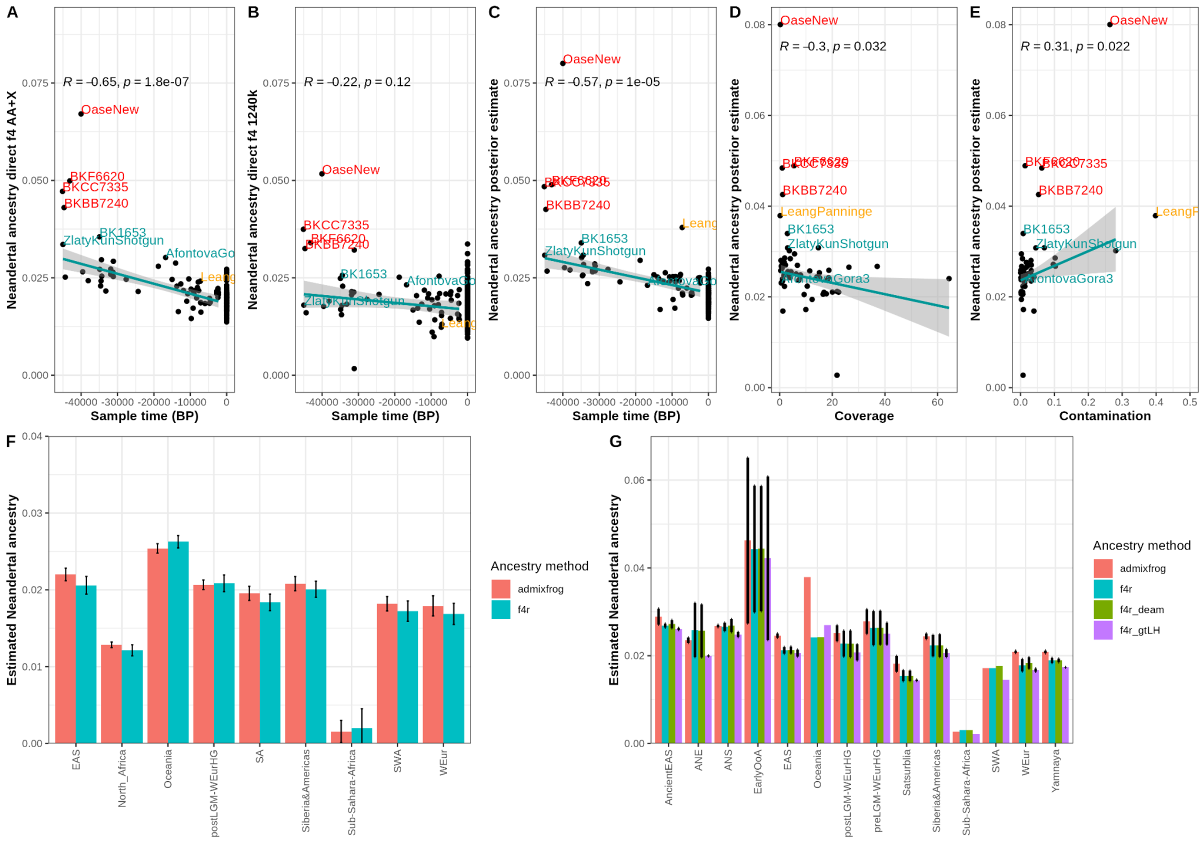

**Fig. S8.** Ancestry estimates using the overall posterior probability from *admixfrog* and direct f4 ratios on the archaic admixture sites. **A - E** direct f4 estimates or posterior probability correlated with: the age of the individuals, coverage and contamination. Individuals excluded from the fitting due to being previously suggested to have more than 1 pulse of Neandertal ancestry are marked in red. Pearson's correlation coefficient with p-value is given in the upper left corner (**F**) Comparison between direct f4 ratio and *admixfrog* for present-day individuals. (**G**) Comparison between *admixfrog* and direct f4 ratio on ancient individuals. F4 ratios are calculated either on input data with all SNPs (blue), deaminated data only (green) or using the genotype likelihoods from *admixfrog* (purple).

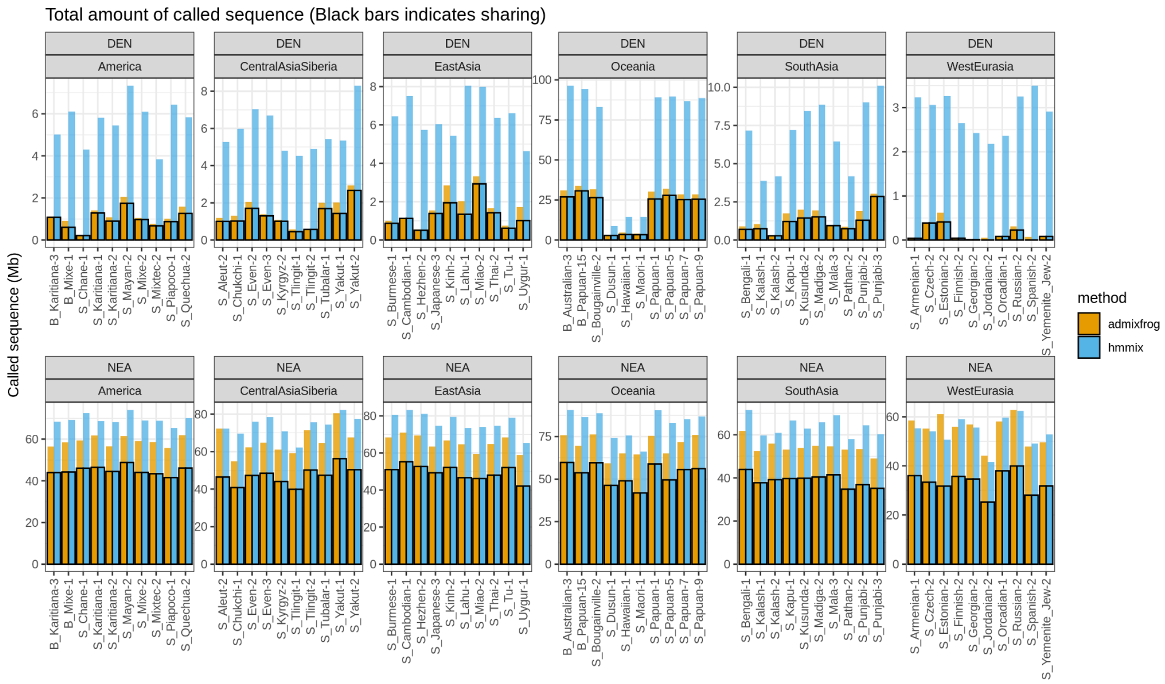

**Fig. S9.** For each individual (10 for each continental group) we show the amount of archaic sequence which is more similar to Denisovans and Neandertals identified by each method (orange for *admixfrog* and blue for *hmmix*). The colored bars indicate the amount of shared archaic sequence.

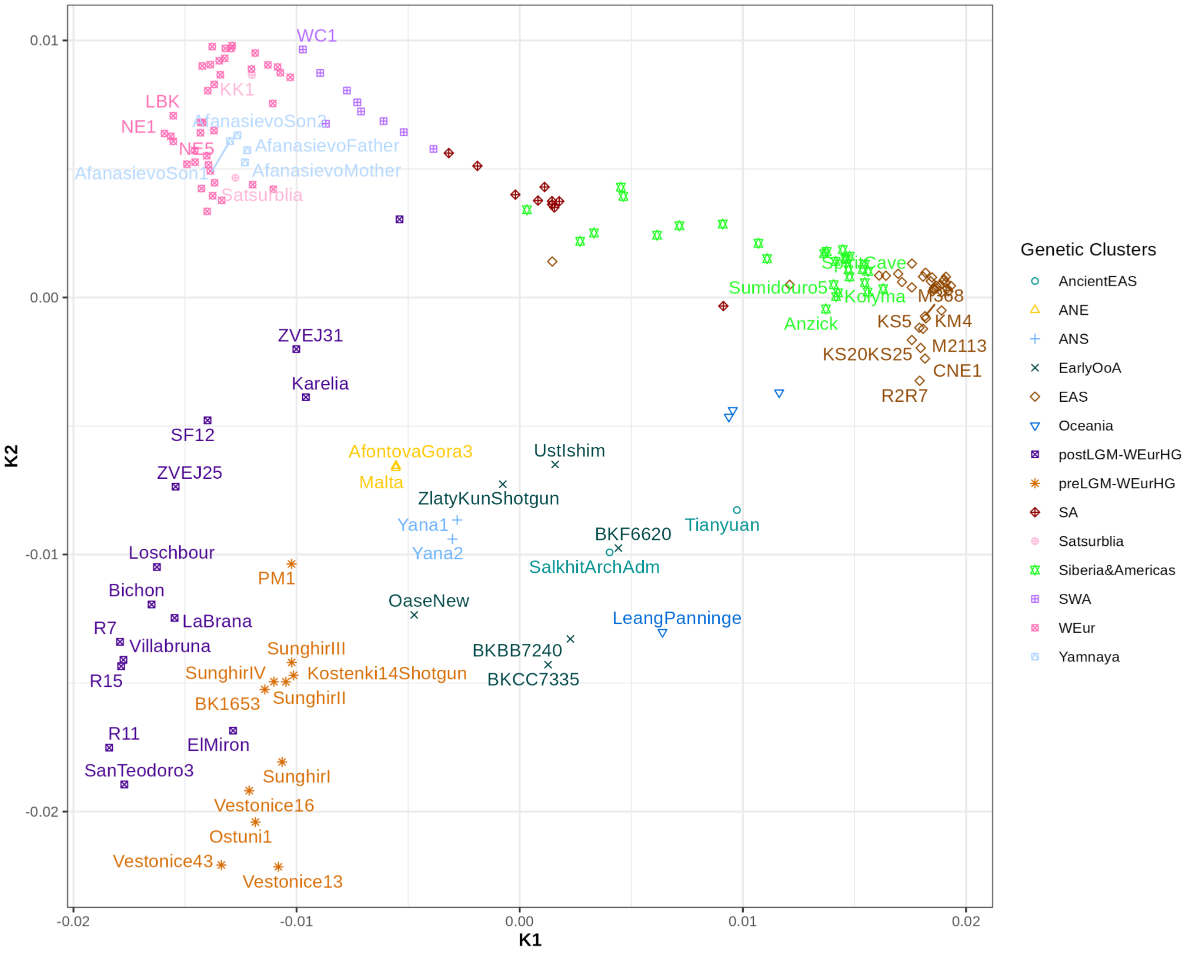

**Fig. S10.** MDS plot of the genetic clusters using outgroup-$f_3$ dissimilarities between populations/individuals for K 1 and K 2. Individuals are colored by the resulting genetic clusters with the following acronyms: ancient East-Asians (ancientEAS), ancient North-Eurasians (ANE), ancient North-Siberians (ANS), early out of Africa (EarlyOoA), East-Asians (EAS), post Last Glacial Maximum West-Eurasian Hunter Gatherers (postLGM-WEurHG), pre Last Glacial Maximum West-Eurasian Hunter Gatherers (preLGM-WEurHG), South-Asians (SA), South-West-Asians (SWA), West-Eurasians (WEur).

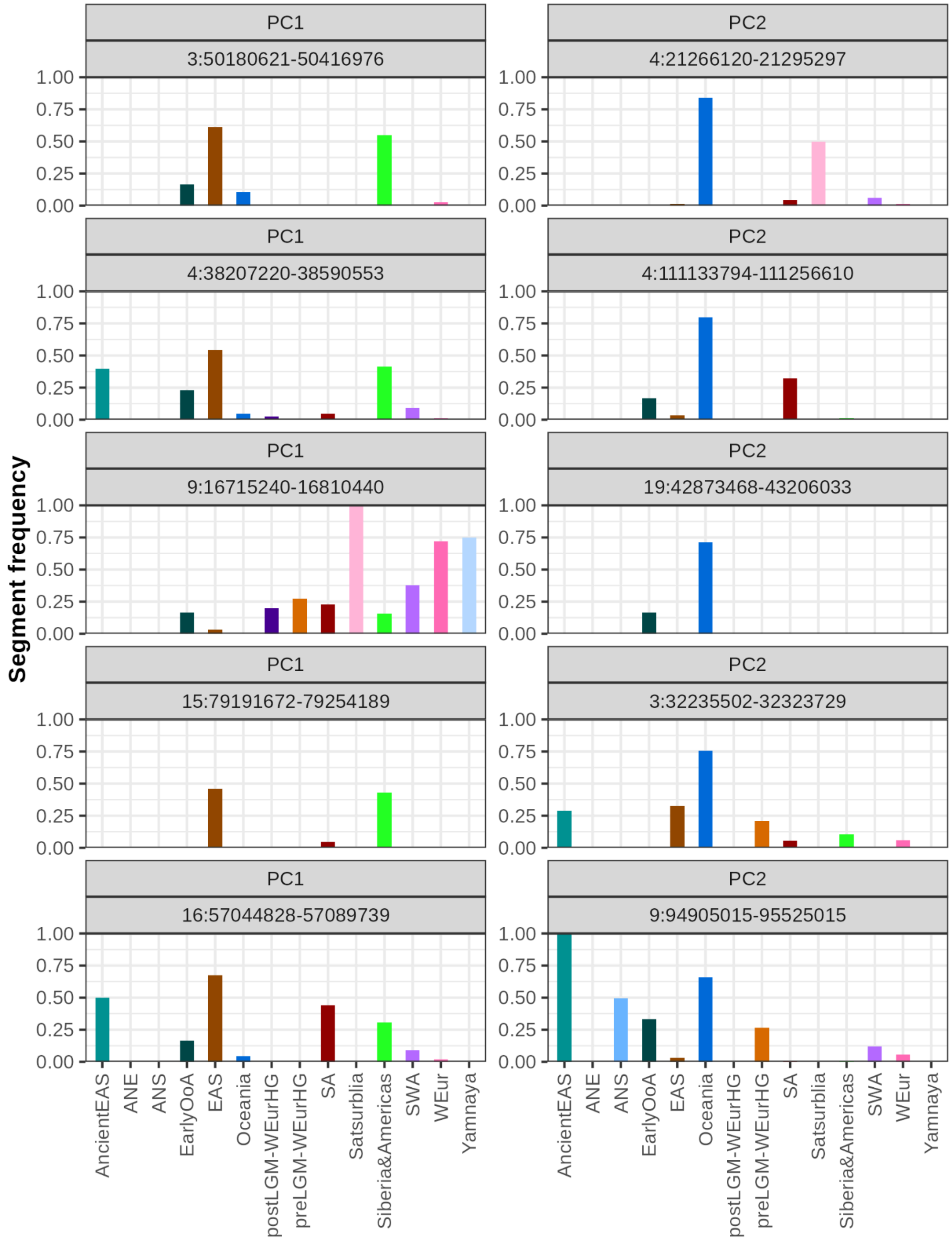

**Fig. S11.** Frequency of the five highest loading segments (sorted top to bottom) on PC 1 and PC 2 for each genetically informed cluster. Upper facet indicates the PC and lower facet the position in chromosome:start in bp - end in bp. Colors correspond to the genetic cluster.

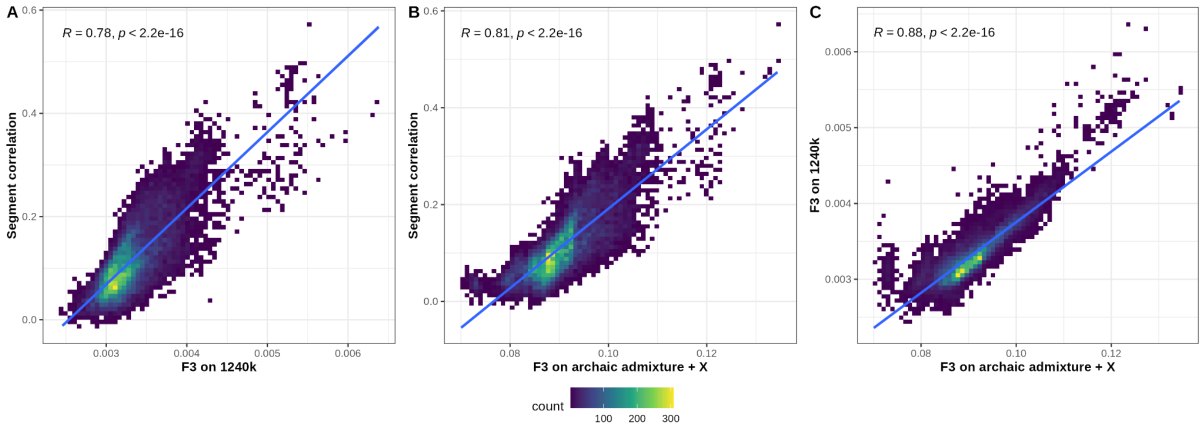

**Fig. S12.** Linear regression for following: (**A**) Neandertal segment correlation vs. $f_3$ values computed on 1240 k sites. (**B**) Neandertal segment correlation vs. $f_3$ values computed on archaic admixture + X sites. (**C**) $f_3$ values computed on 1240 k sites vs. $f_3$ values computed on archaic admixture + X sites. With the spearman correlation coefficient and p-value given in the upper right corner.

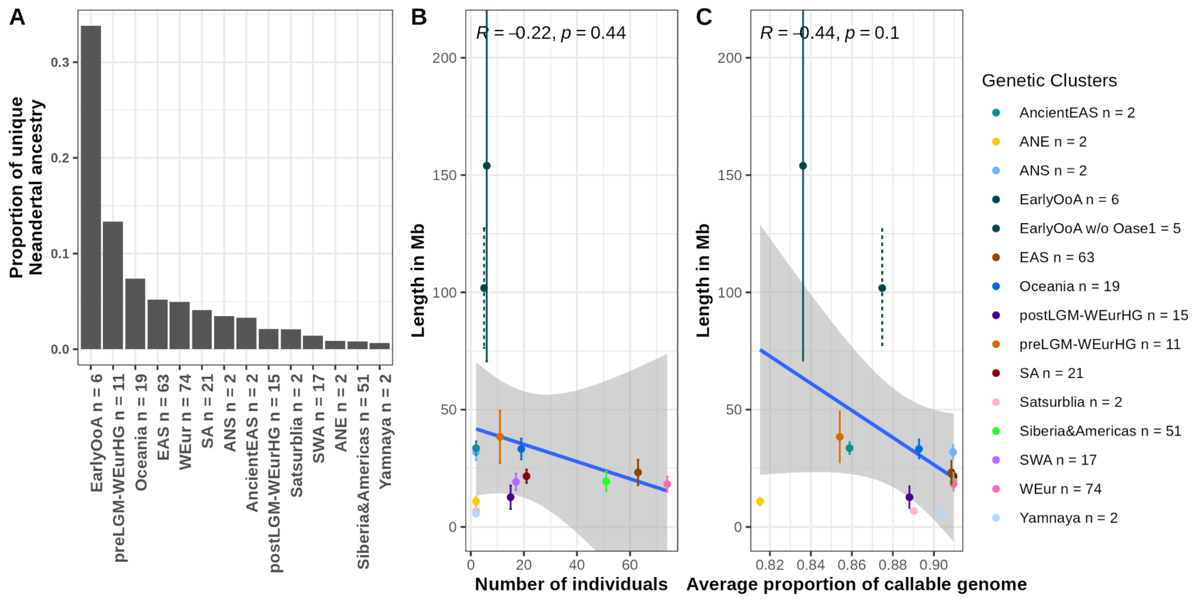

**Fig. S13.** (**A**) Proportion of Neandertal ancestry unique to the population cluster and not shared with any other cluster. Shared Neandertal ancestry is 1 - proportion of unique Neandertal ancestry. (**B**) Correlation of the number of individuals per cluster vs. the amount of unique Neandertal ancestry pre population cluster (with n given the number of individuals per cluster) in Mb from a randomly sampled individual. (**C**) Correlation of average proportion of callable genome in the individuals of a particular cluster vs. the amount of unique Neandertal ancestry pre population cluster in Mb from a randomly sampled individual. Spearman's correlation coefficient rho and p-value indicated. Error bars are calculated by resampling the individuals, the EarlyOoA cluster is calculated with (solid line) and without including the Oase1 individual (dotted line). Spearman correlation was performed with the result depicted in the upper right corner.

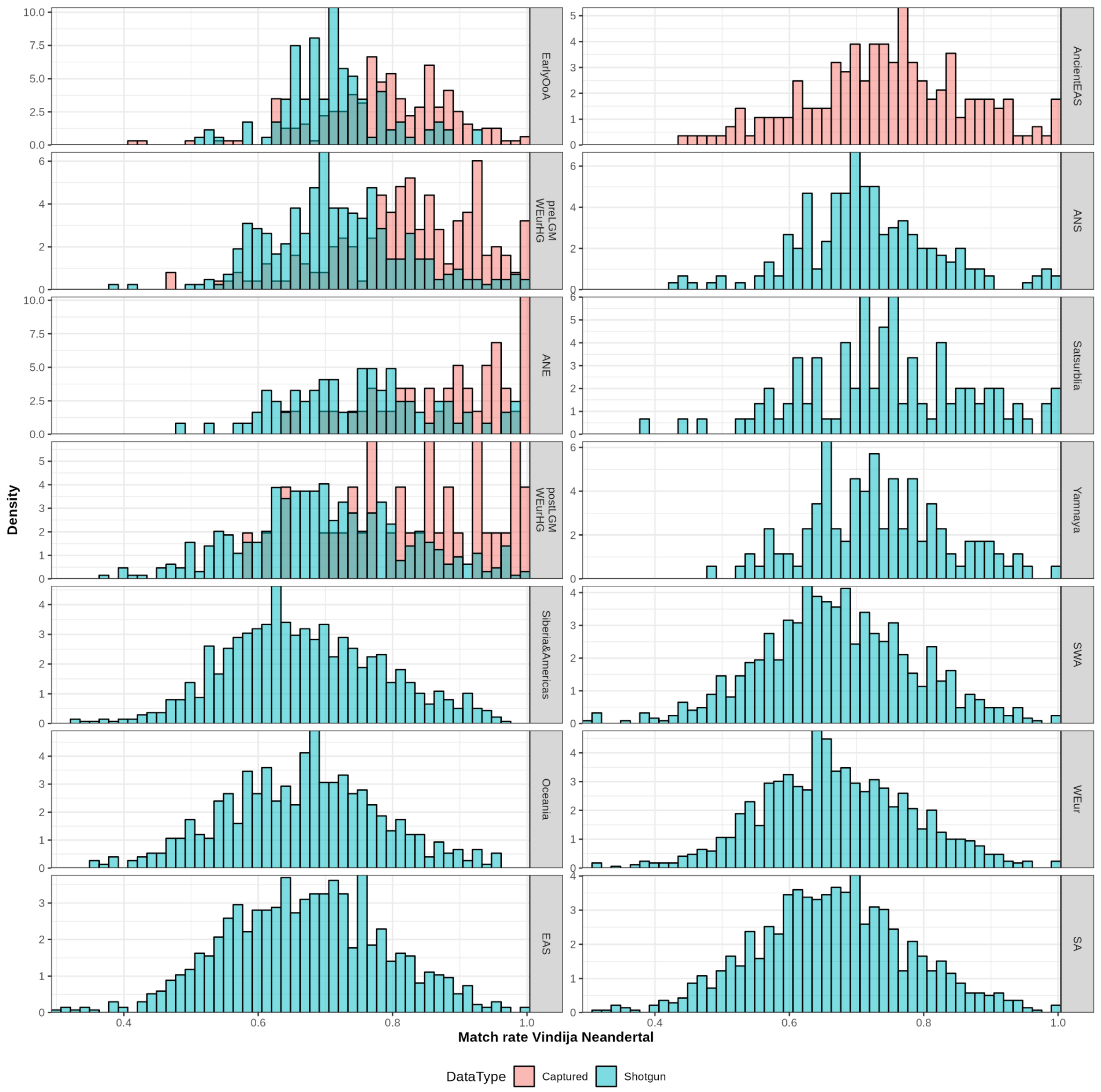

**Fig. S14.** Matching rate of Neandertal segments with at least 15 informative SNPs to the Vindija Neandertal, stratified by data type (shotgun sequenced in blue or captured in red). Individuals are grouped in population clusters ordered from oldest to youngest.

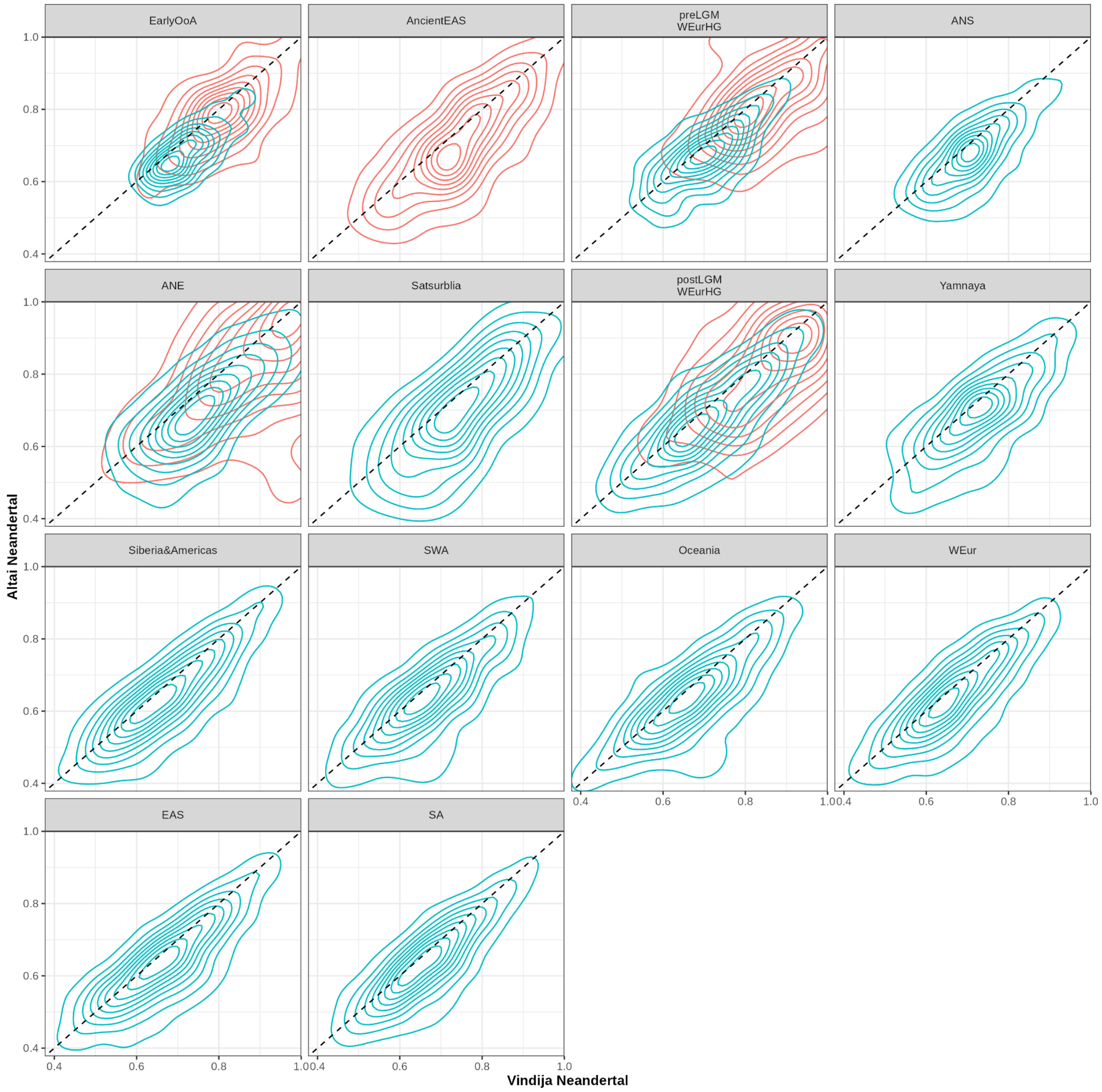

**Fig. S15.** Density plot of the matching of Neandertal segments on autosomes with at least 15 informative SNPs to the Vindija vs. Altai Neandertal by genetic population cluster. Clusters are ordered from oldest to youngest. Blue density lines indicate matching for segments from individuals that are shotgun sequenced and red for individuals which were capture enriched.

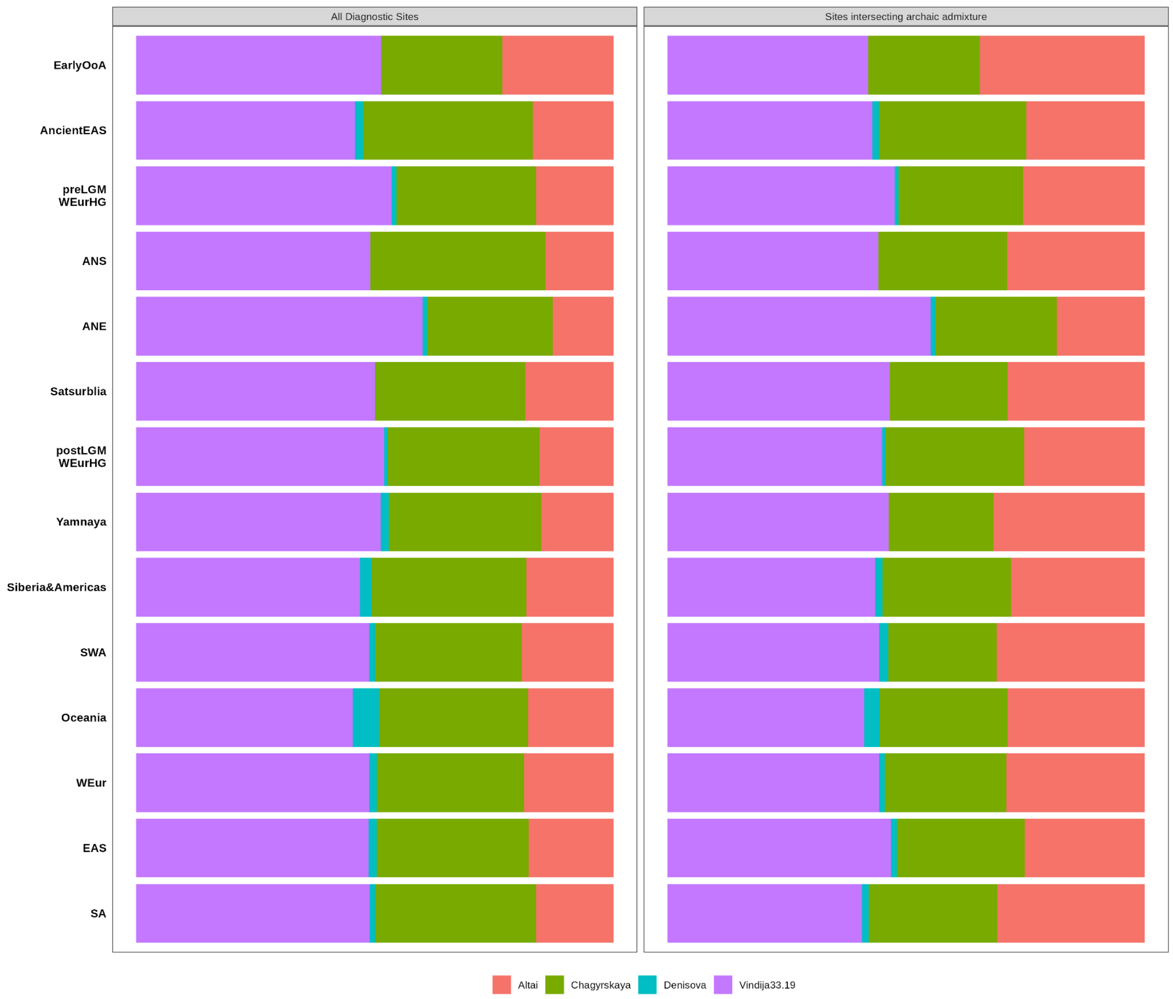

**Fig. S16.** Composition of autosomal segments in a population assigned to a specific archaic reference. The different colored bars represent the proportions of Neandertal segments matching closest to one of the four archaic references using the diagnostic sites ascertainment. The populations are stratified by time into individuals from oldest to youngest. Compositions are depicted for segments using all sites from the all diagnostic sites ascertainment or only sites that also overlap with the archaic admixture ascertainment.

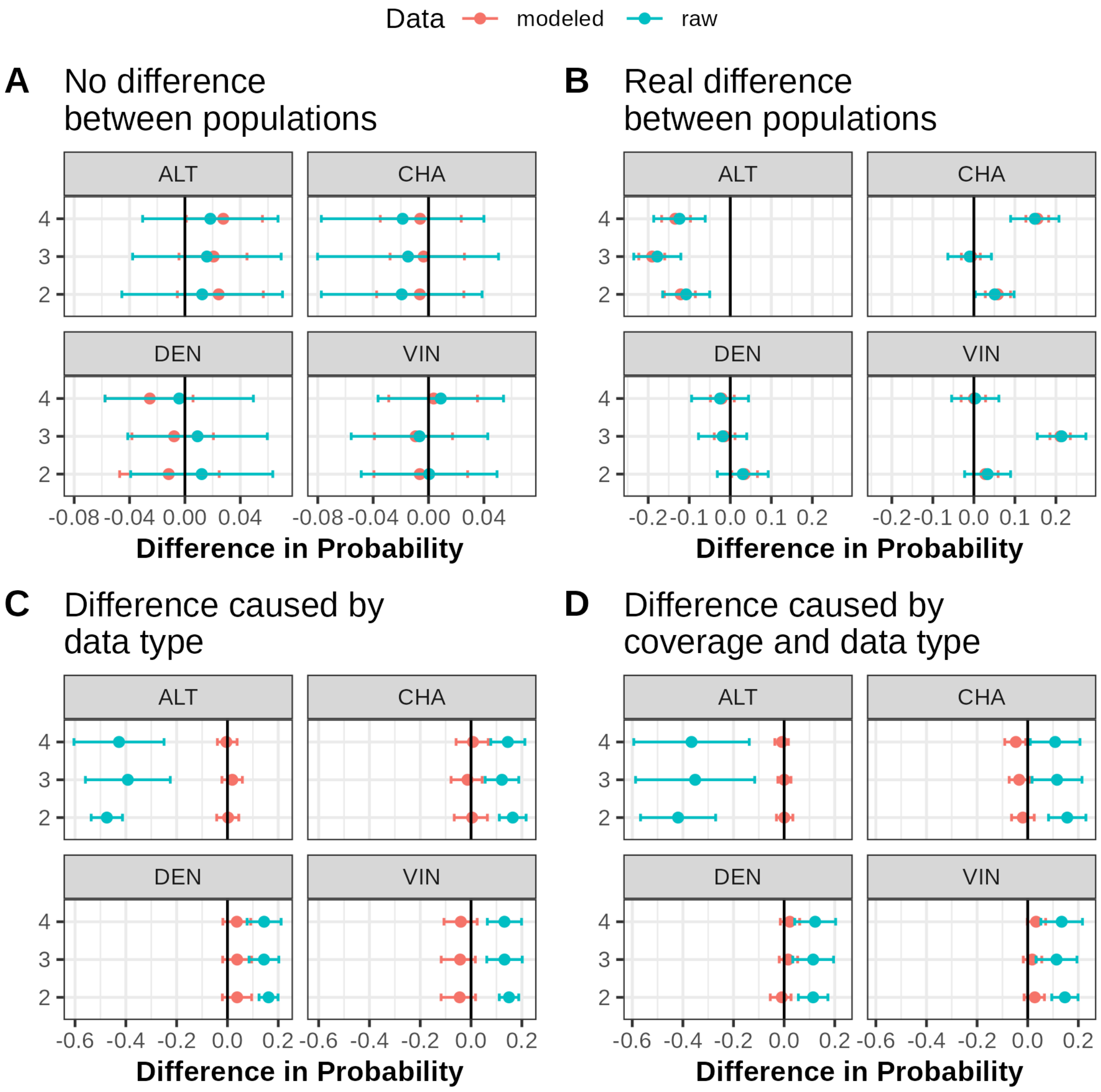

**Fig. S17.** Simulation to test the modeling for the probability of a Neandertal segment coming from a certain archaic reference population using the Generalized Mixed Linear Model. Blue points with error bars indicate the raw mean and standard deviation (sd) of the proportion (y axis) of segments matching to each ancestry between population 2, 3 and 4 (x axis) against the raw mean and sd from the pivot population (population 1). Red points with error bars indicate the modeled mean and 95 % highest posterior density interval (HPDI) of the proportion of segments matching each ancestry between population 2, 3 and 4 against the modeled mean and 95 % HPDI from the pivot population. Error bars overlapping black line indicate that the standard deviation or 95 % HDPI of the pivot population overlaps with the population it is compared to. (**A**) Simulations with no difference between the three simulated populations. (**B**) Simulations with real differences between simulated populations. (**C**) Simulations with differences caused by different proportions of individuals being shotgun vs. capture sequenced between the populations. (**D**) Simulations with differences caused by a combination of differences in proportions of individuals being shotgun vs. capture sequenced and coverage between the populations.

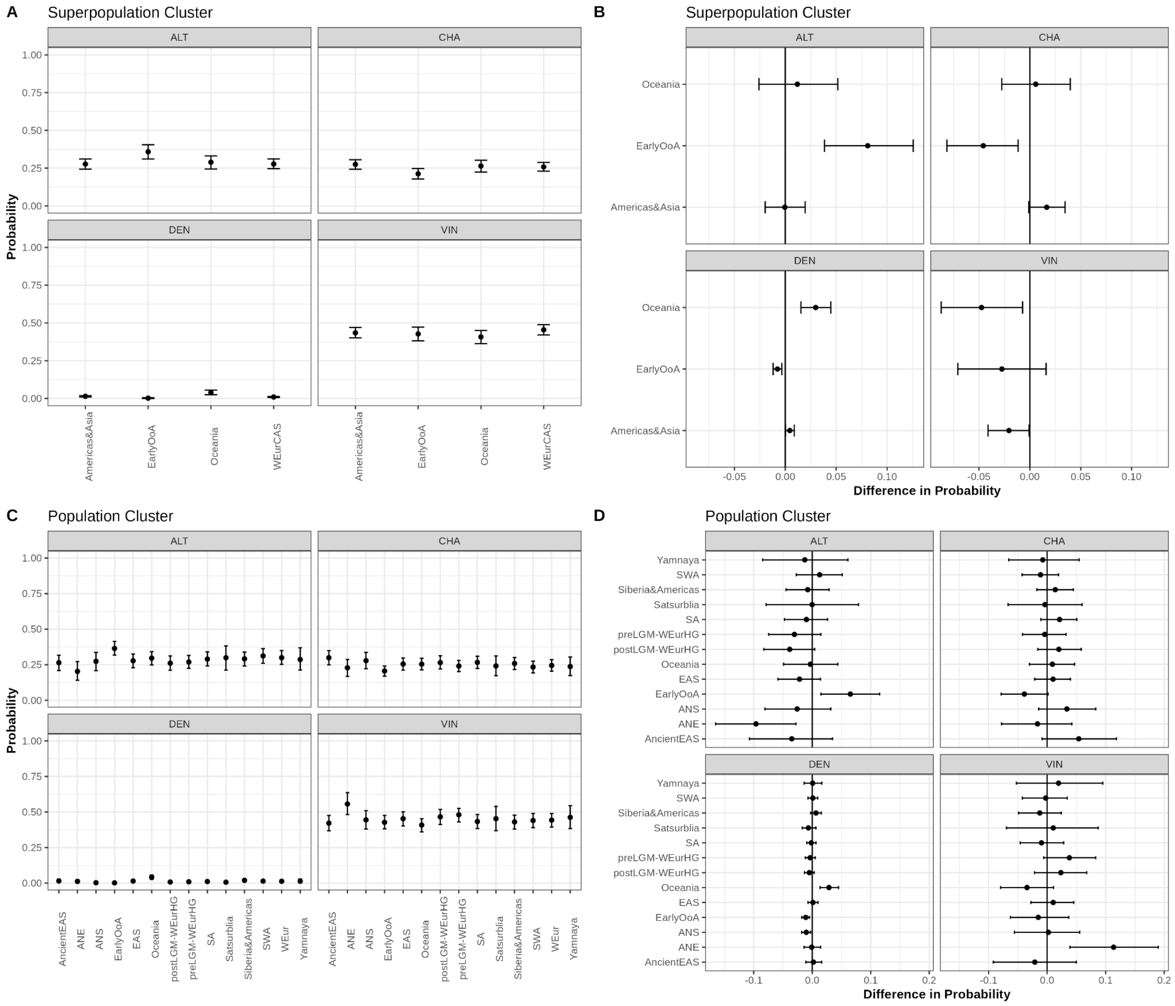

**Fig. S18.** GLM posterior predictions results for the proportion of autosomal segments best matching an archaic reference with predictors being data type (shotgun vs. capture) and sites (all diagnostic sites vs. all diagnostic sites overlapping the archaic admixture ascertainment). (**A**) Probability of a segment coming from a certain archaic reference population across superpopulation clusters. (**B**) Difference in probability of a segment coming from a certain archaic reference population between West-Eurasians Central Asians and Siberians (black line) taken as a pivot and the three other superpopulation clusters. Errorbars overlap black line indicates that the 95 % HDPI of the West-Eurasians Central Asians and Siberians cluster and the estimate for the other superpopulations cluster overlap. (**C**) Same as A but for population clusters. (**D**) same as B but for population clusters with the pivot population being West-Eurasians.

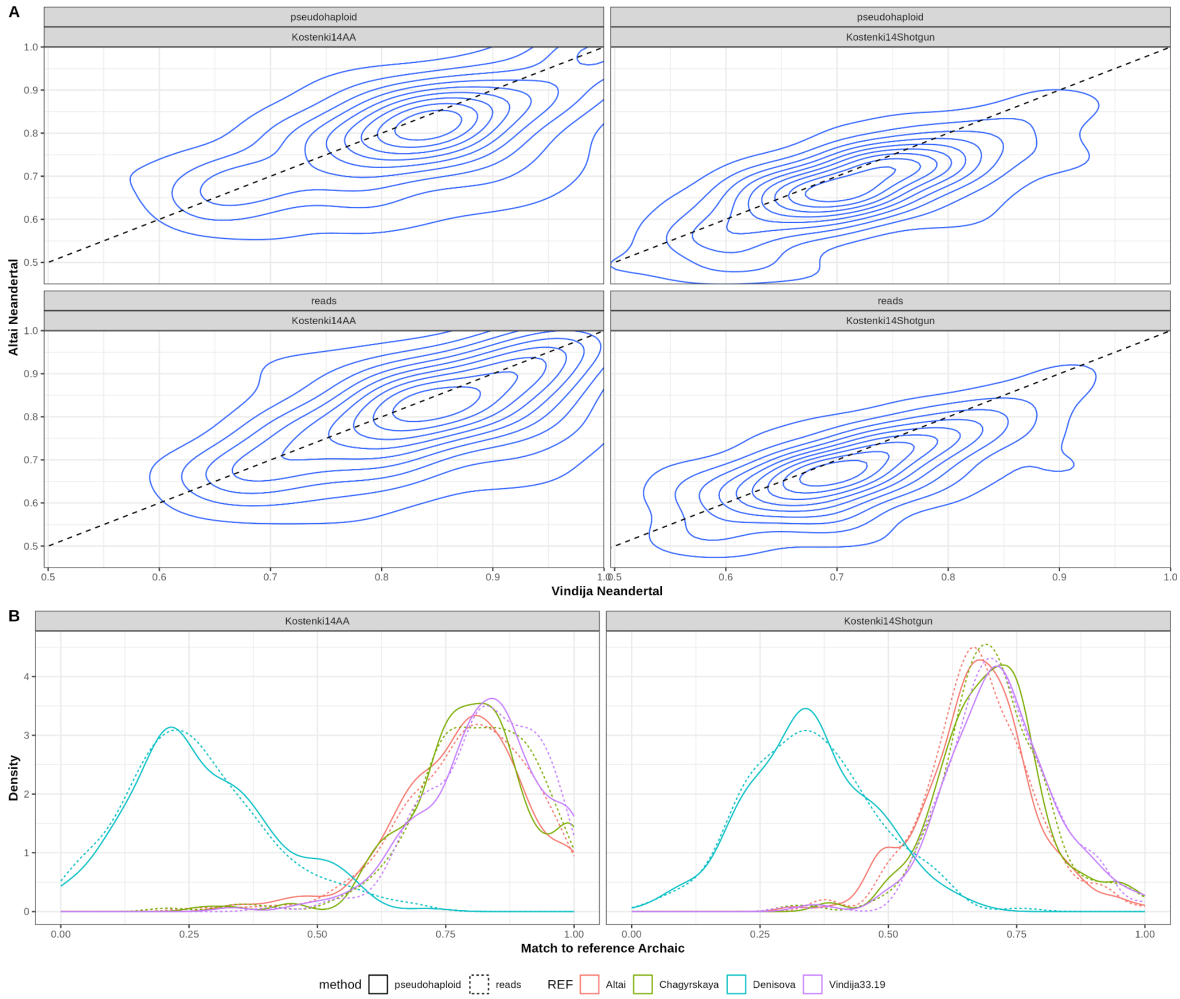

**Fig. S19.** Validation of lineage assignment using Kostenki14 shotgun and captured data calculated using proportions of reads carrying the derived allele of a randomly sampled allele. (**A**) Contour plot of matching of segments with minimum length of 0.2 cM and at least 10 informative SNPS to the Vindija and Altai reference Neandertals. (**B**) Density plot of matching of segments to all archaic references.

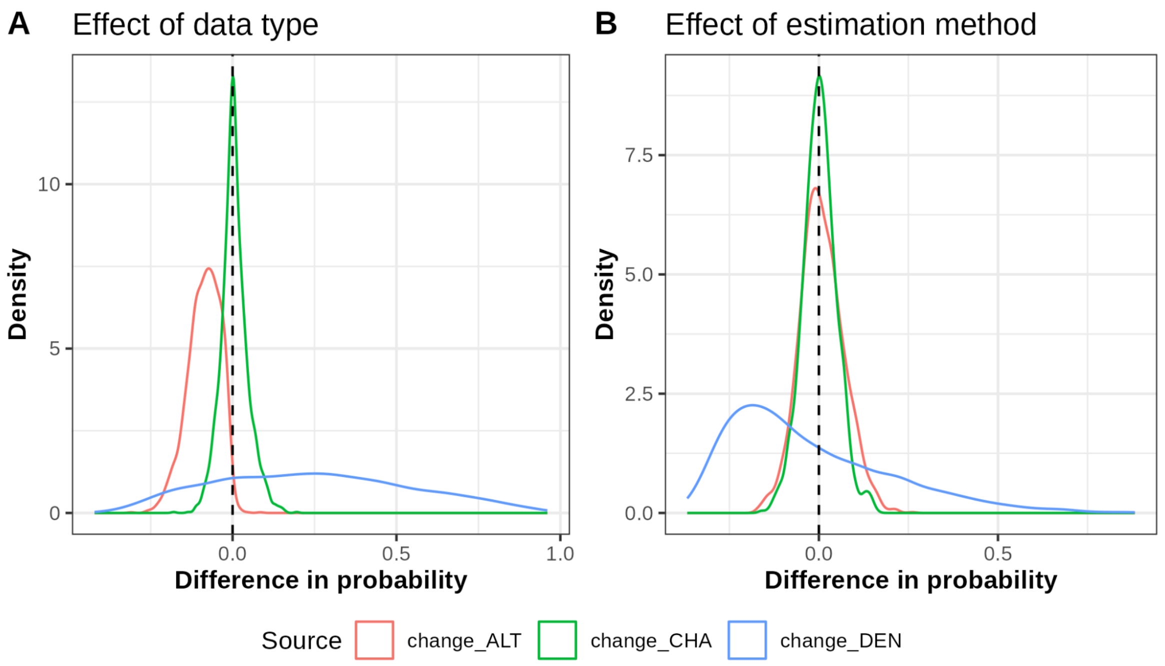

**Fig. S20.** GLM posterior probability estimates on the validation of lineage assignment using Kostenki14 shotgun and captured data calculated using proportions of reads carrying the derived allele of a randomly sampled allele. (**A**) Posterior probability of the effect estimated for data type (**B**) Posterior probability of the effect estimated for method. Dotted line indicates no significant difference in probability i.e., there is minimal effect of data type or method on our inference.

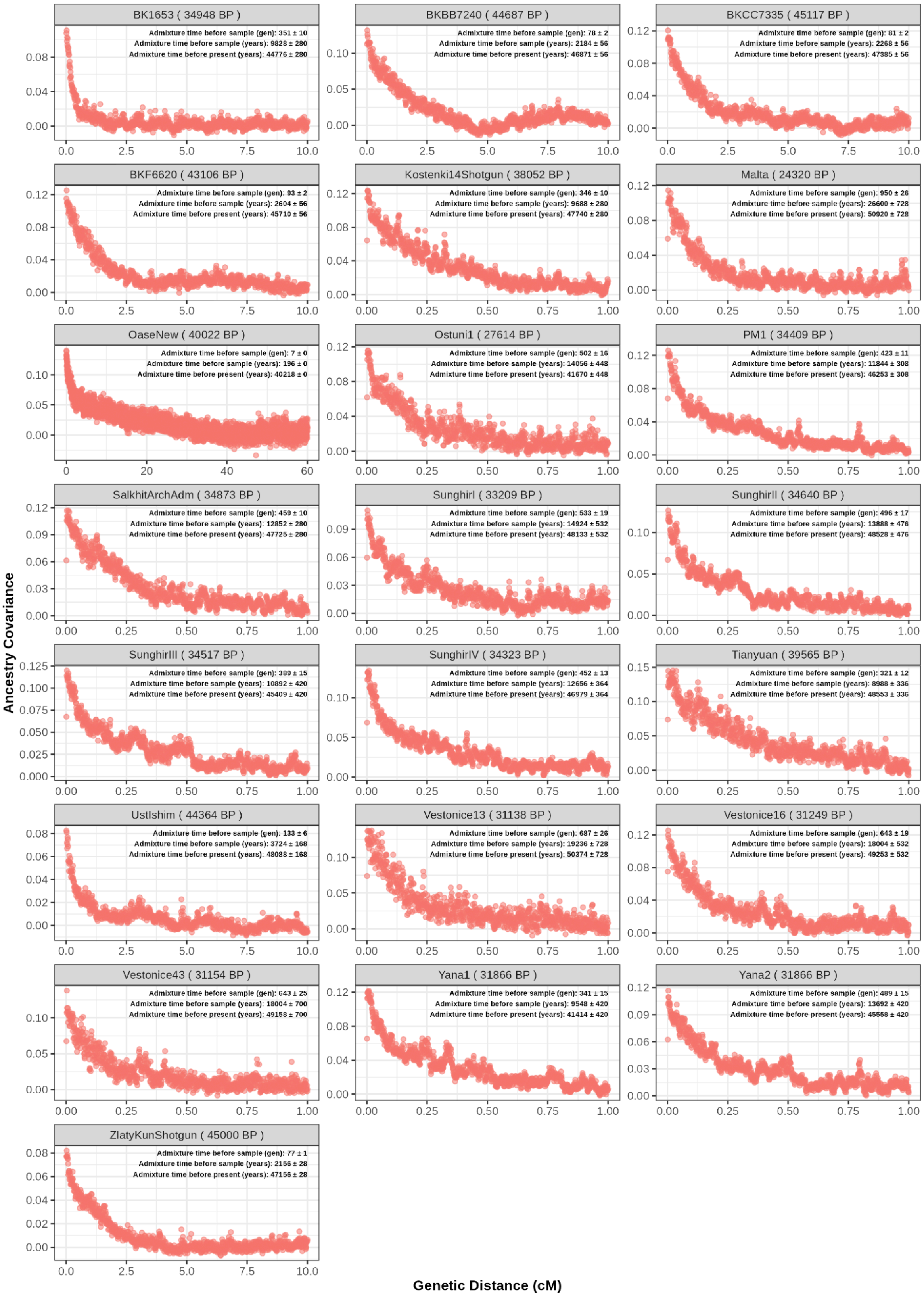

**Fig. S21.** Ancestry covariance curves from 22 ancient individuals with their inferred dates of admixture in generations and years. We assume a human generation time of 28 years. We show the individual name and average sampling age (BP) in the title of each subplot.

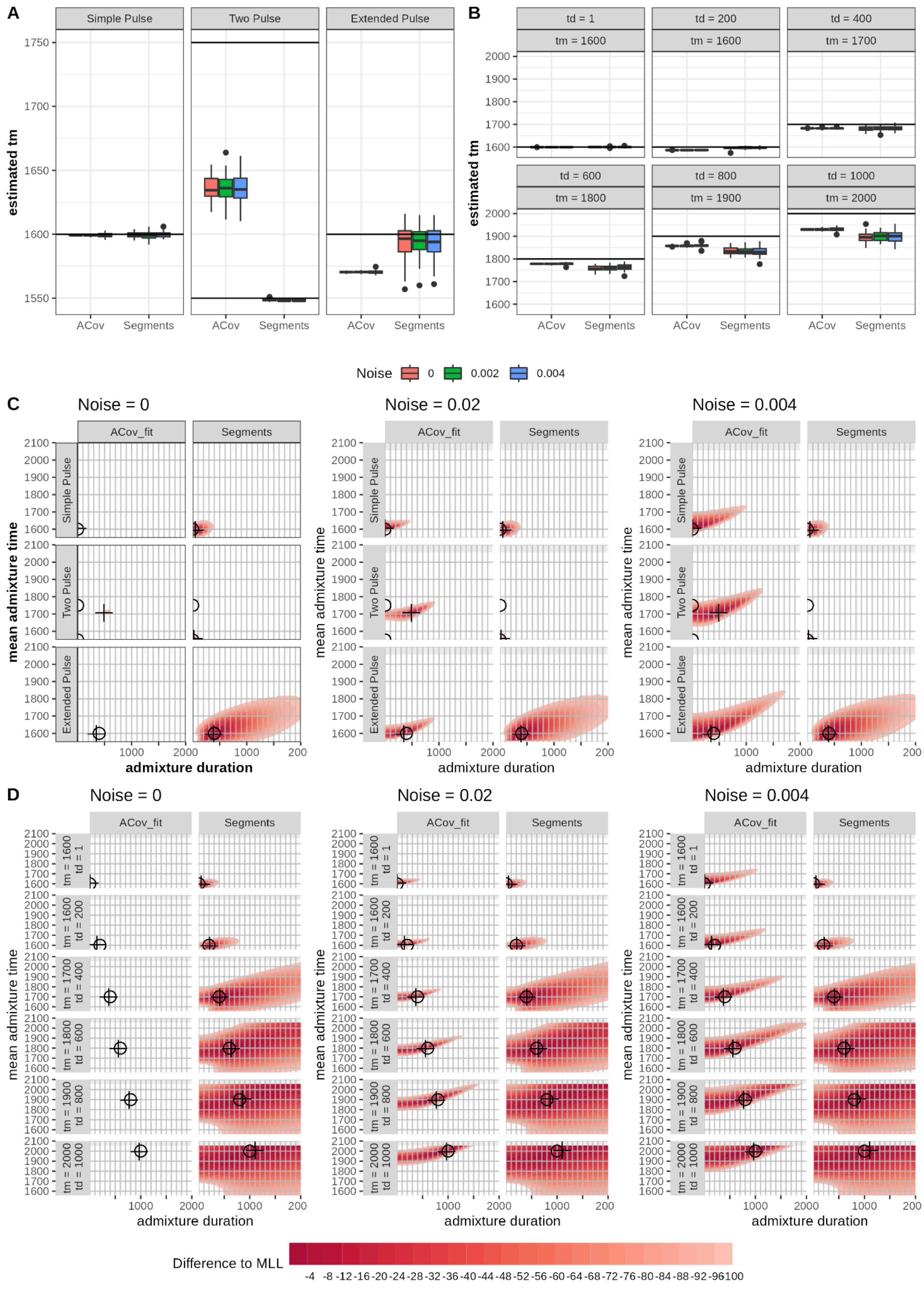

**Fig. S22.** Estimates of mean admixture time and admixture duration from simulations under different admixture models using Ancestry Covariance (ACov) or segments (**A**) IG model fit to 50 replicates of a IG model, two pulses and extended pulse model with different levels of gaussian noise. Extended pulse has a duration of 400 generations. (**B**) IG model fit to 50 replicates of extended pulses with different mean times and durations of admixture with different levels of gaussian noise. Vertical lines indicate true values. (**C**) Grid of likelihoods for extended pulse fit to 50 replicates of a IG model, two pulses and extended pulse model with different levels of gaussian noise. (**D**) Grid of likelihoods for extended pulse fit to 50 replicates of an extended pulse with different mean times and durations of admixture with different levels of gaussian noise. Shades of red indicate differences in log-likelihood between the highest log-likelihood being 0 and all subsequent being smaller. Circles indicate true values, pulse signs the combination of mean time and duration with the highest log-likelihood across the 50 replicates.

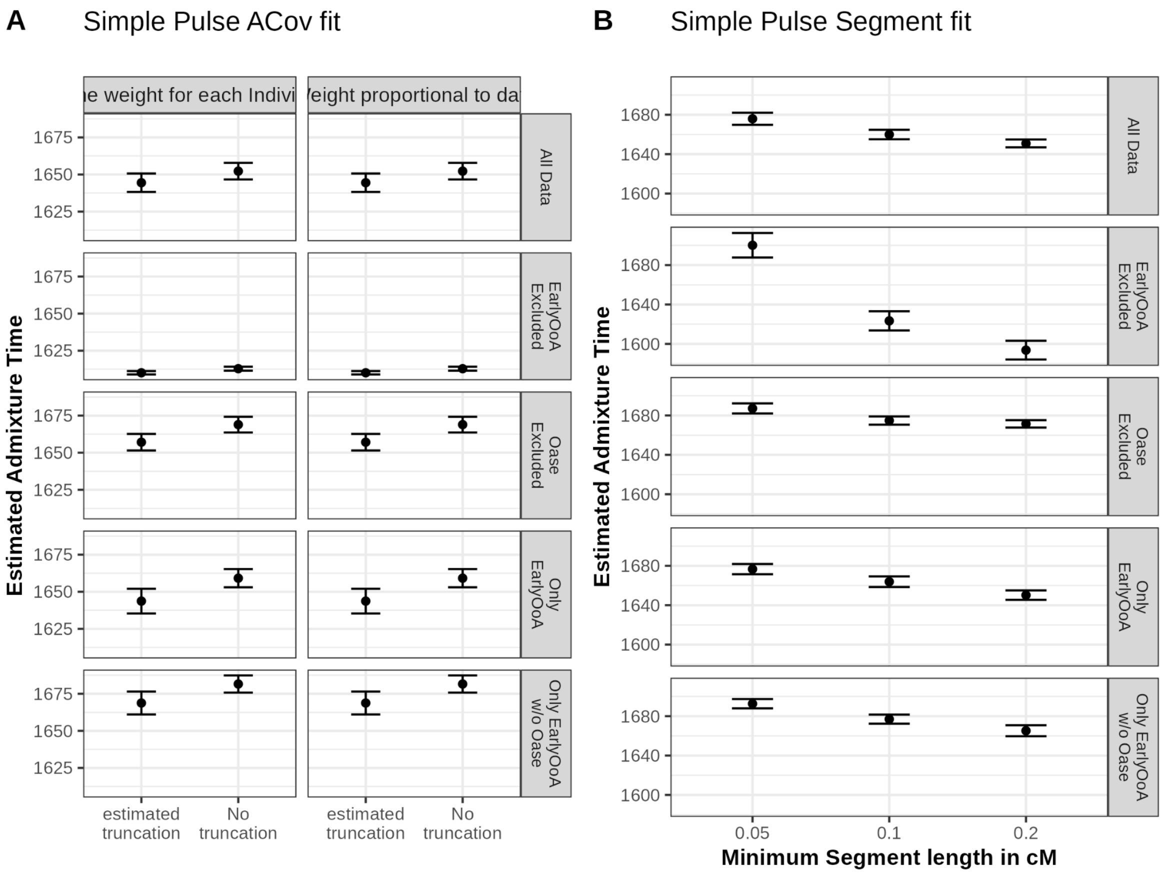

**Fig. S23.** Mean admixture time estimates with upper and lower value for the joint fitting of a one generation introgression pulse (IG model) to multiple subsets of individuals older than 22 kyBP. This is either to all data (n = 22), the Early out of Africa (EarlyOoA) individuals excluded (n = 16), Oase1 excluded (n = 21), Only EarlyOoA (n = 6) and only EarlyOoA without Oase1 (n = 5). (**A**) Mean time of admixture on subsets fitted to the ancestry covariance curves with either estimated maximum length or given maximum length (estimated truncation vs. no truncation) and either all individuals are weighted equally or the weight on the log-likelihood is proportional to the amount of data. (**B**) Mean time of admixture on data subsets fitted to the ancestry covariance curves for 100 bootstraps (on the age of the individuals) and jackknife (by leaving out one individuals at a time) replicas using estimated truncation and individuals with equal weight on the log-likelihood.

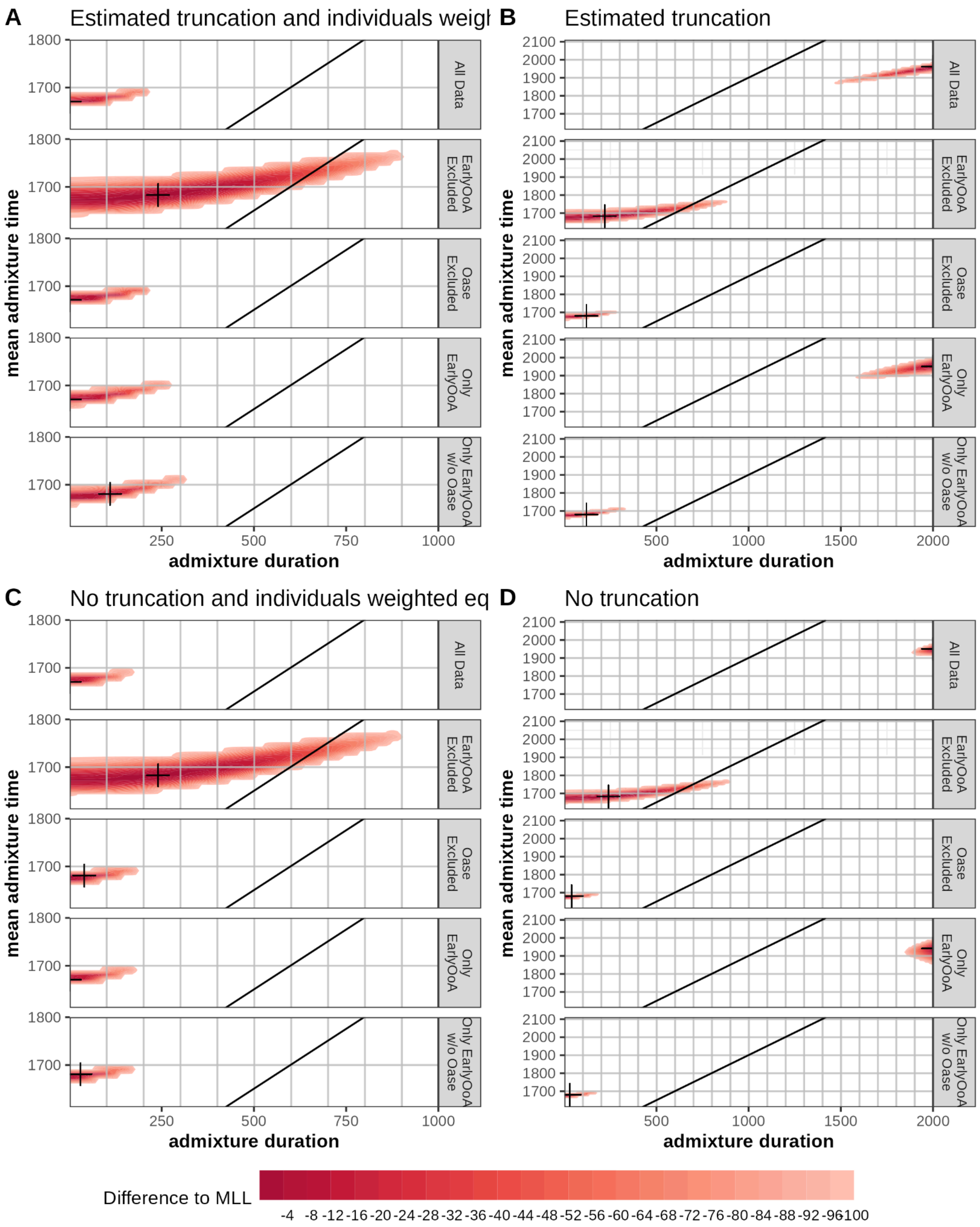

**Fig. S24.** Log-likelihood surface for different possible values for the mean time $t_m$ vs. duration of admixture $t_d$ fitted to ancestry covariance curves to multiple subsets of individuals older than 22 kyBP. Either to all data (n = 22), the Early out of Africa (EarlyOoA) individuals excluded (n = 16), Oase1 excluded (n = 21), Only EarlyOoA (n = 6) and only EarlyOoA without Oase1 (n = 5). (**A**) Using Ancestry covariance curves with an estimated truncation and individuals having an equal weight on the log-likelihood. (**B**) Using Ancestry covariance with an estimated truncation and no weighting. (**C**) Using Ancestry covariance with no further truncation and individuals having an equal weight on the log-likelihood. (**D**) Using Ancestry covariance with no truncation and no weighting. Shades of red indicate the difference between any log-likelihood and the highest log-likelihood. Black diagonal line indicates the last archeological date of Neandertals taken from (*30*).

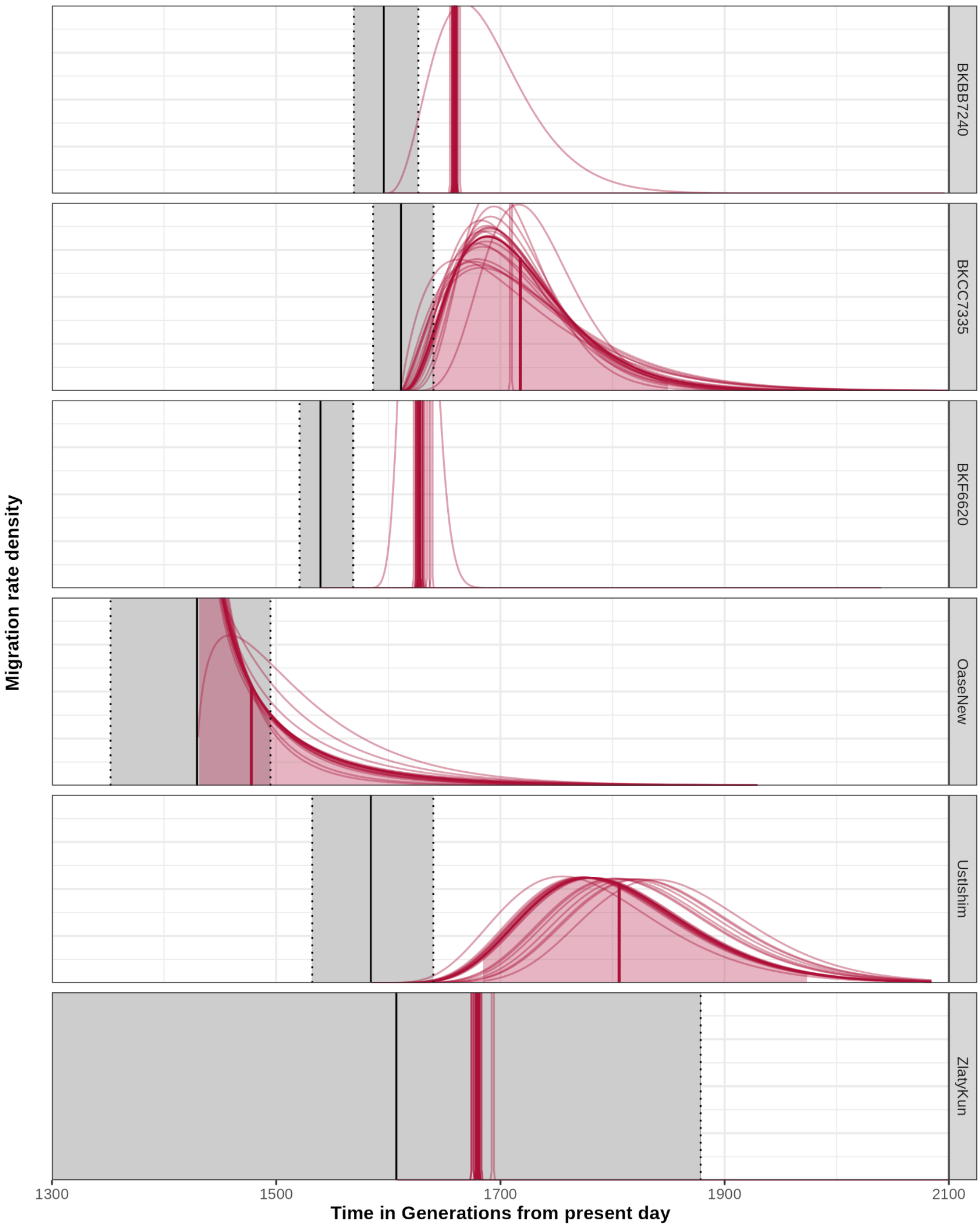

**Fig. S25.** Model prediction for the EarlOoA individuals. Predicted migration rate density over time for each of the six Early out of Africa (EarlyOoA) individuals under the extended pulse. Dark red line indicates the estimate for all chromosomes with the vertical line gives the estimated mean line of admixture and the shaded are the 95.5 percentile of the distribution. Light red lines indicating Jackknife results for leaving one chromosome out at a time. Gray rectangles is the range of the most likely sample time per individual (back line gives the mean and dotted line the upper and lower 95.5 percentile)

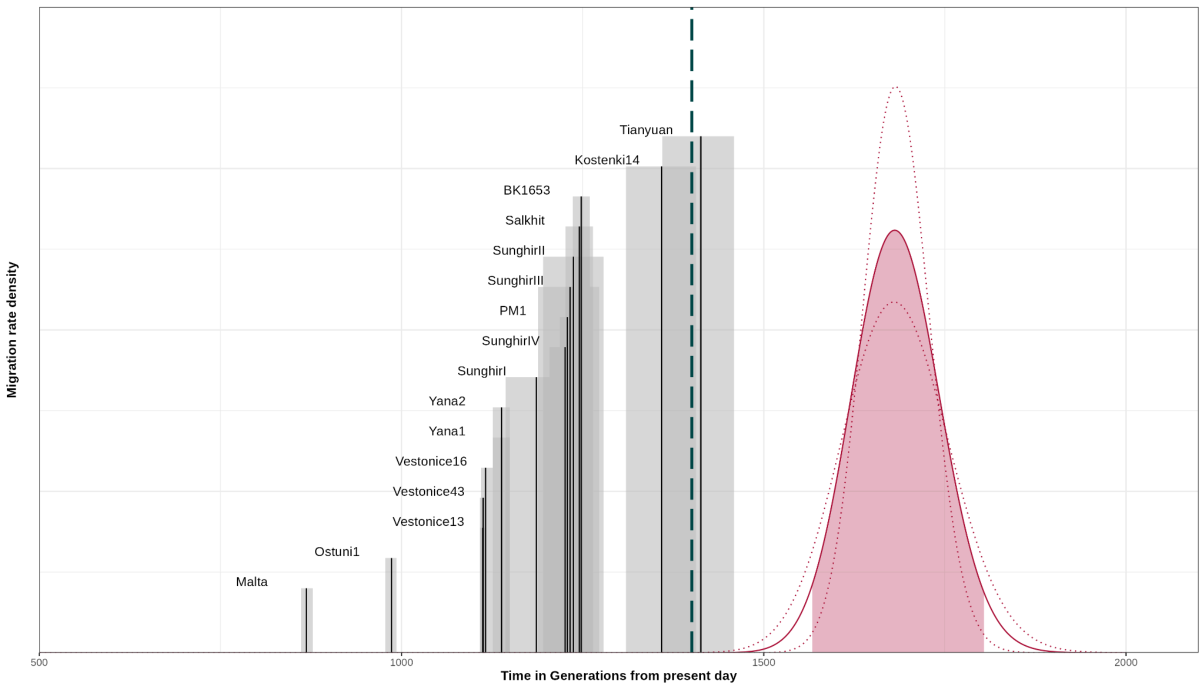

**Fig. S26.** Model prediction for the extended pulse model inferred jointly on 16 ancient individuals older than 20 kyBP. Predicted migration rate density over time with the dark red line indicates the estimate with the highest log-likelihood. Vertical line gives the estimated mean line of admixture and the shaded are the 95.5 percentile of the distribution. Dotted red lines indicating minimum and maximum estimates on the duration. Gray rectangles is the range of the most likely sample time per individual with back line giving the mean. Dashed dark green line indicates the inferred timing of Neandertal disappearance (*30*).

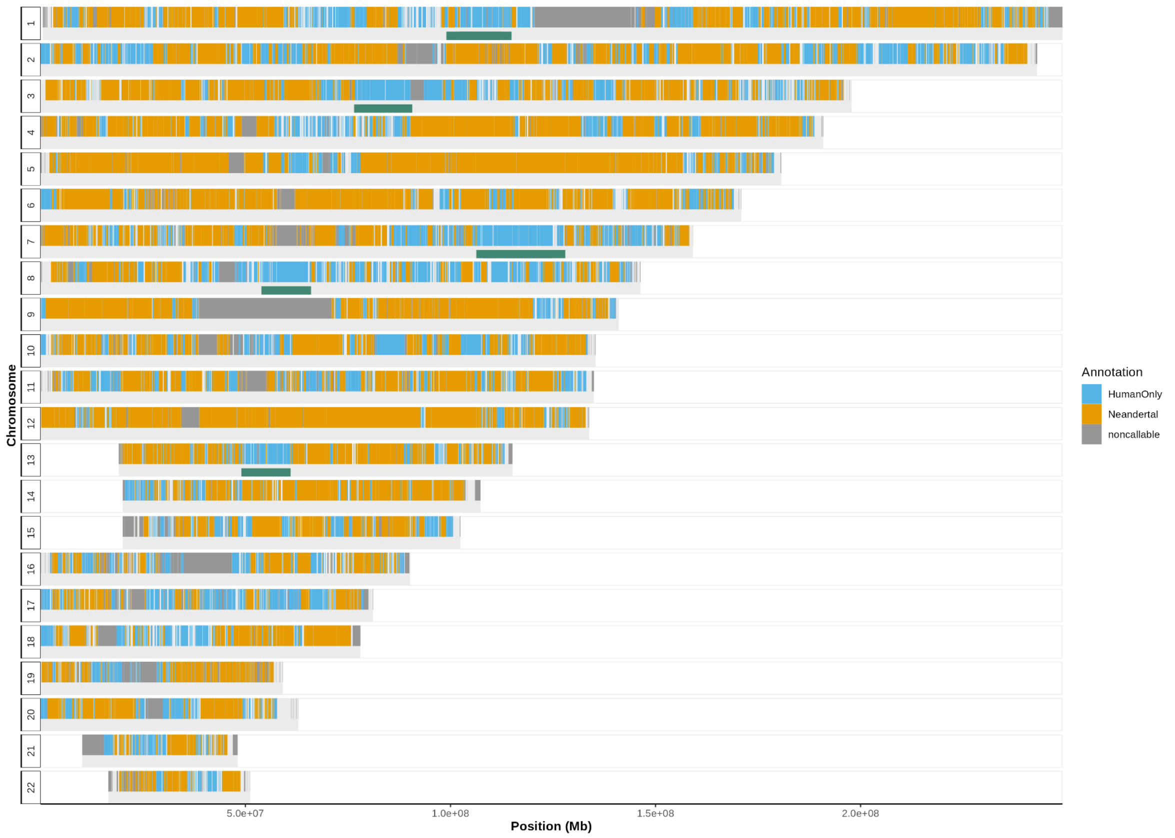

**Fig. S27.** Distribution of regions with Neandertal introgressed ancestry and genomic regions with no sign of any archaic ancestry in our data set across the human autosomal genome. Orange indicates called Neandertal ancestry in any of the individuals in our sample set. Blue indicates that all individuals in our sample set have homozygous African ancestry assigned at these positions. Non callable regions are marked in dark gray. Previously described deserts from (*13*, *15*) are indicated below the chromosomes in dark green.

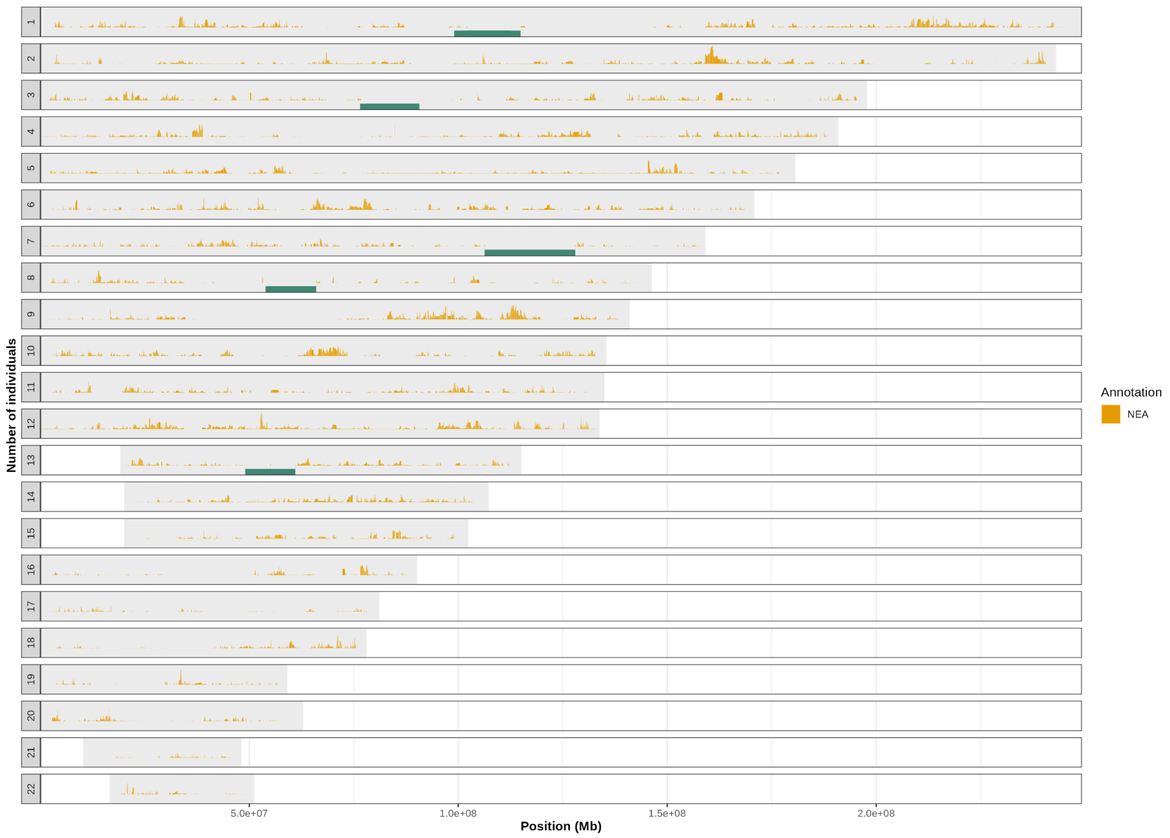

**Fig. S28.** Number of individuals having Neandertal ancestry at a given genomic location throughout the genome. Individuals include all ancient and two randomly sampled SGDP individuals per population cluster. Previously described deserts from (*13*, *15*) are indicated below the chromosomes in dark green.

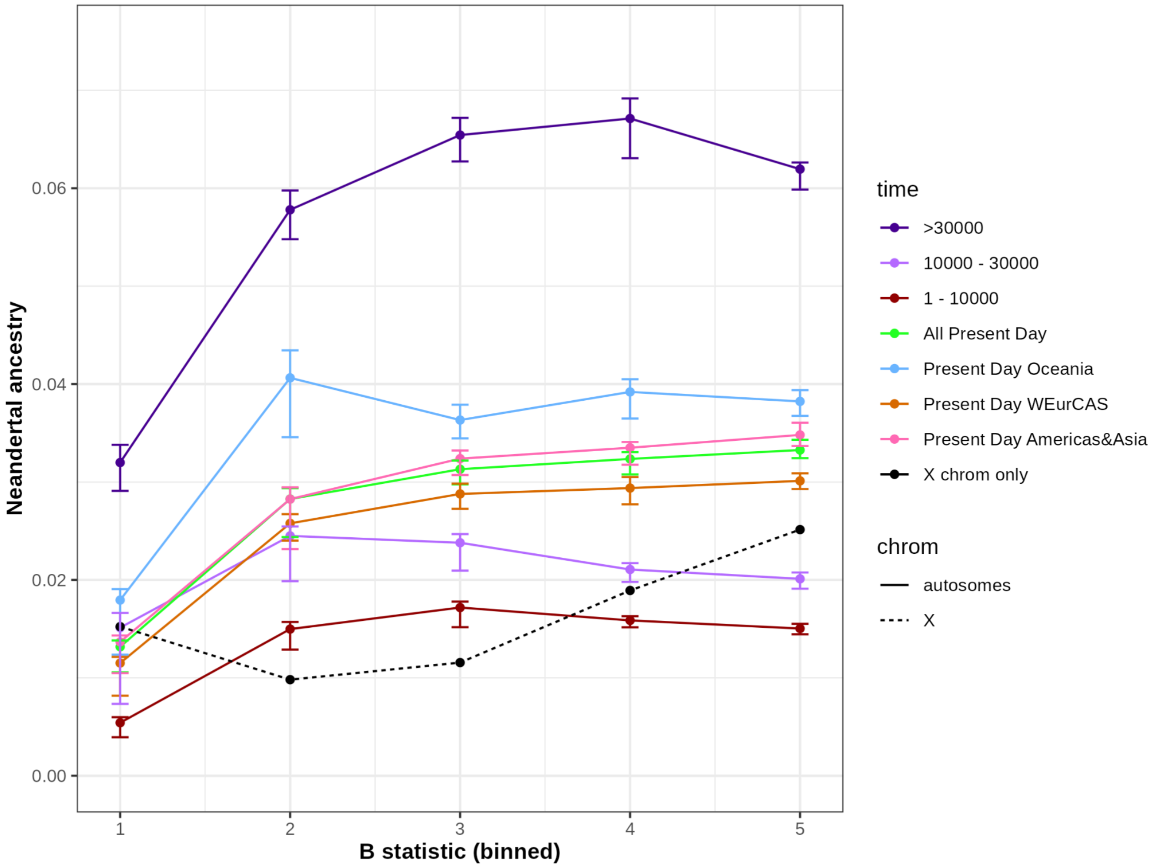

**Fig. S29.** Binned B  statistic (x-axis) versus average Archaic ancestry (y-axis) for individuals of different time intervals. Error bars are calculated by jackknife resampling where we leave out one chromosome. X chromosome estimates are without error bars. See Materials and Methods Section 7 for details on the X.

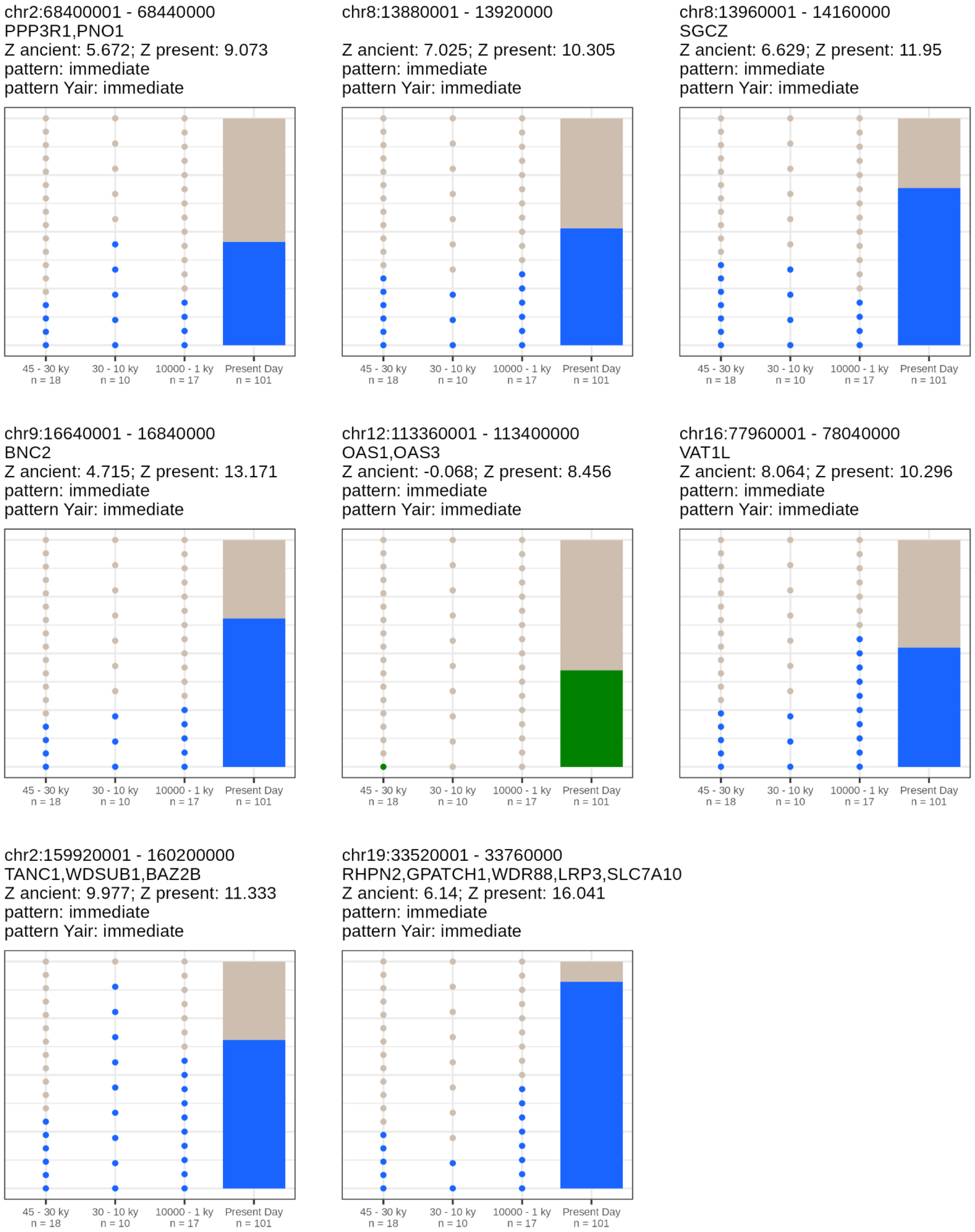

**Fig. S30.** Trajectory of Neandertal ancestry over four time intervals for regions identified in our data set that overlap previously identified regions. Number of dots corresponds to the number of individuals per time interval. Blue indicates regions identified to be constantly high in our data set. Green indicates regions identified to be high only in present day individuals. Annotation gives the genetic coordinates in hg19 for the region and genes overlapping that region.

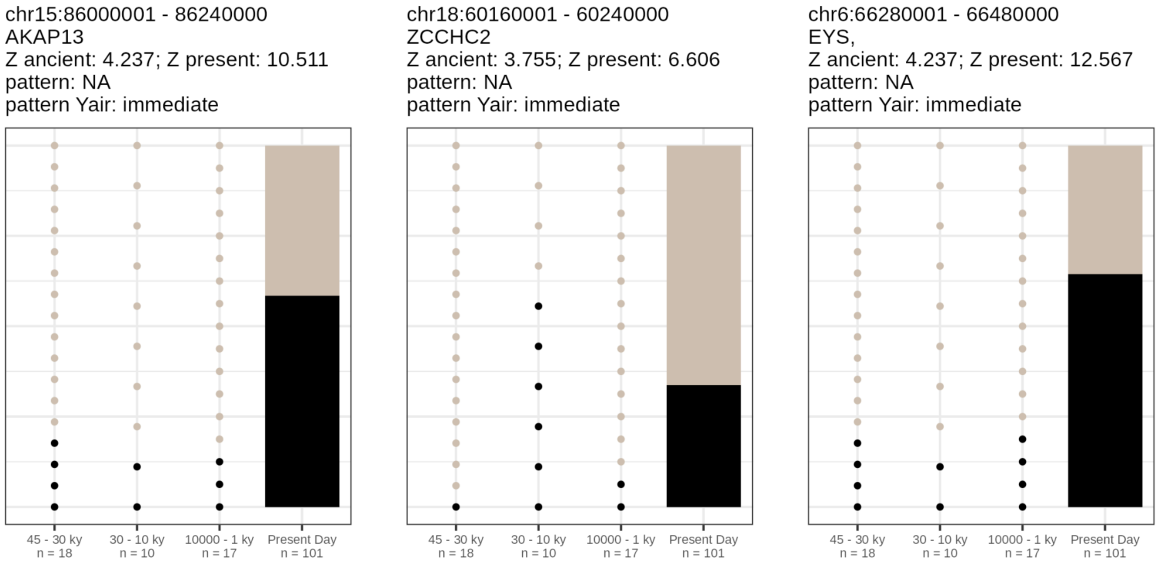

**Fig. S31.** Trajectory of Neandertal ancestry over three time intervals for regions previously identified that are not significant outliers in our data set. Number of dots corresponds to the number of individuals per time interval. Annotation gives the genetic coordinates in hg19 for the region and genes overlapping that region.

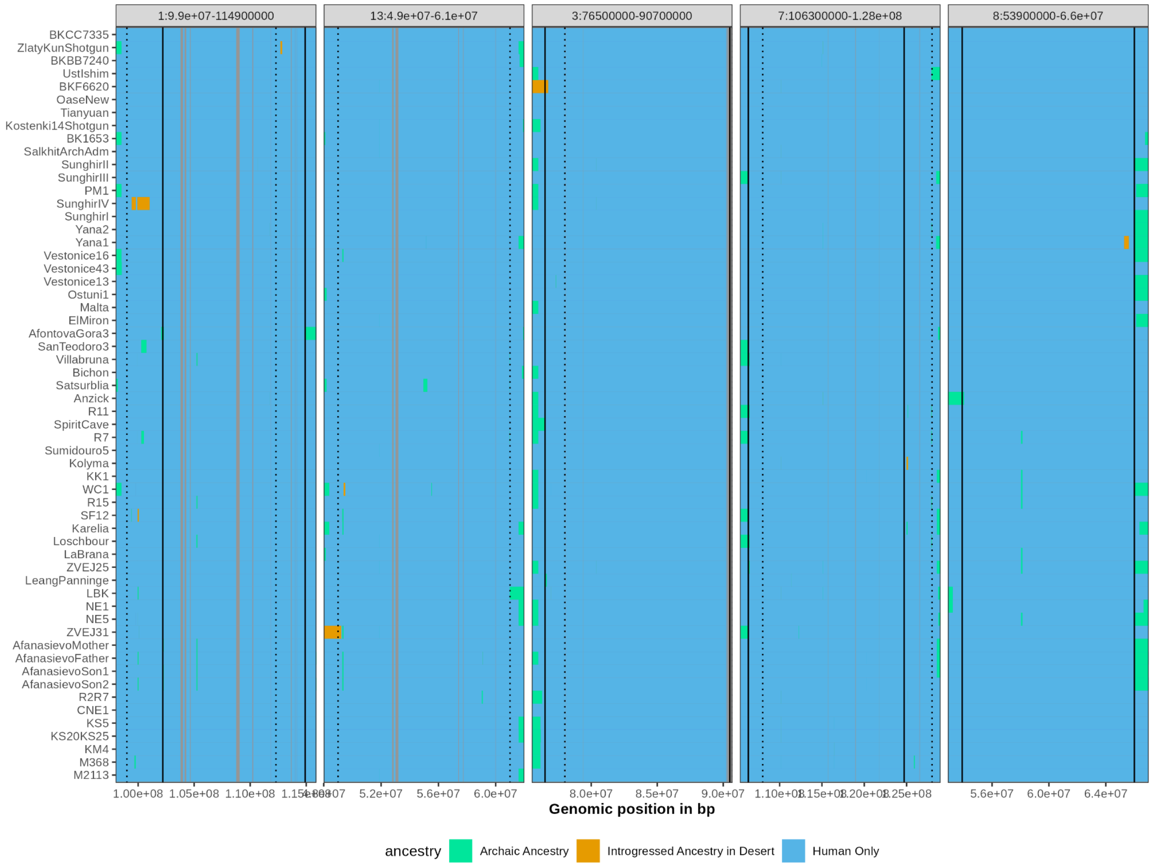

**Fig. S32.** Previously defined regions of depleted archaic ancestry from (*15*) (dotted line indicating start and end) and (*13*) (solid line) with the ancestry of ancient individuals sorted by age colored either blue if the ancestry is assigned homozygous African (Human Only), short Archaic ancestry segments in turquoise and putatively introgressed Archaic ancestry inside the deserts in orange. Non callable regions are marked in gray. Ancestry is called introgressed ancestry if the segment is 0.1 cM or 150 kb long with reads overlapping at least 5 uniquely derived sites of a single archaic reference (see fig. S33).

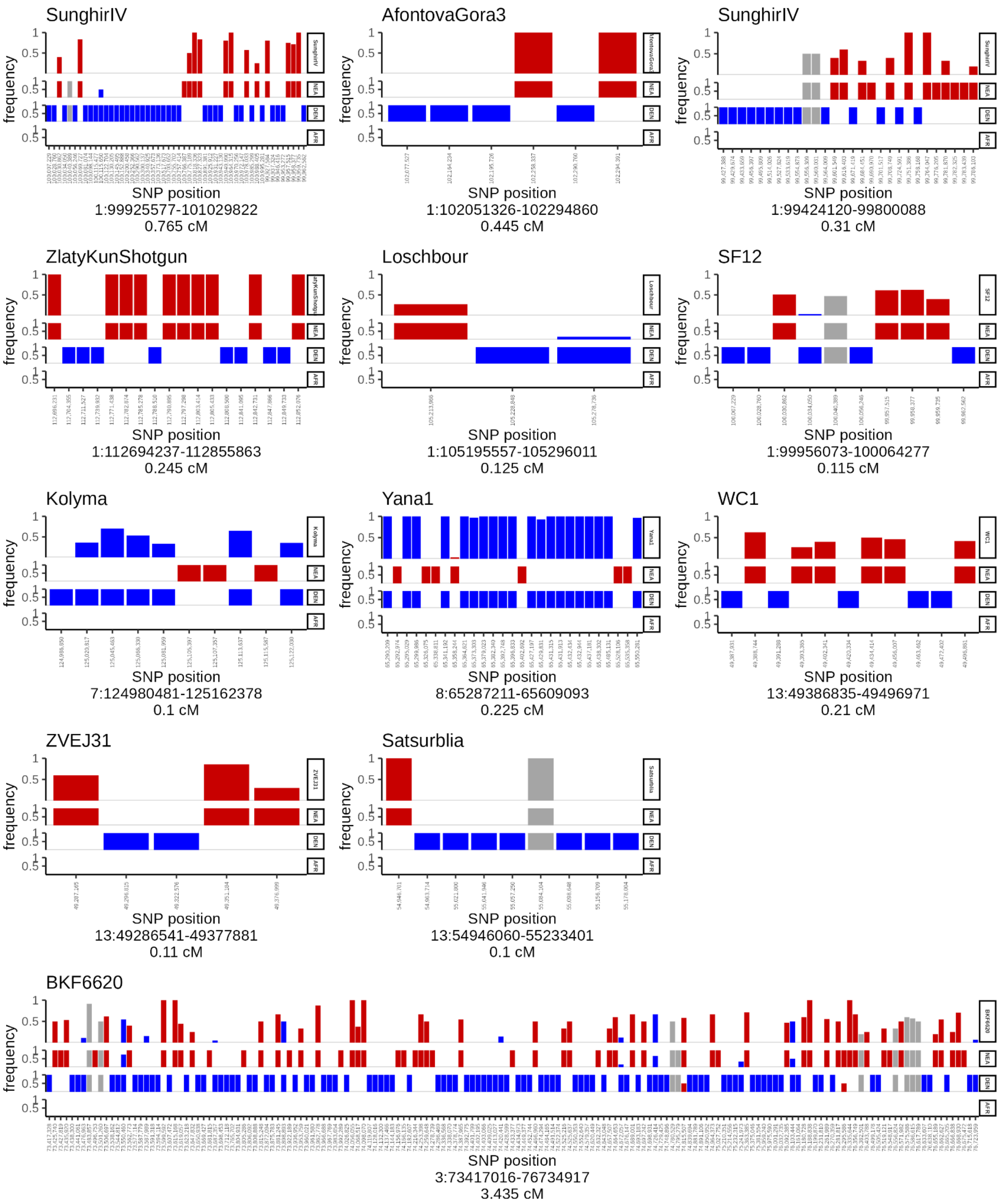

**Fig. S33.** Putatively introgressed segments in the union of deserts identified by Vernot et al. 2016 and Sankararaman et at. 2016. Each subplot represents an introgressed segment of an individual found in a desert with its position (chrom: start-end) and length in cenitMorgan (cM) annotated below the plot. Each segment is depicted as the frequency of reads per SNPs on the segment overlapping the derived allele in an individual (bars in upper facet). The position of the SNP is given on the x axis. The derived allele frequency of the reference populations Neandertal (red), Denisovans (blue) and African (gray) are indicated below the individuals facet. The bars in the individual's facet are colored red if they uniquely match with the Neandertal or blue if they uniquely match the Denisovan reference. If the SNP is not diagnostic, i.e. found to be derived in two or more references, it is colored light gray.

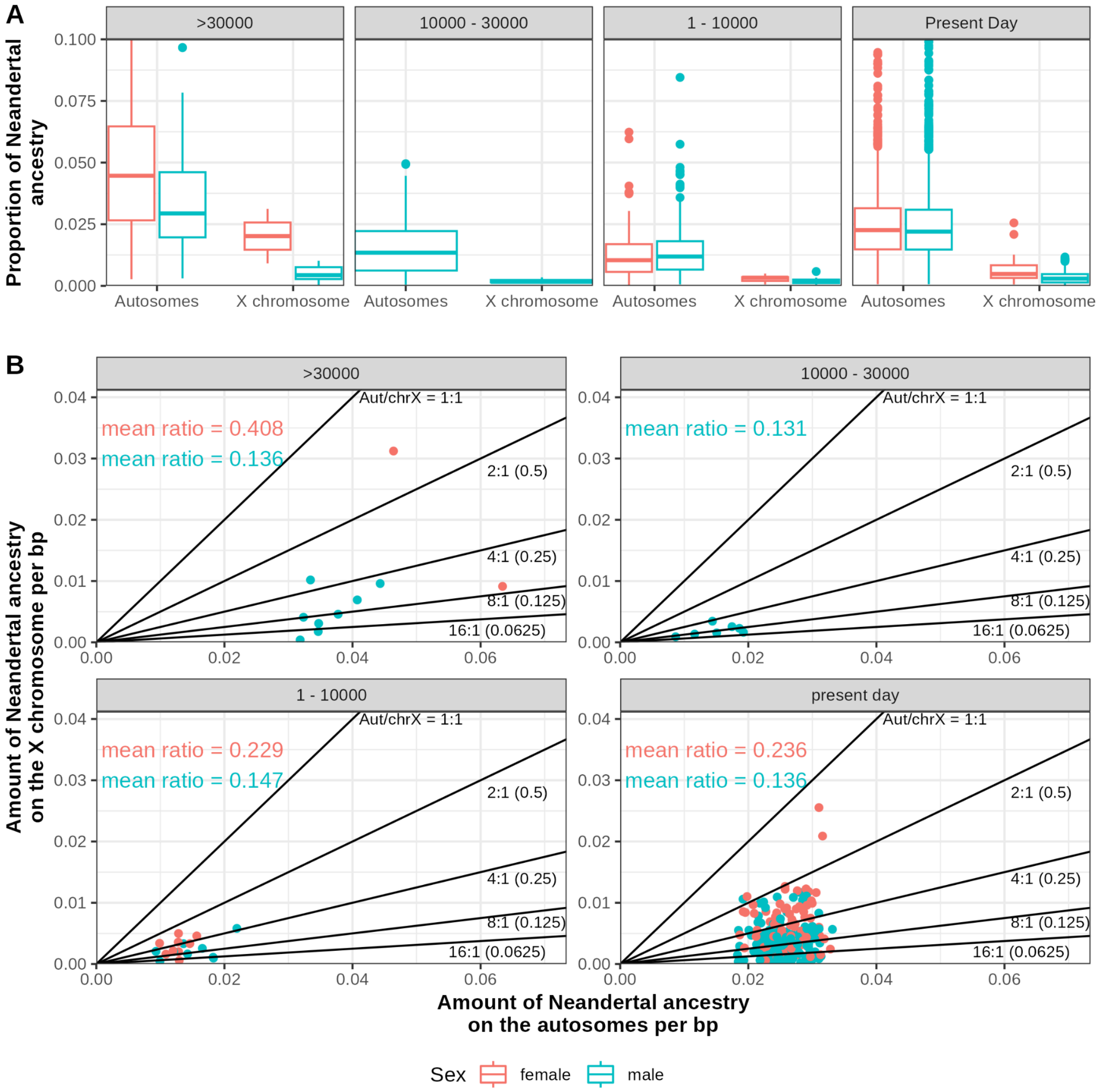

**Fig. S34.** Comparison of Neandertal ancestry on the autosomes vs. the X chromosome. (**A**) Boxplot of the proportion of Neandertal ancestry for individuals older than 30 ky, between 30 and 10 kyBP, younger than 10 ky and present-day individuals for X and autosomes for male and females. (**B**) Amount of Neandertal ancestry per base pair of callable genome for the X chromosome on the y-axis and autosome on the x-axis. Red dots indicate female individuals and blue dots male individuals. Lines indicate autosome to X ratios of 1, 0.5, 0.25, 0.125 and 0.0625.

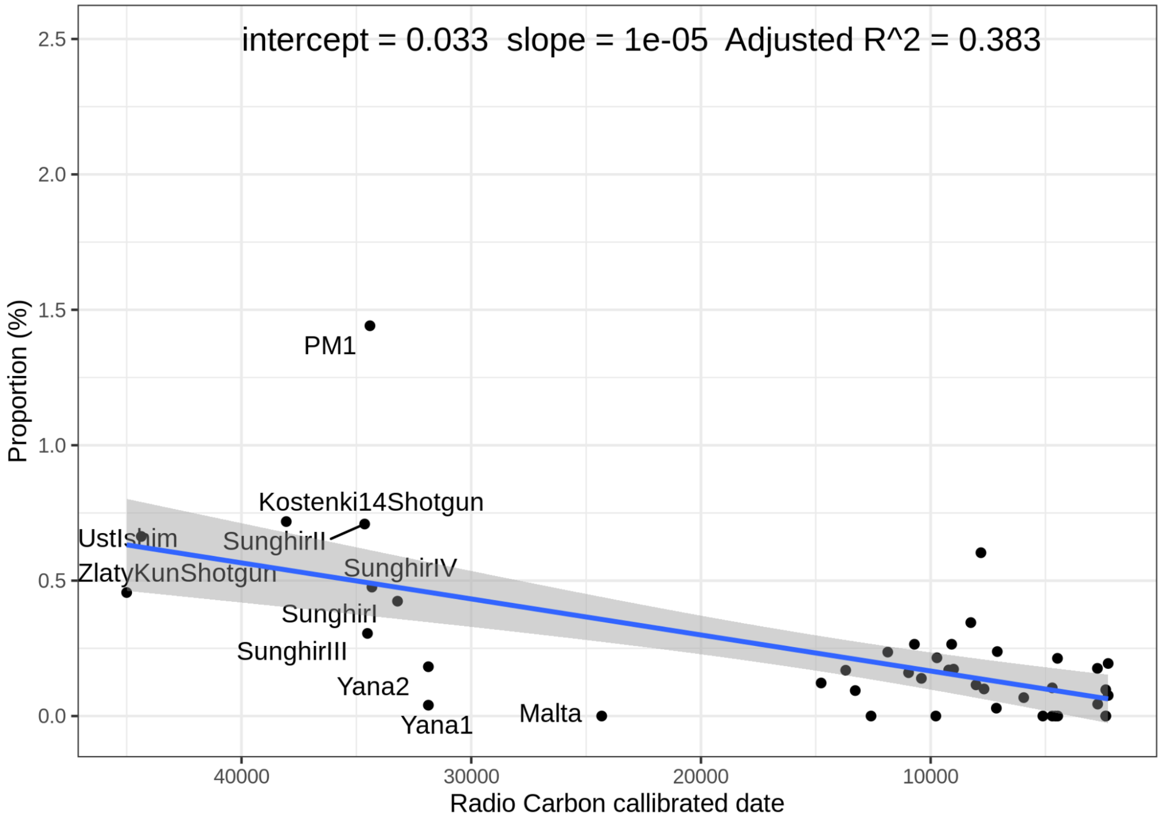

**Fig. S35.** Proportion of archaic ancestry on the X chromosome as a function of calibrated radiocarbon date. We show the fitted regression line with 95% confidence intervals and the estimated intercept, slope and coefficient of determination r2.

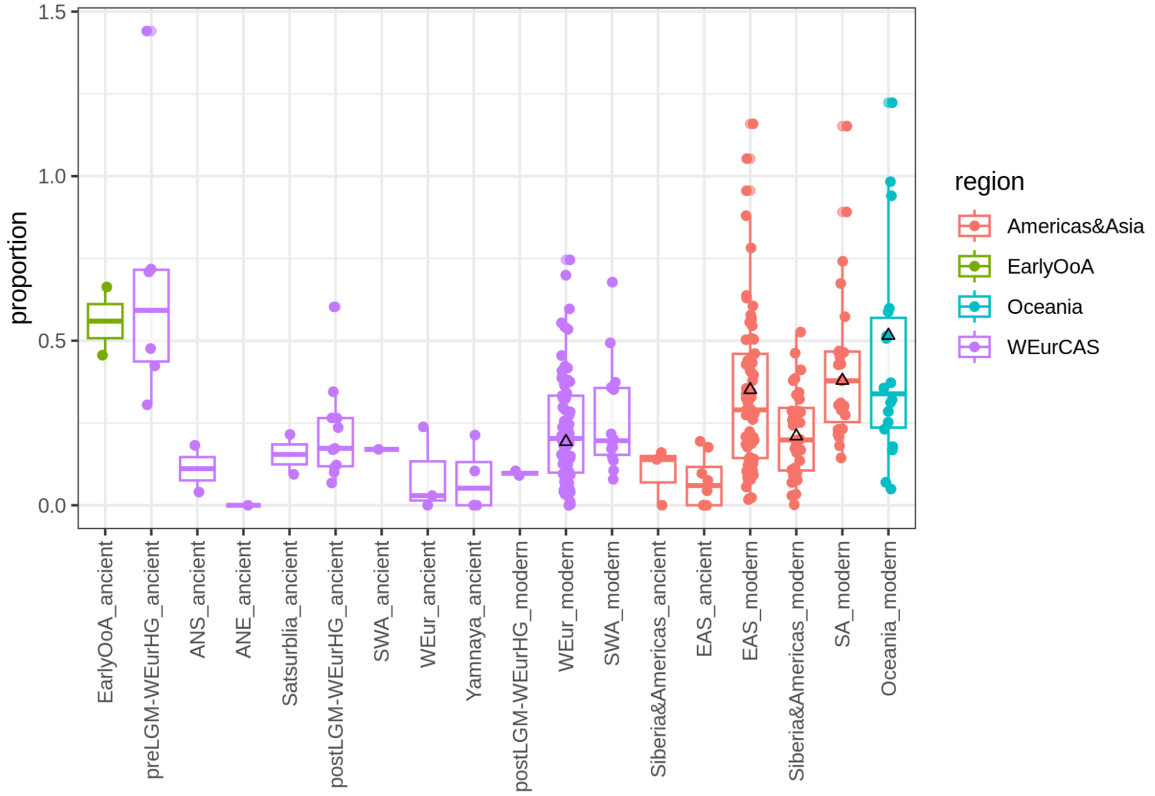

**Fig. S36.** Proportion of archaic ancestry colored by each region and sorted by mean calibrated radiocarbon date. Triangles indicate estimates of mean Neandertal proportion from Skov et al 2023.

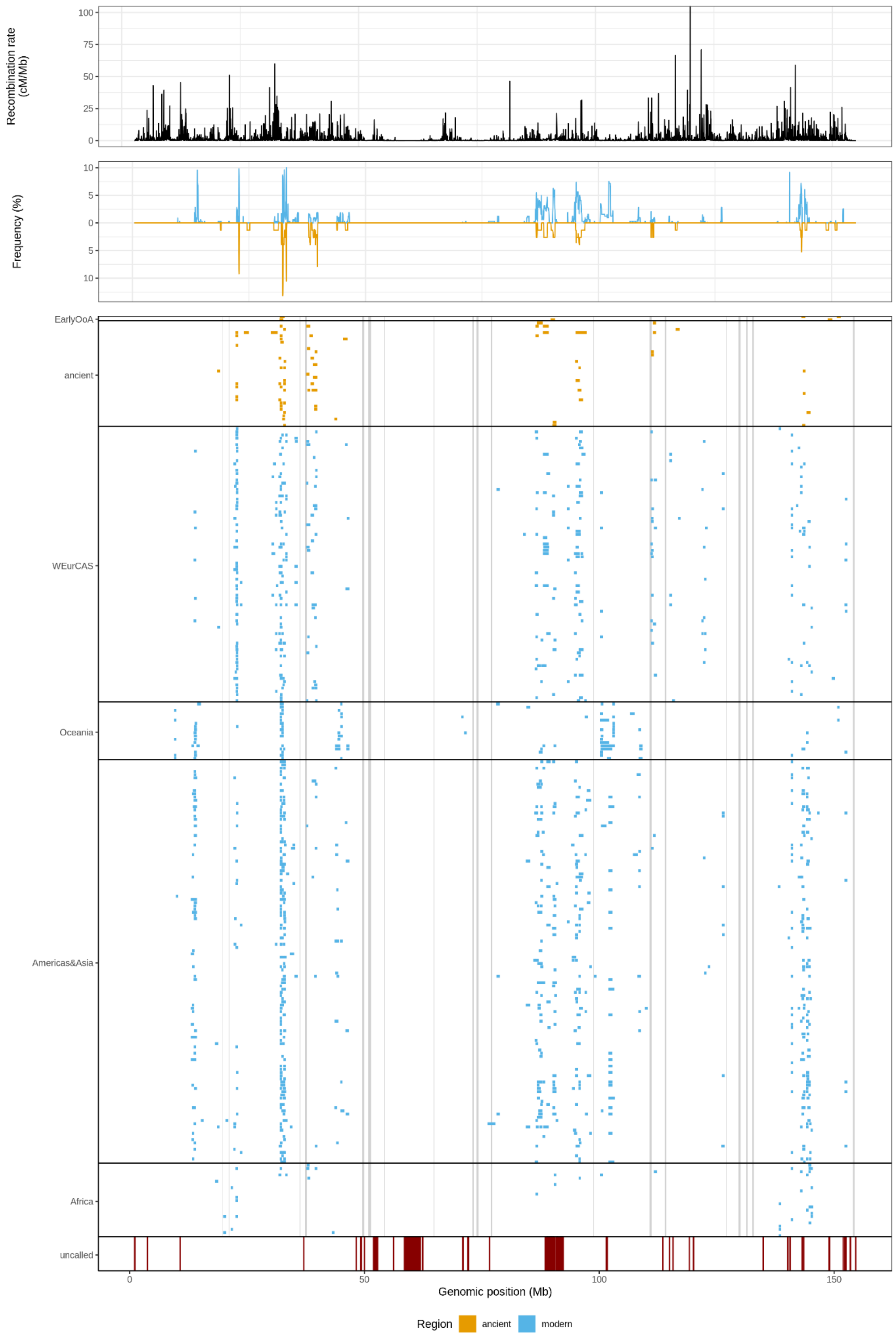

**Fig. S37.** We show the average recombination rate in 20kb windows using the African American map and truncate the y axis at 100 cM/Mb. We show the frequency of introgressed Neandertal segments for each 20kb window. We show the introgressed Neandertal segments in orange for ancient individuals and blue for modern individuals grouped by region. We show the callable regions in the genome - red boxes indicate the 10.2 Mb of the X chromosome where the SNP array contains no SNPs. Gray vertical lines show the location of sweeps (*91*).

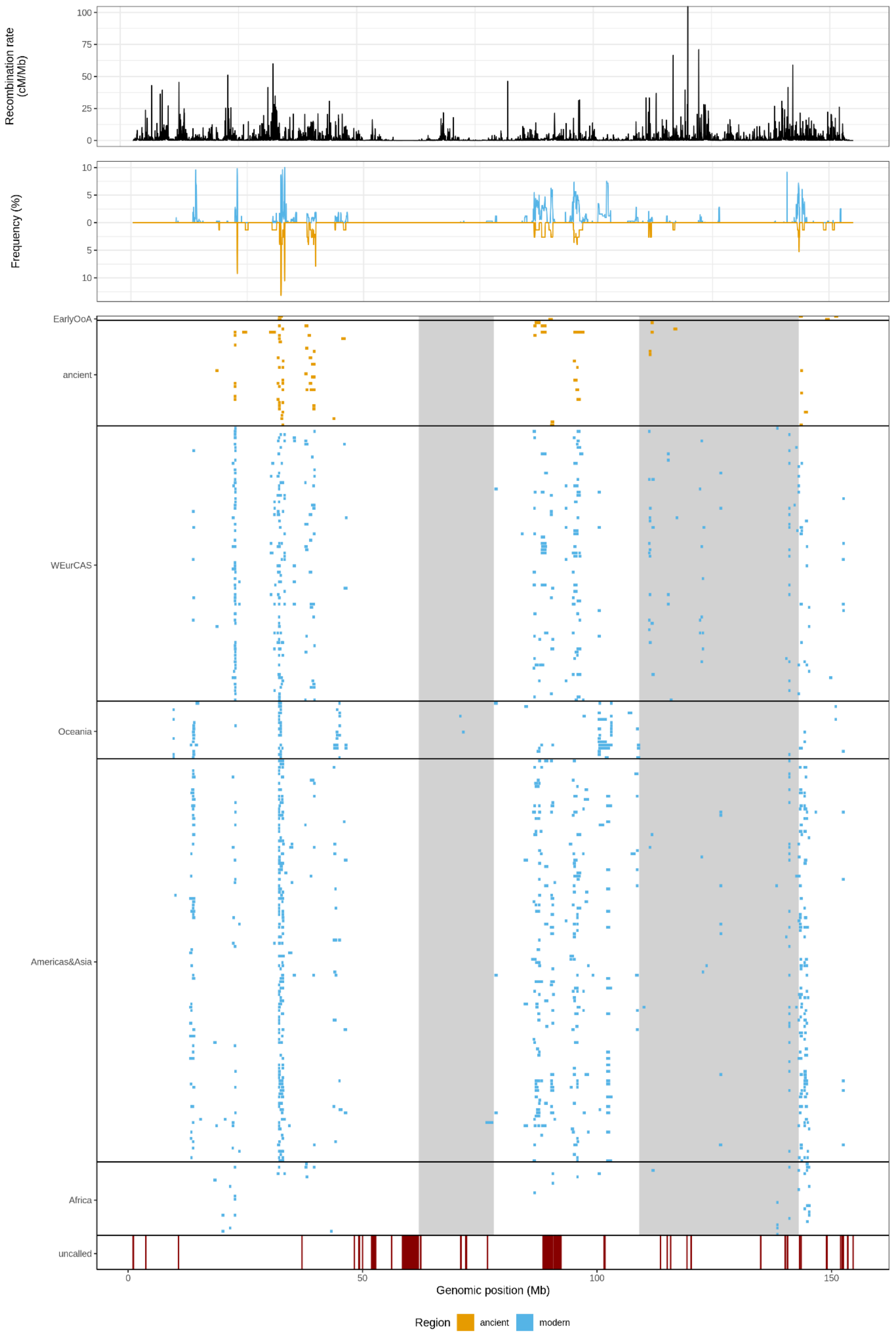

**Fig. S38.** We show the average recombination rate in 20kb windows using the African American map and truncate the y axis at 100 cM/Mb. We show the frequency of introgressed Neandertal segments for each 20kb window. We show the introgressed Neandertal segments in orange for ancient individuals and blue for modern individuals grouped by region. We show the callable regions in the genome - red boxes indicate the 10.2 Mb of the X chromosome where the SNP array contains no SNPs. Gray vertical lines show the location of deserts (*13*).

#### Supplementary Tables

| Type | Ancestral count | Ancestral % of all sites | Derived count | Derived % of all sites | Ascertainment |
|---|---|---|---|---|---|
| Shared between archaics | 38,997 | 2.78% | 144,409 | 10.28% | Archaic Admixture + ArchaicX |
| Neandertal only | 28,450 | 2.03% | 499,787 | 35.59% | Archaic Admixture + ArchaicX |
| Denisovan only | 40,597 | 2.89% | 524,999 | 37.38% | Archaic Admixture + ArchaicX |
| Shared between archaics | 12,681 | 1.16% | 8,829 | 0.80% | 1240k |
| Neandertal only | 2245 | 0.20% | 13,007 | 1.19% | 1240k |
| Deniosvan only | 8,228 | 0.75% | 6,040 | 0.55% | 1240k |
| Shared between archaics | 60,252 | 2.25% | 162,291 | 6.07% | AllDiagnosticSites |
| Neandertal only | 33,851 | 1.27% | 630,083 | 23.56% | AllDiagnosticSites |
| Denisovan only | 48,333 | 1.81% | 461,136 | 17.24% | AllDiagnosticSites |

**Table S3**. Properties of the reference panels with different ascertainments used for testing or analysis. For sites where either Neandertal and/or Denisovan references have an alternative allele not seen in the sub-Saharan African reference individuals. Or the Neandertal and/or Denisovan references have a high frequency alternative allele (>=50%) and the sub-Saharan African reference is at low frequency (<10%) or the African reference is nearly fixed (>90%) with no alternative allele seen in the Neandertal/Denisovan reference.

| Data | Slope | Intercept |
|---|---|---|
| All 22 individuals | 28.29 ± 0.61 | 46334.45 ± 257.05 |
| 16 individuals (Oase, BKBB7240, BKCC7335, BKF6620 and Ust'-Ishim, Zlatý kůň excluded) | 28.40 ± 1.42 | 46363.72 ± 681.4 |

**Table S18.** Slope (generation time) and intercept (Neandertal admixture time) estimates and standard error of the Bayesian model.

| IG tm | IG LL | EP MLL tm | EP MLL td | EP LL | LR | p_value | truncation | weighted | Data |
|---|---|---|---|---|---|---|---|---|---|
| 1653 | 66306 | 1672 | 1 | 66334 | 56 | 6.21E-14 | estimated truncation | Same weight for each Individual | All Data |
| 1659 | 57813 | 1675 | 1 | 57828 | 33 | 6.70E-09 | estimated truncation | Same weight for each Individual | Oase Excluded |
| 1657 | 15542 | 1672 | 1 | 15538 | -7 | 1.00E+00 | estimated truncation | Same weight for each Individual | Only EarlyOoA |
| 1612 | 50275 | 1683 | 244 | 51041 | 1533 | < 1E-22 | estimated truncation | Same weight for each Individual | EarlyOoA Excluded |
| 1666 | 6947 | 1675 | 26 | 6938 | -16 | 1.00E+00 | estimated truncation | Same weight for each Individual | Only EarlyOoA w/o Oase |

**Table S22**. Comparison of the IG and EP model. With the estimated values for tm and td (for EP model only), corresponding log-likelihood, likelihood-ratios between IG and EP and corresponding p-value for comparison using different parameters for truncation, weight and data used.

| time interval | percentile empirical distribution | n windows | n windows lower than any desert | average proportion of archaic ancestry in published deserts |
|---|---|---|---|---|
| Present-day | 0.008 | 516 | 4 | 0.0016 |
| 1 - 10000 | 0.025 | 516 | 13 | 0.0018 |
| 10000 - 30000 | 0.058 | 516 | 30 | 0.0020 |
| >30000 | 0.031 | 516 | 16 | 0.0017 |

**Table 30**. Percentile of published deserts with the highest amount of archaic ancestry from an empirical distribution of archaic ancestry in windows across the genome for different time intervals.

| B score bin | range B score | Neandertal frequency |
|---|---|---|
| 1 | 0-199 | 0.015 |
| 2 | 200-399 | 0.01 |
| 3 | 400-599 | 0.012 |
| 4 | 600-799 | 0.019 |
| 5 | 800-10000 | 0.025 |

**Table 32.** Amount of Neandertal ancestry in regions with different B scores, where low B scores are conserved regions.

| minimum segment length | window_size | Spearman's rank test correlation coefficient | p_value |
|---|---|---|---|
| 0.05 / 0.2 | 20000 | 0.051 | 0.044 |
| 0.05 / 0.2 | 50000 | -0.015 | 0.704 |
| 0.05 / 0.2 | 100000 | -0.047 | 0.361 |
| 0.05 / 0.2 | 200000 | -0.023 | 0.730 |
| 0.05 / 0.2 | 500000 | -0.017 | 0.851 |
| 0.05 / 0.2 | 1000000 | 0.049 | 0.671 |
| 0.01 | 20000 | -0.002 | 0.925 |
| 0.01 | 50000 | -0.055 | 0.102 |
| 0.01 | 100000 | -0.091 | 0.036 |
| 0.01 | 200000 | -0.063 | 0.251 |
| 0.01 | 500000 | 0.020 | 0.784 |
| 0.01 | 1000000 | 0.078 | 0.414 |

**Table 33.** Correlation of the amount of Neandertal ancestry with local recombination rate of the African-American Map for different window sizes and minimum segment length cutoffs.

| sweep region | start | end | Introgressed haplotypes | Introgressed haplotype present in |
|---|---|---|---|---|
| 1 | 19800000 | 19900000 | 0 | - |
| 2 | 21200000 | 21300000 | 0 | - |
| 3 | 36200000 | 36400000 | 0 | - |
| 4 | 37300000 | 37700000 | 0 | - |
| 5 | 49500000 | 50000000 | 0 | - |
| 6 | 50800000 | 51400000 | 0 | - |
| 7 | 54200000 | 54400000 | 0 | - |
| 8 | 64700000 | 64900000 | 0 | - |
| 9 | 73000000 | 73300000 | 0 | - |
| 10 | 73900000 | 74300000 | 0 | - |
| 11 | 76900000 | 77200000 | 1/493 | S_Brahmin-1 |
| 12 | 98700000 | 98800000 | 0 | - |
| 13 | 110700000 | 111100000 | 0 | - |
| 14 | 114000000 | 114300000 | 0 | - |
| 15 | 127000000 | 127100000 | 0 | - |
| 16 | 129700000 | 130100000 | 0 | - |
| 17 | 131300000 | 131600000 | 0 | - |
| 18 | 132600000 | 132900000 | 0 | - |
| 19 | 154000000 | 154400000 | 0 | - |

**Table 31**. Location of sweeps with the frequency of the sweep is maximal on the X chromosome in hg19 coordinates. We show the number of introgressed haplotypes and which individuals carry them.
